# Adhesome Receptor Clustering is Accompanied by the Co- localization of the Associated Genes in the Cell Nucleus

**DOI:** 10.1101/2023.12.07.570697

**Authors:** Louis V. Cammarata, Caroline Uhler, G. V. Shivashankar

## Abstract

Proteins on the cell membrane cluster to respond to extracellular signals; for example, adhesion proteins cluster to enhance extracellular matrix sensing; or T-cell receptors cluster to enhance antigen sensing. Importantly, the maturation of such receptor clusters requires transcriptional control to adapt and reinforce the extracellular signal sensing. However, it has been unclear how such efficient clustering mechanisms are encoded at the level of the genes that code for these receptor proteins. Using the adhesome as an example, we show that genes that code for adhesome receptor proteins are spatially co-localized and co-regulated within the cell nucleus. Towards this, we use Hi-C maps combined with RNA-seq data of adherent cells to map the correspondence between adhesome receptor proteins and their associated genes. Interestingly, we find that the transcription factors that regulate these genes are also co-localized with the adhesome gene loci, thereby potentially facilitating a transcriptional reinforcement of the extracellular matrix sensing machinery. Collectively, our results highlight an important layer of transcriptional control of cellular signal sensing.

## Introduction

Cells must respond to various physical and biochemical signals in our tissues ^1^. To achieve this, the plasma membrane of cells comprises several sensors including receptors, ion channels, and others ^2^. Using such sensors, cells can rapidly adapt to varying signals in the microenvironment via, e.g., cell matrix adhesion through integrin receptors ^3^, cell-cell interactions through E-cadherin receptors ^4^, or antigen binding through T-cell receptors ^5^. A common underlying mechanism in such signal sensing and adaptation requires the rapid clustering of the receptors and various protein adaptors to reinforce signal sensing and cellular response ^6^. While some of these receptors already exist on the plasma membrane, signal sensing and maturation require the production of additional proteins involved in these sensing mechanisms and thus require the transcriptional regulation of their respective genes ^7^. To allow for rapid sensing and adaptation, we hypothesized that there may exist a map between the proteins that cluster on the cell membrane and the genes that code for these proteins within the nucleus. More specifically, we hypothesized that the genes that code for the proteins as well as the transcription factors required for their regulation are spatially clustered within the nucleus to allow for optimal control and reinforcement of the sensing mechanisms of a cell; a schematic of our hypothesis is shown in Fig. 1.

**Fig. 1:**
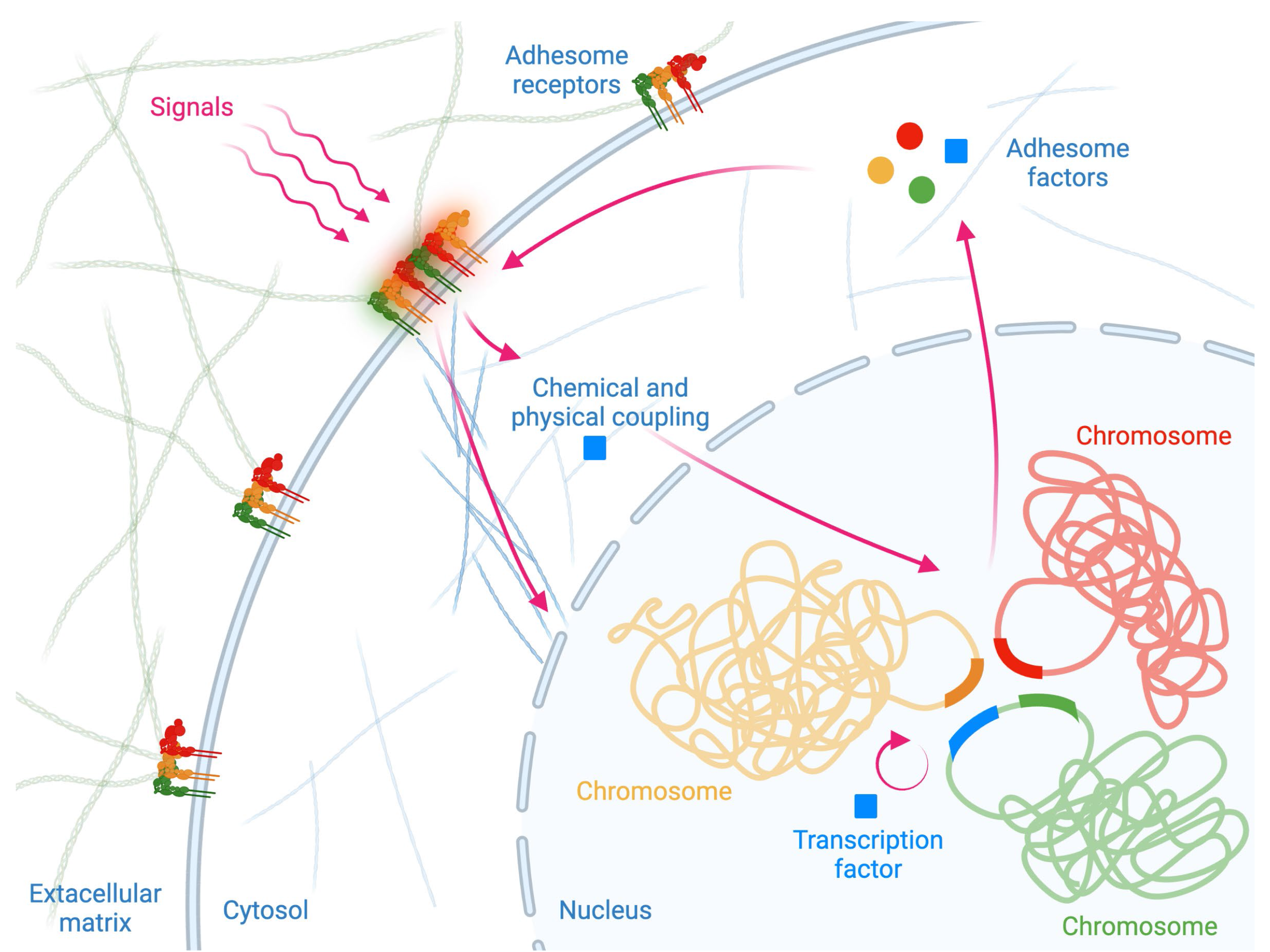
Graphical abstract. Schematic outlining our proposed hypothesis, i.e., that there exists a map between the receptor proteins that cluster on the cell membrane and the genes that code for these proteins within the cell nucleus. More specifically, we hypothesize that the genes that code for the proteins (e.g., integrin receptors) as well as the transcription factors required for their regulation are spatially clustered within the nucleus to allow for optimal control and reinforcement of the sensing mechanisms of a cell. Schematic created with BioRender.

Recent progress in genomic and proteomic technologies have allowed the characterization of various proteins involved in signal sensing and transduction ^8–11^ as well as measuring the spatial organization of genomes ^12–15^. For example, many of the proteins involved in cell matrix sensing to assemble focal adhesions have been annotated. In addition, recent technological developments in genomics coupled with efforts in bioinformatics have resulted in the annotation of many of the transcription factors that participate in regulating adhesome genes ^16–18^. Furthermore, recent progress in chromosome conformation capture assays have allowed mapping the contact frequencies of the 3D organization of genomes in multiple cell types ^13,19,20^. Additionally, fluorescence in situ hybridization (FISH) technologies have enabled the identification of the spatial location of particular genes and chromosomes in single cells ^21,22^, thereby validating the 3D organization of genomes obtained using chromosome conformation capture methods. Importantly, advances in sequencing methods have allowed to obtain not only high resolution maps of the spatial organization of the genome, but also the transcriptional state of the whole genome ^23,24^. Given the vast datasets that exist in the context of focal adhesion, we decided to study this critical cell matrix sensing process and test our hypothesis of whether the adhesome genes are spatially co-clustered and co-regulated in the cell nucleus of adherent fibroblast cells.

Chromosomal conformation capture technologies provided early evidence for the spatial clustering of functionally related genes (located on different chromosomes). A data-driven study of the human lymphoblastoid cell line GM06990 revealed that genes belonging to the same functional group (e.g., genes that code for interacting proteins, participate in the same complex, or pathway) tend to colocalize in space ^25^. More recently, interchromosomal genomic clusters that are both spatially colocalized and coregulated were identified in the IMR-90 cell line ^26^. These findings align with the condensate and transcription factory model according to which actively transcribed genes across multiple chromosomes cluster around sites of highly concentrated transcription machinery ^27^, possibly being held together through non-coding RNA ^28^ and transcription factor binding ^29^. They are also consistent with experimental studies of the past two decades describing the spatial clustering of olfactory receptor genes, globin genes, immunoglobulin genes, Hox genes, heat shock protein genes, and Plasmodium virulence genes, among other functional gene groups ^30^. Interchromosomal spatial clustering could be an efficient way to reinforce the co-regulation of these gene groups in a cell type-specific manner ^31,32^, as shown for KLF1-regulated genes ^33^ and NFκB- responsive genes ^34^.

In this article, we show the existence of a map between adhesome proteins that cluster on the cell membrane and the genes that code for these proteins and their regulators in the nucleus of IMR- 90 cells. Using transcriptomic data, we first confirm that most adhesome genes are active in adherent cells. Using chromosome conformation capture data, we then show that the active adhesome genes are significantly more co-localized in the cell nucleus as compared to other active non-adhesome genes. Interestingly, the adhesome genes also show strong transcriptional co-regulation. We then leverage transcription factor binding data together with chromosome conformation capture data to show that the transcriptional regulators of the adhesome genes are also co-localized with these genes, thereby facilitating rapid reinforcement of the extracellular matrix sensing. These different data modalities enable identifying tight clusters of adhesome genes and their transcriptional regulators, which we further validate using publicly available genome-wide FISH data. This study reveals a novel fundamental layer of transcriptional control to facilitate rapid cellular signal sensing in complex physiological environments.

## Results

### Genes coding for focal adhesion proteins that are clustered on the plasma membrane are co-localized in the cell nucleus

Cells use their receptors on the plasma membrane to rapidly sense their external microenvironment; for example, different focal adhesion proteins have to cluster on the plasma membrane and work together in order to adhere to the extracellular matrix and activate downstream cellular processes. We hypothesized that such rapid clustering of different proteins may require the genes that code for these proteins to be spatially clustered in the nucleus for optimal control of their co-regulation. Similarly, for the reinforcement of focal adhesions, a host of additional proteins are required, and we hypothesized that the genes coding for these proteins are also spatially co-clustered for their efficient co-regulation and for the production of these proteins. Evidence of an interplay between transcription and genome organization upon signal sensing was reported in previous studies including in fibroblasts ^15^, for heat shock proteins ^35^, and upon TNFα stimulation ^34^.

Towards this, we thought to analyze the proximity of adhesome genes in an adherent cell type. We used IMR-90, a human female fetal lung fibroblast cell line as a model system for this study. As a first step, we obtained a curated list of 232 proteins that reside directly or indirectly in adhesion sites from different cell types ^36^. These proteins include integrins, actin regulators, adaptor proteins that link the cytoskeletal structures to the cytoplasmic tails of integrins, as well as a variety of signaling molecules (*SI Appendix*, Supplementary Fig. 1). Gene set enrichment analysis confirmed the key role of these genes in regulating cell-matrix adhesion, migration, and cytoskeletal organization, among others (*SI Appendix*, Supplementary Fig. 2A). A number of studies have shown that adhesome proteins are densely linked via binding or signaling interactions, suggesting that these proteins have to work together in the adhesome complex ^37^ (*SI Appendix*, Supplementary Fig. 2B). In our work, we focused on adhesome genes located on the autosomes 1 to 22, thus leaving out from the analysis five adhesome genes located on chromosome X (ARHGEF6, FLNA, MSN, SH3KBP1 and SMPX).

Next, we analyzed expression data of adherent cells to verify that the adhesome genes are actively being transcribed. For this, we retrieved from previous studies bulk RNA-seq data for IMR-90 ^38^, as well as preprocessed cistromic data on 48 regulatory marks ^39^. The regulatory marks included chromatin accessibility data (DNase-seq) and a selection of histone and transcription factor ChIP-seq data (*SI Appendix*, Supplementary Fig. 3A). This data was collectively used to determine the activity status of genes in IMR-90. In particular, we identified regulatory signals corresponding to 25,959 genes, whose genomic locations were obtained from a previous study ^40^. We represented each gene by a 48-dimensional feature vector of regulatory marks, where each feature was centered by its mean and scaled by its standard deviation across all genes. In this representation, the affinity between two genes was measured by the cosine similarity between their feature vectors. We applied hierarchical clustering with average linkage to this data to identify two clusters of genes, cluster 1 containing 14,122 genes and cluster 2 containing 11,837 genes (*SI Appendix*, Supplementary Fig. 3B and 41). These two clusters are depicted in blue and orange in the t-distributed Stochastic Neighborhood Embedding (t-SNE) plot of Fig. 2A. By comparing the activity status of each of the two gene clusters based on the distribution of RNA-seq expression, we identified cluster 1 to correspond to active genes and cluster 2 to correspond to inactive genes (Fig. 2B); the difference in expression is highly significant (Wilcoxon rank-sums test, *p*-value < 3e-308) and consistent with peaks from known active marks (e.g., POLR2A, H3K36me3) and repressive marks (e.g., H3K9me3, H3K27me3).

**Fig. 2:**
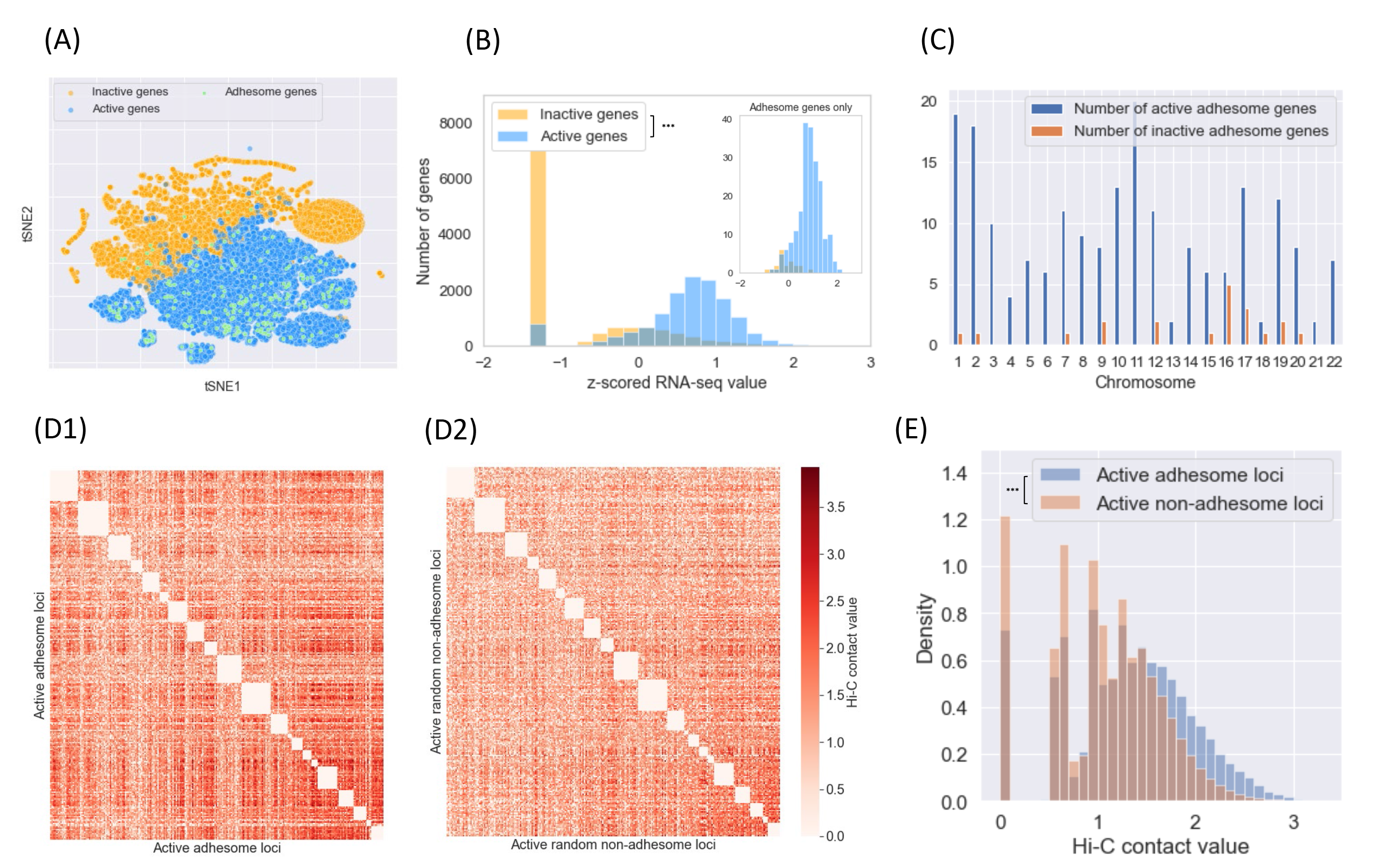
Adhesome genes are active and spatially co-localized in the nucleus of IMR-90 cells. (A) tSNE representation of 25,959 genes based on 48 IMR-90 regulatory marks. Orange dots correspond to inactive genes (as determined by the corresponding expression of regulatory marks), and blue dots correspond to active genes. Out of the 222 adhesome genes studied (green dots), 202 are active. (B) Histogram of RNA-seq expression in IMR-90 for all active genes (blue) and all inactive genes (orange). Active genes have a significantly higher expression than inactive genes (*p*-value < 3e-308, Wilcoxon rank-sums test). The embedded graph in the upper right corner corresponds to the histogram of RNA-seq expression in IMR-90 for all active adhesome genes and all inactive adhesome genes. Active adhesome genes have a significantly higher expression than inactive adhesome genes (*p*-value < 2e-9, Wilcoxon Rank-Sums test). (C) Bar plot of the number of active (blue) and inactive (orange) adhesome genes per chromosome. (D) Interchromosomal Hi-C contact matrices for active adhesome loci (D1) and a randomly selected set of active non-adhesome loci chosen on the same chromosomes (D2). For robustness, an active locus is defined to be a locus that contains at least one active gene and that was labeled as active in the locus clustering. (E) Histograms of Hi-C contact frequencies across all adhesome loci (blue) and all active non-adhesome loci (orange). Hi-C contacts are significantly higher among active adhesome loci compared to active non-adhesome loci (*p*-value < 3e-308, Wilcoxon Rank-Sums test).

Out of the 222 adhesome genes whose genomic locations on the hg19 reference genome were available in the UCSC Genome Browser ^40,42^, 202 genes were found to be active in IMR-90 cells. These active adhesome genes are represented as light green dots in the t-SNE plot of Fig. 2A. This representation was obtained using available transcriptomic data as well as data on 1D genomic marks (due to limited public data availability, noncoding RNAs, which are known to play a key role in chromatin tethering ^30,43^, could not be considered in this analysis). For comparison, conducting a similar analysis in GM12878, a human lymphoblastoid cell line, showed that only 172 adhesome genes were identified as active in this non-adherent cell line (*SI Appendix*, Supplementary Fig. 4). While, as expected, this number is lower compared to IMR-90, a good proportion of adhesome genes remain active in GM12878 as they participate in other signaling pathways in this cell type. For example, the B cell receptor (BCR) signaling pathway is mostly active in GM12878, which is a B cell lymphoblastoid (63 BCR genes are active out of 81 in our data); this pathway is related to other signaling pathways ^44^ including actin cytoskeleton organization, MAPK signaling, NF-κB signaling and calcium-mediated signaling, which contain genes that overlap with active adhesome genes in GM12878 (*SI Appendix*, Supplementary Fig. 5).

The identified adhesome genes are located on all 22 autosomes (Fig. 2C), suggesting that interchromosomal interactions may be essential for their co-regulation. To test if active adhesome genes are spatially co-clustered in the cell nucleus, we assessed the proximity of the 202 active adhesome genes in IMR-90 across chromosomes. This was carried out using publicly available genome-wide chromosomal conformation capture data assayed with the in situ Hi-C protocol ^20^. While the Hi-C data set allows resolution up to 5 kb, we chose to study interactions on a 250 kb contact map, since this resolution allows to locate genes of interest while reasonably controlling the sparsity of interchromosomal Hi-C data. As described in our earlier work ^26^, Hi-C contact frequencies were normalized genome-wide using the Knight-Ruiz matrix balancing algorithm ^45^, log-transformed and processed to filter out repeat regions, centromeric and pericentromeric regions obtained from the UCSC Genome Browser ^41^.

We compared the distribution of interchromosomal Hi-C contact frequencies among active adhesome loci (defined as loci containing active adhesome genes) and among all active non-adhesome loci (defined as loci containing active non-adhesome genes, but no active adhesome genes). Throughout our work, interchromosomal interactions refer to non-homologous chromosomal contacts (NHCCs) ^46^. Figure 2D1 and Fig. 2D2 show heatmaps of interchromosomal Hi-C contacts between all active adhesome loci as compared to interchromosomal Hi-C contacts between a set of randomly chosen active non-adhesome loci. To obtain an unbiased comparison, we used the same number of active adhesome and non-adhesome loci on each chromosome. It is apparent from these heatmaps that Hi-C contacts are higher among active adhesome loci. As expected, contacts tend to be higher for the small, gene-rich autosomes ^19^ (chromosomes 16-22, lower right corner of Fig. 2D1), whose chromosomal territories are preferentially located in the center of the nucleus, while the chromosome territories of larger chromosomes are usually found at the nuclear periphery ^47^. Figure 2E shows the distribution of Hi-C contacts among all active adhesome loci (blue histogram) and among all active non-adhesome loci (orange histogram). Adhesome loci were found to be significantly closer in the cell nucleus compared to non-adhesome loci (*p*-value < 3e-308, two-sided Wilcoxon rank-sums test). The active adhesome genes that exhibit the highest proximity in Hi-C (top percentile) are shown in the network of Supplementary Fig. 6 (*SI Appendix*), containing 74 genes distributed on 18 chromosomes connected by 321 edges. These genes participate in many adhesome-related pathways, in particular the transmembrane receptor protein tyrosine kinase signaling pathway.

Because Topologically Associating Domains (TADs) are a well-known unit of chromosomal organization, we sought to reproduce this finding at the level of TADs instead of 250 kb genomic loci. 4,443 TADs were called at 5 kb resolution ^41^, including 134 TADs containing 132 active adhesome genes (Methods). These TADs are referred to as adhesome TADs in the sequel, and their Hi-C contacts are shown in Supplementary Fig. 7B1 (*SI Appendix*). For comparison, we identified random non-adhesome TADs that are similar in size, similarly distributed over the chromosomes, and have a similar transcriptional expression level. Supplementary Fig. 7E (*SI Appendix*) confirms that the distribution of interchromosomal contacts between active adhesome TADs significantly dominates the distribution of interchromosomal contacts between random TADs (Wilcoxon Rank-Sum test, *p*-value < 5e-124).

Interestingly, colocalization of active adhesome loci was not found at the intrachromosomal level. These loci are not significantly closer on the linear genomic sequence than randomly chosen active genes on the same chromosomes, and they do not exhibit increased intrachromosomal Hi-C contact values; see Supplementary Fig. 8 (*SI Appendix*). Consistent with this observation, active adhesome genes rarely belong to the same TADs. Overall, this finding positions interchromosomal contacts as an important level of organization for adhesome genes.

### The adhesome genes that are co-clustered in the cell nucleus are transcriptionally co-regulated

In the previous section we showed that the adhesome genes in adherent cells are mostly active as well as spatially co-clustered in the cell nucleus. To test if the co-clustered adhesome genes are coregulated, possibly to enable them to function together, we next quantified the association between co-clustering and co-regulation. To evaluate this, we quantified the co-regulation of active adhesome genes using Pearson correlation of gene regulatory profiles, containing bulk RNA-seq expression and the 48 cistromic marks described in the previous section. The resulting correlation heatmaps are shown in Fig. 3A1 and Fig. 3A2 for all 202 active adhesome genes and a random selection of 202 active non-adhesome genes with equal per-chromosome distribution, respectively. Rows and columns are arranged using hierarchical clustering with average linkage to better visualize the gene co-regulation. The heatmaps suggest that active adhesome genes are highly co-regulated, with an apparent community structure, including a strongly associated gene cluster in the center of Fig. 3A1. In Fig. 3B, comparing the distributions of Pearson correlations among active adhesome genes (blue histogram), active non-adhesome genes (orange histogram) and all non-adhesome genes (green histogram) confirmed that active adhesome genes are significantly more co-regulated in IMR-90, when compared to all active non-adhesome genes (*p*-value < 3e-308) and all non-adhesome genes (*p*-value < 3e-308) under a two-sided Wilcoxon rank-sums test.

**Fig. 3:**
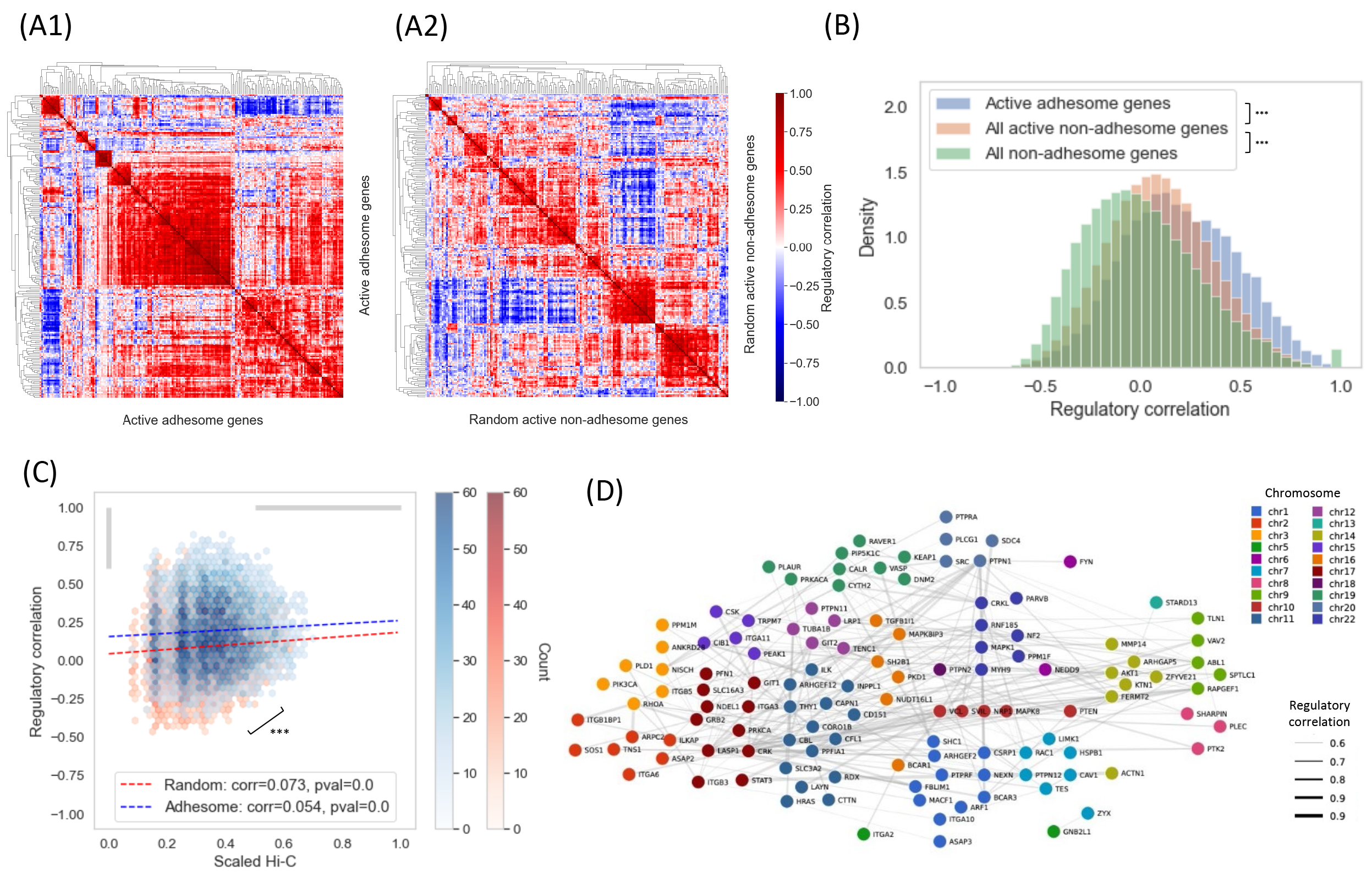
Active adhesome genes are co-regulated in IMR-90 cells. (A) Heatmaps of co-regulation for all active adhesome genes (A1) and a set of random active non-adhesome genes (A2). Co-regulation is measured using Pearson correlation between a set of 48 regulatory marks. Hierarchical clustering based on average linkage was applied to rows and columns for visualization. Overall, active adhesome genes have more similar regulatory profiles compared to other active non-adhesome genes. (B) Histograms of regulatory Pearson correlation for three groups of genes: active adhesome genes (blue), all active non-adhesome genes (orange) and all non-adhesome genes (green). Co-regulation is significantly stronger for active adhesome genes compared to active non-adhesome genes (Wilcoxon Rank-Sums test, *p*-value < 3e-308), and similarly for active non-adhesome genes compared to all non-adhesome genes (Wilcoxon Rank-Sums test, *p*-value < 3e-308). (C) 2D histogram of scaled Hi-C and regulatory Pearson correlation for active adhesome genes (blue) and a randomly selected set of active non-adhesome genes (red). The two multivariate distributions are significantly different (permutation test *p*-value < 0.01, see ^41^). For both gene sets, there is a mild, yet significant association between scaled Hi-C and regulatory Pearson correlation (the *p*-values are two-sided with respect to the exact distribution of the empirical Pearson correlation of a bivariate normal sample, see ^41^). (D) Co-regulatory network of active adhesome genes. The network skeleton is obtained by lower-thresholding the pairwise Hi-C contact values (scaled between 0 and 1) at 0.5 and lower-thresholding the pairwise regulatory Pearson correlation at 0.6, which corresponds to the grey-shaded areas of the x- and y-axes in (C). Network weights are proportional to the corresponding pairwise regulatory Pearson correlation. Genes are colored and grouped by chromosome.

This regulatory difference is partially reflected in the proximity of adhesome genes compared to other genes. Figure 3C shows the joint distribution of pairwise max-scaled Hi-C contact frequencies and Pearson regulatory correlations among active adhesome genes (blue 2D histogram) and the random non-adhesome genes (red 2D histogram). Complementary 1D histograms of the marginal distributions and 2D histograms of the joint distributions are reported in Supplementary Fig. 9A-D (*SI Appendix*). Hi-C values include both interchromosomal and intrachromosomal contacts, separately max-scaled in order to reduce the prominence of intrachromosomal contacts in the analysis. Indeed, it is known that Hi-C data is biased towards the detection of intrachromosomal contacts, which typically occur at substantially closer spatial distances than interchromosomal contacts ^48^. The histograms reveal a significant association between scaled Hi-C contacts and regulatory correlation for both gene groups (correlation test, *p*- value < 3e-308). Importantly, the joint distribution for active adhesome genes is significantly shifted upwards compared to random active non-adhesome genes (permutation test *p*-value < 0.01, see ^41^ and *SI Appendix*, Supplementary Fig. 9E). Next, for visualization purposes we selected pairs of active adhesome genes corresponding to scaled Hi-C contacts above 0.5 and regulatory correlation above 0.6 (*SI Appendix*, Supplementary Fig. 9F), resulting in a network of highly co-localized and co-regulated genes (121 genes, 255 edges) revealing important chromosomal intermingling at active adhesome gene loci. The network is displayed in Fig. 3D, with genes colored and grouped per chromosome and edge thickness representing the strength of the regulatory correlation.

### Transcription factors targeting adhesome genes in IMR-90 cells are co-clustered with active adhesome genes, facilitating transcriptional reinforcement

Because transcription factors (TFs) play a key role in the regulation and homeostasis of gene expression programs, we next sought to identify the TFs involved in the expression of active adhesome genes in IMR-90 cells. We retrieved data on 5,896,536 TF-target relationships corresponding to 333 TFs and 55,714 target genes in several tissues from hTFtarget ^16^, a curated database constructed from large-scale ChIP-seq data of human TFs in many conditions, as well as predicted interactions via TF binding site scanning. Among this data, we selected TF-target links present in several normal human lung fibroblast cell lines or samples from tissues, including IMR-90, WI38, AG04450, HPF and NHLF. While the majority of the TF-target data stems from IMR-90 (*SI Appendix*, Supplementary Fig. 10A), also considering the other human lung fibroblast cell lines provided additional TF-target relationships *(SI Appendix*, Supplementary Fig. 10B), as well as four additional TFs (SUMO1, SUMO2, UBE2I and PIAS4) which are present in the WI38 TF-target data yet absent from the IMR-90 data, but active in IMR-90 cells (*SI Appendix*, Supplementary Fig. 10C-D). All 31 TFs are shown as magenta dots on the tSNE plot of Fig. 4B, along with the adhesome genes (light green dots). We observed that 28 out of the 31 TFs present in these cell lines in hTFtarget are active in IMR-90 according to our regulatory analysis (CHD1, SNAPC4 and SPI1 were discarded as inactive). Among these 28 TFs, 26 of them target at least one active adhesome gene (SNAPC1 and SNAPC5 only target inactive adhesome genes). In the sequel, we use “adhesome TF” to refer to a TF (or the gene that codes for this TF) that targets at least one active adhesome gene.

**Fig. 4:**
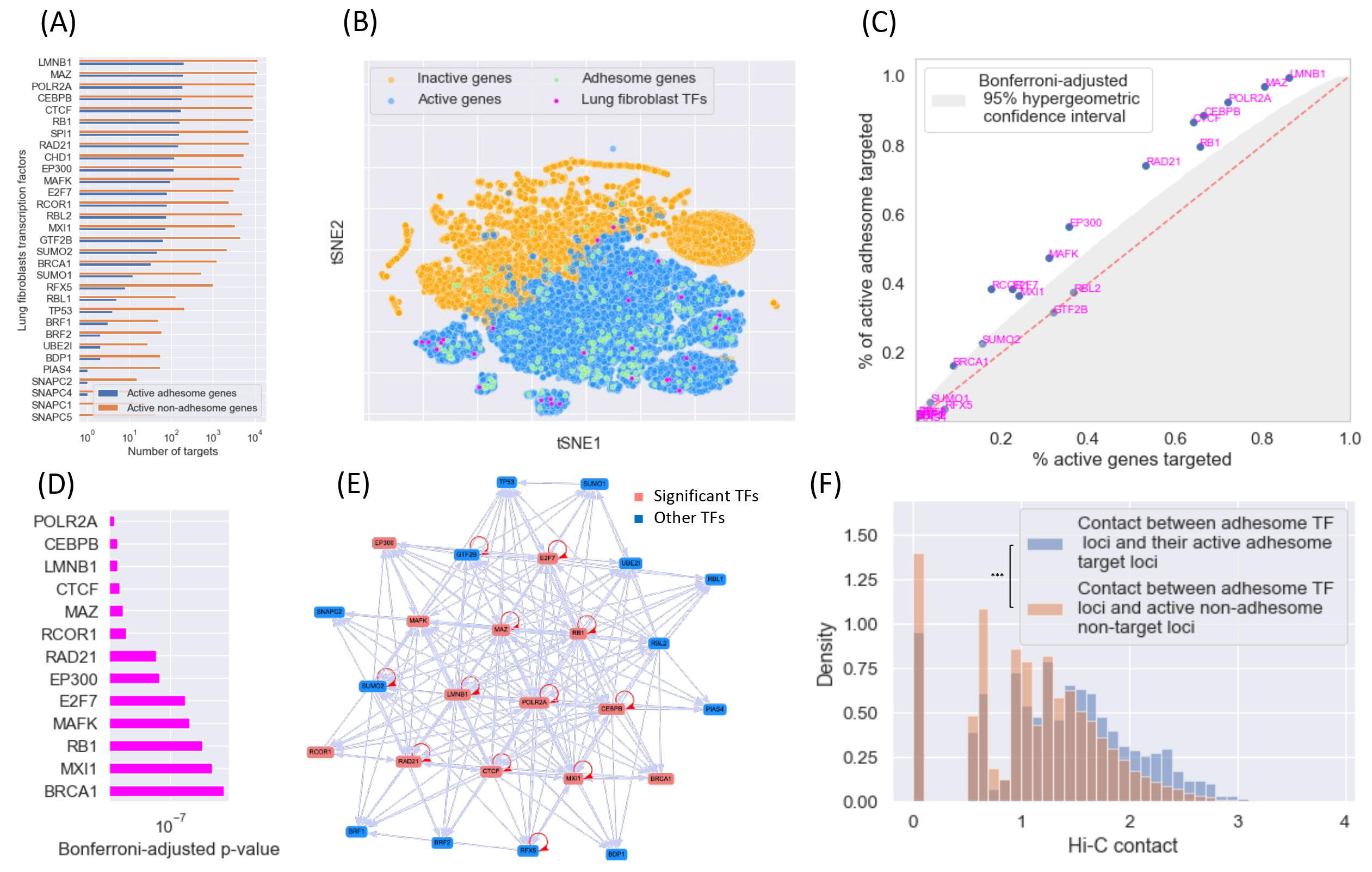
Transcription factors (TFs) targeting adhesome genes are co-regulated and co-localized in the nucleus of IMR-90 cells. (A) Bar plot of the number of active adhesome gene targets (blue) and active non-adhesome gene targets (orange) for all TFs present in normal human lung fibroblasts (namely the IMR-90, WI38, AG04450, HPF and NHLF cell lines). (B) tSNE representation of all 25,959 genes based on 48 IMR-90 regulatory marks. Orange dots correspond to inactive genes (as determined by the corresponding expression of regulatory marks), and blue dots correspond to active genes. Adhesome genes are shown in green, and TFs targeting active adhesome genes in the lung are shown in magenta. Only three of these adhesome TFs are inactive (CHD1, EP300, SNAPC4). (C) Scatter plot of the active lung fibroblast TFs that target active adhesome genes showing the proportion of total active genes targeted (x-axis) and the proportion of active adhesome genes targeted (y-axis) for each TF. The shaded region corresponds to the Bonferroni-corrected 95% hypergeometric confidence interval conditional on the number of active genes targeted by each TF ^41^. The TFs in this figure correspond to all TFs in (A) except for inactive TFs (CHD1, EP300, SNAPC4) and active TFs targeting inactive adhesome genes only (SNAPC1, SNAPC5). (D) Bar plot of Bonferroni-adjusted hypergeometric *p*-values for all significant TFs from (C) at the 95% level. (E) Network of interactions between all active lung fibroblast TFs targeting active adhesome genes. TFs identified as significant in (C)-(D) are represented as red nodes, and the remaining TFs are represented as blue nodes. Red self-loops indicate that a TF targets itself. (F) Histograms of interchromosomal Hi-C contact values between all active adhesome TF loci (including loci of both significant and non-significant TFs identified in (C)-(D)) and their active adhesome target loci (blue) as well as all other active non-adhesome, non-target loci (orange). Active adhesome TF loci are significantly closer to their active adhesome target loci compared to the other loci (*p*-value < 4e-90, Wilcoxon Rank-Sums test).

There is heterogeneity in the overall number of active genes targeted by these TFs, with some TFs targeting less than 50 active genes (UBE2I, SNAPC1, SNAPC2, SNAPC4) and other TFs targeting more than 10,000 active genes (LMNB1, MAZ, POLR2A); see Fig. 4A. This heterogeneity may reflect both the specificity of a given TF as well as potential missing data from hTFtarget. Interestingly, active adhesome genes tend to be more targeted by the identified TFs than similar random active non-adhesome genes (Wilcoxon Rank-Sum test, *p*-value < 2e-10; see Supplementary Fig. 11A in *SI Appendix*), supporting the potential role of the identified TFs in adhesome regulation. Conditional on their total number of active gene targets, we identified 13 TFs whose target genes contained adhesome genes at a significantly higher proportion than expected under a hypergeometric test (family-wise error rate control, FWER, at 5% with Bonferroni correction). These 13 TFs lie outside of the shaded 5% Bonferroni-corrected hypergeometric confidence region in Fig. 4C. These include TFs that have been identified to play a role in adhesion in previous research: LMNB1 plays an important role in cellular response to mechanical stress ^49^, RAD21 regulates the cadherin and cell adhesion pathway ^50^, and EP300 actively controls cell adhesion ^51^, among others. The factors POLR2A, CTCF, MAZ, CEBPB and LMNB1 have the smallest FWER-adjusted *p*-values and target more than 80% of active adhesome genes, suggesting a high level of redundancy and co-regulation (Fig. 4D).

The 13 significant adhesome TF genes correspond to 15 distinct 250 kb genomic loci, and the 13 remaining adhesome TF genes correspond to 14 distinct genomic loci. In aggregate, these 29 adhesome TF loci are shown in Fig. 4F to be significantly closer to their respective active adhesome target loci than to other active non-adhesome, non-target loci (Wilcoxon Rank-Sums test, *p*-value < 4e-90). This observation is also confirmed by an analysis of the 3D proximity of active adhesome target genes for each adhesome TF separately, as shown in Supplementary Fig. 11B-C (*SI Appendix*). Most of these TFs exhibit a significant colocalization of their active adhesome gene targets compared to randomly selected active non-target genes on the same chromosomes. This suggests that active adhesome genes and the genes coding for their TFs co-localize in the nucleus.

We also compared the proportion of adhesome genes targeted by the 13 significant TFs in other cell lines for which sufficient TF-target data was available in hTFtarget (*SI Appendix*, Supplementary Fig. 12A-B), including cell lines with adherence properties (A549, HepG2, HEK293) and a non-adherent cell line (GM12878). Interestingly, some of the significant TFs we identified are quite specific to lung fibroblasts, such as LMNB1, RB1, RCOR1, BRCA1, and, to a lesser extent, MAZ and CEBPB. In other cell lines, these TFs tend to target a lower proportion of adhesome genes (*SI Appendix*, Supplementary Fig. 12C). On the contrary, some TFs that do not target adhesome genes in IMR-90 may target a high number of adhesome genes in other cell lines (*SI Appendix*, Supplementary Fig. 12D, bottom rows). Only CTCF targets a high proportion (>86%) of adhesome genes in all the cell lines under consideration.

Next, we built a regulatory network of adhesome TFs using the TF-target link data (Fig. 4E). TFs identified as significant in Fig. 4C-D are represented as red nodes, while the remaining TFs are represented as blue nodes. The network is densely connected (242 directed edges among the 26 active TFs targeting active adhesome genes, density = 0.37); notably, 12 TFs activate themselves, including POLR2A, CTCF, MAZ, CEBPB and LMNB1. These findings support a high degree of self-reinforcement in the regulation of adhesome genes in IMR-90 cells. TFs found to significantly target active adhesome genes are also the most central TFs in this network (XL-minimal hypergeometric test ^52^ with X=1 and L=20, *p*-value < 6e-4). Centrality of a TF was measured via its Katz centrality score, which computes the relative influence of a TF through its targets in the directed network (*SI Appendix*, Supplementary Fig. 10E).

### Clustering analysis identifies interacting regions of tightly co-localized and co-regulated adhesome genes, their transcription factors, as well as genes of the ubiquitin-proteasome system

The above analysis suggests that adhesome genes and the genes coding for their transcription factors, many of which can activate themselves, are tightly co-localized in the cell nucleus of IMR- 90 cells. In order to further dissect the spatial organization of adhesome genes as well as identify non-adhesome genes that are co-clustered and co-regulated with adhesome genes, we ran the Large Average Submatrix (LAS) algorithm ^26,53^ on the IMR-90 interchromosomal Hi-C matrices to identify specific groups of genes that are spatially clustered in the cell nucleus despite lying on different chromosomes (Methods). Seeding the algorithm in active adhesome loci, the LAS algorithm greedily identifies contiguous submatrices in which the average contact frequency is significantly higher than expected by chance ^41^. The output of the LAS algorithm is a list of interchromosomal interactions between genomic regions and is shown in Fig. 5A for the example of the Hi-C map between chromosomes 19 and 20.

**Fig. 5:**
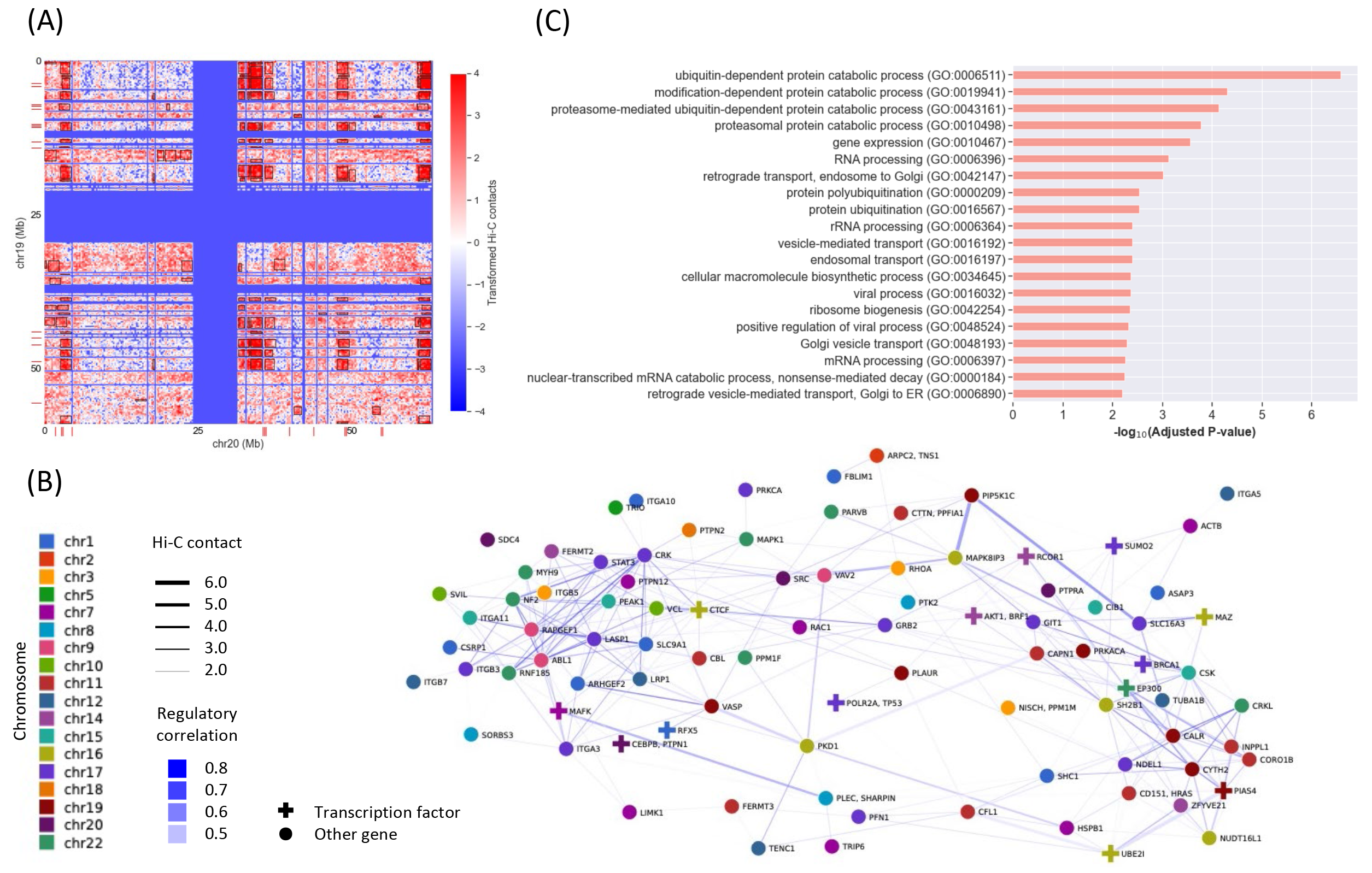
Analysis using the LAS algorithm identifies interacting regions in Hi-C of tightly co-localized and co-regulated adhesome genes, their transcription factors, as well as non-adhesome genes of the ubiquitin-proteasome system. (A) Heatmap of transformed Hi-C contact frequencies between genomic loci on chromosome 19 and chromosome 20 (frequencies were log-transformed and centerdized using the mean and standard deviation of contact values across the entire genome). Adhesome loci are indicated by red ticks on the x-axis (for chromosome 20) and the y-axis (for chromosome 19). Significant submatrices identified by the LAS algorithm with an average Hi-C contact value higher than 2 are shown in black boxes. (B) Network of adhesome LAS nodes obtained from keeping all edges that are detected by the LAS algorithm and that have a regulatory correlation in the top 15%-percentile. These edges are in the upper right quadrant of Supplementary Fig. 15C (*SI Appendix*), as defined by the red dashed marginal thresholds. Adhesome LAS nodes are colored by chromosome, and a cross indicates that the node contains at least one TF targeting active adhesome genes. Edge width corresponds to the average Hi-C contact value and edge color corresponds to the regulatory correlation. (C) Significant Gene Ontology (GO) terms associated to active non-adhesome genes belonging to genomic regions that interact with adhesome interacting regions. Only genes having a strong positive regulatory correlation (upper decile) with the adhesome genes they interact with (in LAS) are considered. A GO term is reported as significant if its FDR-adjusted *p*-value is lower than 1e-2. Many of the identified GO terms are associated with the ubiquitin-proteasome system.

The LAS algorithm identified 24,809 large average submatrices, 5,540 of which contained active adhesome genes or adhesome TF genes. In the remainder of this article, a set of contiguous genomic loci corresponding to either the rows or the columns of a large average submatrix identified by LAS is called an interacting region; if an interacting region contains at least one adhesome gene or an adhesome TF, it is called an adhesome interacting region. The LAS submatrices projected onto the Hi-C map containing only the active adhesome loci and TF adhesome loci are shown as black boxes in Supplementary Fig. 13A (*SI Appendix*). The adhesome interacting regions contain 1 to 3 adhesome genes (*SI Appendix*, Supplementary Fig. 13B-C). Supplementary Fig. 13A (*SI Appendix*) shows the resulting adhesome LAS network, where the nodes (which we call adhesome LAS nodes) are adhesome genes, and two adhesome LAS nodes are linked by an edge if they are linked by an LAS submatrix. Adhesome genes belonging to the same chromosome are merged into one adhesome LAS node if they always co-appear in the LAS submatrices. The resulting network contains 118 nodes with 2,250 edges, with an average degree of 38. Supplementary Fig. 14 (*SI Appendix*) compares various properties of the adhesome LAS network to a null distribution obtained by building 500 LAS networks based on a random active non-adhesome gene set of the same size (∼200 genes), confirming that adhesome genes and adhesome TF genes interact more strongly spatially than expected by chance.

To mitigate some of the noise associated with using Hi-C contact frequencies alone, edges in the adhesome LAS network were weighted (and then pruned) by the regulatory correlation between the two adhesome LAS nodes as follows: for a given pair of adhesome LAS nodes, the regulatory correlation is equal to the average of Spearman correlations between the regulatory profiles of all tuples of adhesome genes such that one of the adhesome genes belongs to the first LAS node and the other gene belongs to the second LAS node. This allowed us to identify chromosome intermingling regions with coordinated activity, which might be controlled by the same set of TFs or epigenetic marks. The scatter plot of interactions between adhesome LAS nodes based on average Hi-C contact value (x-axis) and regulatory correlation (y-axis) is shown in Supplementary Fig. 15C (*SI Appendix*), where red dots correspond to interactions involving at least one adhesome TF. By construction, all edges in the adhesome LAS network have an average Hi-C contact value higher than 2 (*SI Appendix*, Supplementary Fig. 15A). We used a lower threshold on the regulatory correlation (85% quantile of its marginal distribution in the network, see *SI Appendix*, Supplementary Fig. 15B) to prune edges of the network corresponding to weak or negative co-regulation. The resulting network has 92 nodes (100 adhesome genes) and 338 edges (average degree = 7.3) and is displayed in Fig. 5B, where nodes are colored by chromosome, edge widths reflect the average Hi-C contact frequency between nodes, and edge colors reflect the regulatory correlation between regions. The network includes 14 adhesome TFs (displayed as crosses), 8 of which were found to be significant in our previous analysis (BRCA1, CEBPB, CTCF, EP300, MAFK, MAZ, POLR2A, RCOR1). Note that MAZ and POLR2A target 100% of genes in the network, while CTCF and CEBPB target more than 85% of genes, EP300, RCOR1 and MAFK target more than 35% of genes, and BRCA1 targets 12% of genes. Interestingly, the maximal cliques of this network involve at most 5 chromosomes (*SI Appendix*, Supplementary Fig. 15D), which is consistent with the observation that the simultaneous interaction of genes from more than five chromosomes is physically unlikely ^46,48^.

The LAS analysis also allows us to identify non-adhesome genes that are co-clustered and co-regulated with adhesome genes. Towards this, we considered the genes in non-adhesome interacting regions that interact with adhesome interacting regions. This gene list was further refined by only keeping active genes that showed a strong positive regulatory correlation (upper decile) with the adhesome genes in the LAS submatrices in which they were found (see *SI Appendix*, Supplementary Fig. 16A). This yielded a list of 2,826 active non-adhesome genes that were spatially co-clustered and co-regulated with active adhesome genes. We performed a gene set enrichment analysis on these genes. The significant Gene Ontology (GO) terms (at the 1e-2 level after adjustment for False Discovery Rate) are reported in the bar chart of Fig. 5C. Interestingly, many of these GO terms are related to the ubiquitin-proteasome system, which has previously been shown to regulate focal adhesions ^54^, as well as terms related to vesicle transport, which plays a role in the regulation of adhesome receptors in the context of 3D cell migration ^55^. Repeating this analysis using a more stringent regulatory correlation threshold (top percentile) selected 755 active non-adhesome genes and resulted in enrichment for the same GO terms (see *SI Appendix*, Supplementary Fig. 16B-C).

### FISH experiments confirm the clustering of adhesome genes and their transcription factors

To validate our computational findings of the proximity of adhesome genes in the nucleus of IMR- 90 cells, we retrieved multiplexed error-robust fluorescence in situ hybridization (DNA-MERFISH) data ^22^, which imaged 1,041 genomic loci in 5,400 IMR-90 cells across 5 biological replicates. The probe loci used for DNA-MERFISH were ∼30 kb in size, covering the 22 autosomes and chromosome X approximately uniformly, with each chromosome containing 30-80 loci. After filtering out probe loci with missing values as well as loci located on chromosome X, we obtained 990 probe loci in 3,533 cells. Supplementary Fig. 17A (*SI Appendix*) shows the 3D scatter plot of all probe loci in a specific cell; the nucleus dimensions are consistent with the typical size of IMR- 90 nuclei (∼30μm). We identified 82 MERFISH probe loci falling within 500 kb upstream or downstream of an active adhesome gene or an adhesome TF gene on the genomic sequence. These probe loci cover 95 active adhesome genes and adhesome TF genes (i.e., 42%) and are referred to as adhesome probe loci in the sequel.

We computed the average Euclidean distance in 3D among the 82 adhesome probe loci across all 3,533 cells. To account for different cell sizes, probe-probe distances in a cell were normalized by the maximum probe-probe distance in that cell (see *SI Appendix*, Supplementary Fig. 17B). Because some probe loci may be missing in some of the cells, the average distance for a given pair was taken with respect to the number of cells in which that pair was observed (see *SI Appendix*, Supplementary Fig. 17C). The resulting histogram of average cell-normalized Euclidean distances between adhesome probe loci is shown in Supplementary Fig. 17D (*SI Appendix*). Consistent with previous findings ^48^, these distances are well-correlated with the scaled Hi-C frequencies (Pearson correlation of 0.494 for interchromosomal distances and 0.787 for intrachromosomal contacts; see *SI Appendix*, Supplementary Fig. 17E). For comparison, we also considered the average distances among a group of 82 randomly selected active non-adhesome probe loci (a probe locus was defined as active if it was no more than 500 kb upstream or downstream from an active gene in IMR-90). We constrained the number of random probe loci per chromosome to be the same as the number of adhesome probe loci per chromosome, to allow for a fair comparison. The resulting distance heatmaps are shown in Fig. 6A confirming that adhesome probe loci are closer compared to non-adhesome probe loci; this is even more apparent if we restrict the heatmaps to probe loci that are less than 100 kb away from adhesome genes (Fig. 6B). The histograms of average cell-normalized Euclidean distances in Fig. 6C confirm that active adhesome probe loci are significantly closer compared to all other active non-adhesome probe loci (Wilcoxon rank-sums test, *p*-value < 3e-308). These findings remained valid when we restricted the analysis to only MERFISH probe loci that overlap with active adhesome genes (19 probes covering 19 active adhesome genes), as shown in Supplementary Fig. 18 (*SI Appendix*).

**Fig. 6:**
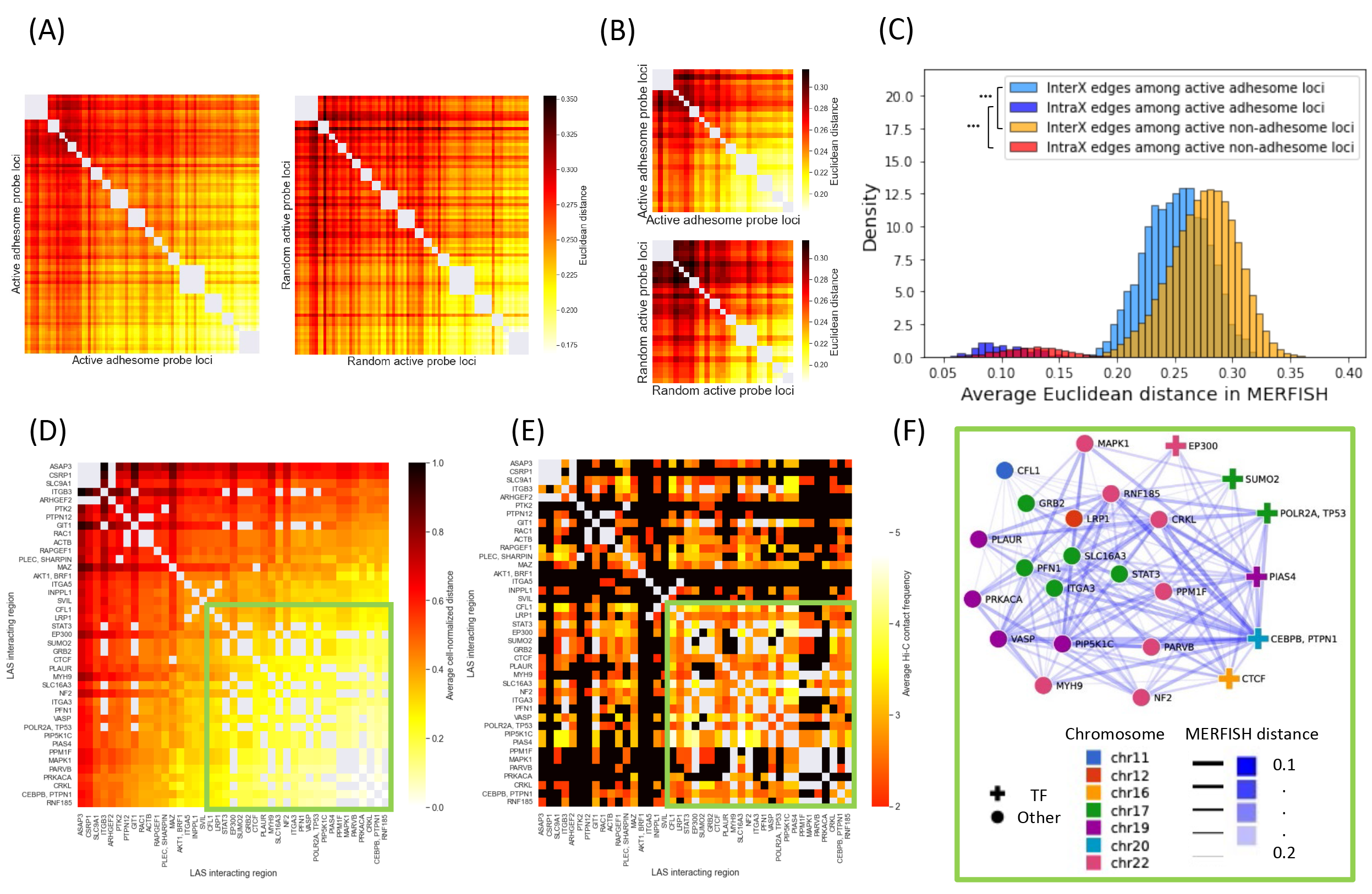
DNA-MERFISH data confirms the proximity of the identified co-regulated adhesome genes and their transcription factors. (A) Left: Heatmap of cell-normalized MERFISH Euclidean distance between active adhesome probe loci. Active adhesome probe loci correspond to probe loci within 500 kb upstream or downstream of an active adhesome gene or an adhesome transcription factor gene. Loci are ordered according to chromosome and genomic position. Cell-normalized Euclidean distance between loci i and j is the cell-normalized Euclidean distance between i and j averaged over all cells in the MERFISH data set containing probes i and j. Cell-normalization corresponds to dividing the physical distance between i and j in cell c by the maximum probe-probe distance in that cell. Right: Heatmap of Euclidean distance between selected random non-adhesome loci. The number of random probe loci per chromosome is chosen equal to that of adhesome probe loci, to allow for a fair comparison. (B) Top: Same as left figure of (A) but only considering probe loci that are located within 100 kb upstream or downstream of an active adhesome gene or an adhesome transcription factor gene. Bottom: Same as right figure of (A) but only considering probe loci that are located within 100 kb upstream or downstream of random non-adhesome active genes. (C) Histograms of average Euclidean distance between active adhesome probe loci (light blue: interchromosomal distances, dark blue: intrachromosomal distances) and between all active non-adhesome probe loci (orange: interchromosomal distances, red: intrachromosomal distances). Active adhesome probe loci are significantly closer than active non-adhesome probe loci (Wilcoxon rank-sums test, *p*-value < 3e- 308). (D) Heatmap of average Euclidean distance between nodes in the adhesome LAS network of Fig. 5C. Only nodes containing at least one active adhesome probe locus are shown. Entries corresponding to nodes on the same chromosome are colored in grey. Rows and columns are ordered using hierarchical clustering with single linkage. (E) Heatmap of average Hi-C contact values between nodes in the adhesome LAS network of Fig. 5C. Only nodes containing at least one active adhesome probe locus are shown. Entries corresponding to nodes on the same chromosome are colored in grey; entries corresponding to interactions that were not identified by LAS are set to 0 and colored in black. Rows and columns are ordered as in (C). (F) Subnetwork of adhesome LAS nodes corresponding to the lower right quadrant of (C) and (D) (highlighted in green). Nodes are colored by chromosome and represented with a cross if they contain at least one active adhesome transcription factor. Edge width and color reflect Euclidean proximity.

The MERFISH data also allows us to validate some of the interchromosomal interactions between active non-adhesome genes and active adhesome genes. In the previous section, 2,826 active non-adhesome genes were found to cluster and to be co-regulated with 135 active adhesome genes. We mapped these 2,961 genes to the closest MERFISH probe within 100 kb upstream or downstream on the genomic sequence. This resulted in 135 active non-adhesome genes (associated with 61 probes) and 17 active adhesome genes (associated with 16 probes); all other genes have no associated probe locus.

The distribution of MERFISH distances between these gene pairs is shown in Supplementary Fig. 19A-B in red (*SI Appendix*). Comparing this distribution to the distribution of distances between active adhesome MERFISH probes across LAS clusters (Supplementary Fig. 19A) and within the same LAS cluster (Supplementary Fig. 19B) confirms that the identified active non-adhesome genes are highly co-clustered with the active adhesome genes they are predicted to interact with (Wilcoxon Rank-Sums test, *p*-value < 2e-29). We visualized this distance information by creating a bipartite network in which active adhesome genes are linked to active non-adhesome genes if the distance between these genes falls in the lower quartile of the distance distribution; after discarding isolated nodes, we obtained a bipartite network with 52 nodes (10 active adhesome genes and 42 active non-adhesome genes) and 50 edges (*SI Appendix*, Supplementary Fig. 19B). Interestingly, various of the identified non-adhesome genes are known to be related to adhesion, including PEPD (involved in collagen metabolism), FXYD5 (involved in down-regulation of E-cadherin) and ACTN4 (actin anchor chain). In addition, COLGALT1 is an enzyme that post-translationally modifies collagen proteins. Of interest are also the interactions between ACTN4 and the adhesome genes ITGA3 (integrin alpha 3) and PFN1 (small actin binding protein). We also highlight that many non-adhesome genes in the graph interact with the adhesome gene CRKL (degree = 22), a gene that plays an important role as a kinase to modulate the focal adhesions^56^.

Finally, we used the processed MERFISH data to validate the interactions between nodes of the adhesome LAS network. The scaled Euclidean distances between nodes in the adhesome LAS network are shown in Supplementary Fig. 20 (*SI Appendix*), where the distance between two adhesome LAS nodes is defined by averaging the distances between all DNA-MERFISH probes associated to the first LAS node and all DNA-MERFISH probes associated to the second LAS node. Nodes without an associated DNA-MERFISH probe are represented as grey rows/columns. Interactions with very low Euclidean distance (darkest entries of the heatmap) correspond to intrachromosomal interactions, which we excluded from the subsequent analysis because of the focus of this work on interchromosomal interactions. The resulting MERFISH Euclidean distance matrix among adhesome LAS nodes with associated MERFISH probes is shown in Fig. 6D. Rows and columns are ordered using hierarchical clustering with single linkage (cophenetic correlation ∼ 0.4, see *SI Appendix*, Supplementary Fig. 21A). The corresponding matrix of average transformed Hi-C contact frequencies is shown in Fig. 6E. Entries corresponding to edges that were not identified by the LAS algorithm are set to 0 and shown in black. It is apparent by considering the heatmaps in Fig. 6E-F that there exists a group of 24 tightly interacting adhesome LAS nodes in the lower right quadrant. Supplementary Fig. 21B-D (*SI Appendix*) confirm that these adhesome LAS nodes are also co-regulated. The network of these tightly interacting adhesome LAS nodes is displayed in Fig. 6F; it contains 24 nodes (corresponding to 26 adhesome genes on 7 chromosomes) and 169 edges (average degree = 14.1). Nodes are colored by chromosome and edges are colored and sized according to MERFISH Euclidean distance. The genes in the network mostly code for adaptor proteins (CRKL, GRB2, NF2, PARVB), or are involved in actin regulation (CFL1, VASP, MYH9, PFN1) and adhesion receptors (LRP1, ITGA3, PLAUR). The network also contains 7 adhesome TFs, including 4 significant adhesome TFs: POLR2A, CTCF and CEBPB, which target more than 88% of genes in the subnetwork, and EP300, which targets 50% of genes in the subnetwork. Protein-protein interaction (PPI) data retrieved from the physical STRING v11 interactome ^57^ shows that some of these proteins also interact physically, such as ITGA3-PLAUR, CTCF-EP300, and MYH9-CTCF (*SI Appendix*, Supplementary Fig. 21E).

## Discussion

Cells have evolved to organize their receptors on the plasma membrane for signal sensing ^1–6^. In addition, the genetic material is packed within the cell nucleus to encode the various cellular proteins ^12–15^. In recent years two aspects of spatial clustering have been identified, one at the level of receptors for optimized signal sensing, and the other at the level of genes for optimized transcriptional regulation. While these two layers of clustering may appear independent of each other, we hypothesized that there may exist a coupling between the two spaces. The main premise of this hypothesis is due to cells having to detect the extracellular signals through their receptors and needing to transcriptionally reinforce the sensing machinery for rapid adaptation ^7^. For such cellular adaptation processes it is prudent that the genes that code for the required proteins are spatially clustered for their co-regulation, thereby enabling rapid cellular response.

In light of recent progress in genomic and proteomic technologies, one now has access to both the protein modules that participate in signal sensing ^8–11^ and genome-wide maps of the spatial arrangement of genes within the cell nucleus ^19,20^. To test our hypothesis, we took advantage of the publicly available datasets in IMR-90 cells and carried out an in-depth analysis of the coupling between protein clustering and gene clustering in the context of the adhesome in an adherent fibroblast cell line. The adhesome is an essential matrix sensing machinery and more than 200 proteins that are part of the adhesome have been well-annotated ^36,37^. The list of adhesome proteins were curated enabling us to map these proteins to their corresponding sequences on the genome. Importantly, high-resolution chromosome conformation maps of IMR-90 cells ^20^ provided us with the ability to explore the spatial arrangement of genes that code for the adhesome proteins. When interrogating these Hi-C maps, we identified that the adhesome genes were spatially clustered in a transcription-dependent manner, with transcriptionally active adhesome genes being co-clustered. These results suggested to us that the role of the coupling between the receptor proteins and their coding genes could be to reinforce the cellular sensing machinery. To verify this, we next tested if the transcriptional regulators of the active adhesome genes were spatially co-localized within the adhesome gene clusters. Upon matrix sensing, various downstream signal processes are triggered including some of the adhesome proteins serving as transcription factors and cofactors. Importantly, we found that many of these transcription factors and co-factors are co-localized within the adhesome gene clusters, suggesting a tight transcriptional control and the possibility to rapidly reinforce cellular responses. Extending our analysis to identify non-adhesome genes that are co-localized within the adhesome gene clusters, we identified genes of the ubiquitin-proteasome system – a system known to be involved in the regulation of focal adhesions ^54^. In order to test if the adhesome gene clusters identified using population-level Hi-C maps are physically close-by in single cells, we took advantage of recent DNA-MERFISH data in IMR-90 cells, where over 1,400 probe loci were mapped including many of the adhesome genes ^22^. Our analysis of this dataset validated that many of the adhesome genes and their transcription factors are spatially clustered also at the single-cell level.

Collectively, our results highlight an important link between protein clustering on the membrane and gene clustering in the cell nucleus. Typical matrix sensing involves the clustering of receptors in the first few minutes, followed by adhesome maturation and cellular responses in the next tens of minutes, which is then followed by cell behavioral responses in subsequent hours to days ^58^. A critical step for the receptor maturation and cellular adaptation is an optimal transcriptional regulatory mechanism. Since the downstream target genes of signal sensing are located on different chromosomes, an efficient mechanism of transcription control is through the spatial clustering of the involved genes, their transcription factors, and the transcription machinery. Our findings highlight a general principle of how such optimal transcriptional regulatory mechanisms are established in living cells. We suggest that our results are not specific to the adhesome, but rather a general evolutionary-conserved mechanism of receptor-mediated rapid cellular adaptation to various microenvironmental signals.

## Methods

### Adhesome data

Adhesome data was retrieved from http://adhesome.org/, a literature-based protein-protein interaction network made of known interactions and cellular components constituting the focal adhesion complex in mammalian cells ^36,37^. Study results for multiple mammalian cell types were used to construct this database, which is not specific to the IMR-90 cell line. The network includes 150 intrinsic components and 82 associated components, with a total of 6,542 interactions (*SI Appendix*, Supplementary Fig. 2B). The distinction between intrinsic and associated components is based on the location of the molecule: intrinsic components reside in the adhesion complex, while associated components transiently belong to the adhesion complex. Performing a Gene Set Enrichment Analysis (GSEA) on this list of 222 genes identifies several significant adhesion-related GO terms, including cell-matrix adhesion (GO:0007160), transmembrane receptor protein tyrosine kinase signaling pathway (GO:0007169) and integrin-mediated signaling pathway (GO:0007229).

### Regulatory data and processing

IMR-90 bulk mRNA-seq data collected by ^38^ was retrieved from the Gene Expression Omnibus (GEO) under the accession number GSE16256 (sample GSM438363). Data from 48 regulatory marks was also obtained from ^39^ who collected it from ENCODE ^24^, Roadmap Epigenomics ^23^, and the GEO, and released it on the TargetFinder GitHub page (https://github.com/shwhalen/targetfinder). In order to obtain the genomic profile for a gene, the number of peaks overlapping the gene’s location on the hg19 reference genome (obtained from ^40^) was calculated for each feature using the Python pybedtools library. The profile vector of each gene was then divided by the gene length to account for the random incidence bias. For each feature, the feature matrix was then log(1+1000000x)-transformed, centered and standardized by computing the mean and standard deviation of the feature across all genes in the genome. To determine gene activity in GM12878, we also retrieved and processed as described above bulk RNA-seq data from the GEO under the accession number GSE90276 ^24^ (hg19-aligned gene quantifications in log(1+TPM) format from samples GSM2400247 and GSM2400248) as well as 100 regulatory marks from ^39^. The likely category of histone ChIP-seq marks (active, repressive, unknown) was retrieved from ^59^ and ^60^, and the category of transcription factors and other protein ChIP-seq marks were inferred from protein function.

### Hi-C data and processing

The Hi-C contact frequency matrices for IMR-90 and GM12878 from ^20^ were obtained from the GEO (accession number GSE63525) at 250kb resolution and mapping quality MAPQ>30. Intrachromosomal matrices were corrected for locus-specific bias using the Knight-Ruiz matrix balancing algorithm ^45^ using the Juicebox software (https://github.com/aidenlab/Juicebox). Interchromosomal matrices were also corrected for locus-specific bias using a matrix balancing approach (see ^20^ for more details). Namely, we ran the Knight-Ruiz matrix-balancing algorithm on the genome-wide contact matrix at resolution 250kb (GWKR normalization). Following normalization, centromeric regions retrieved from ^40^ were filtered out in each interchromosomal Hi-C matrices, along with peri-centromeric regions within 2Mb of the centromere and outliers based on row and column sums (outside of the 1.5xIQR, where IQR denotes the interquartile range). Repeat regions from ^40^, as well as regions already masked in ^20^ were also removed from the analysis. The final Hi-C matrix was log(1+x)-transformed. When we run the LAS algorithm, we further centerdize the final Hi-C matrix using the mean and standard deviation of contact values across the entire genome.

### Topologically Associating Domains

We followed the steps outlined in ^61^ to call TADs with the Arrowhead software ^20^ at a resolution of 5 kb, using a Knight-Ruiz (KR) normalization of intrachromosomal Hi-C maps. TAD search was limited to a sliding window of size 2,000 along the Hi-C matrix diagonal. In a subsequent processing step, we removed nested TADs and merged overlapping TADs. Overall, this process identified 4,443 TADs located on different chromosomes (except for chromosome 9), covering ∼35% of the genome. Interestingly, the resulting average TAD length is 225 kb, which is similar to the 250 kb Hi-C resolution used throughout this article. Illustrations and descriptive statistics of the TAD calling process can be found in Supplementary Fig. 22 (*SI Appendix*). Diagnostic metrics ^61^ showed that the resulting TAD list outperformed TAD lists obtained using different resolutions (5kb, 10 kb, 25 kb) or different Hi-C processing methods (KR normalization, no normalization). Computing the Measure of Concordance between these TAD lists revealed that normalization had a limited impact on TAD calling. Then, investigating the enrichment of CTCF, RAD21, and SMC3 (cohesin complex proteins) at the TAD boundaries showed that TAD partitions obtained at the 5 kb resolution exhibited the highest quality. Finally, enrichment of H3K36me3 (active mark) and H3K27me3 (repressive mark) in the TAD lists (measured as the log10-ratio between their ChIP-seq signals and compared with a null distribution derived from random permutations) confirmed the high quality of TAD lists obtained at the 5 kb resolution. These diagnostic metrics are shown in Supplementary Fig. 23 (*SI Appendix*).

### DNA-MERFISH data

We retrieved DNA-MERFISH data from ^22^, where 1,041 genomic loci were simultaneously imaged in 5,400 IMR-90 cells across 5 biological replicates with a detection efficiency of ∼80% for each locus, yielding on average ∼1,700 probe loci detected in each cell. The probe loci used for DNA-MERFISH are ∼30 kb in size and cover the 22 autosomes and chromosome X approximately uniformly, with each chromosome containing 30-80 loci. The data is freely available on Zenodo (https://zenodo.org/record/3928890#.Yj07vZrMJqs). After filtering out probe loci with missing values or that were located on chromosome X, we obtained 990 probe loci in 3,533 cells to use in our analysis. For this analysis, we considered loci and gene locations with respect to the hg38 reference genome; the gene location data was retrieved from the UCSC Genome Browser ^40,42^.

### Transcription factors-target data

Transcription factor (TF) target data was retrieved from ^16^ containing a curated TF-target dataset from large-scale ChIP-seq data of human TFs (7,190 samples of 659 TFs) in 569 conditions (399 cell lines, 129 classes of tissues or cells, and 141 kinds of treatment). Additional inferred TF-target links were obtained from ^16^ based on epigenomic modification status and TF binding site scanning. We scraped this data for several tissues, including the lung, from the online repository of hTFtarget (http://bioinfo.life.hust.edu.cn/hTFtarget/#!/) using the Python package Requests.

### Protein-protein interaction data

Protein-protein interaction data used in Supplementary Fig. 21E (*SI Appendix*) was retrieved from the STRING v11 (https://string-db.org/cgi/input?sessionId=bSN57PdA9N4N&input_page_show_search=on) interactome ^57^, restricted to physical interactions only. The interaction strength is a measure of confidence combining probabilities from different evidence channels (experiments, text mining, data bases).

### Statistics

*Correlation test*: In Fig. 3C and Supplementary Fig. 17E (*SI Appendix*), we used a two-sided correlation test described in ^62^ to assess the significance of the Pearson correlation between scaled Hi-C contact values and the regulatory correlation based on 48 regulatory marks for two groups of genes: active adhesome genes (blue) and a randomly selected set of active non-adhesome genes (red).

*Multivariate two-sample test*: In Fig. 3C, we also tested for the difference between the joint distribution of scaled Hi-C contact values and Pearson regulatory correlation for the two gene groups of interest: active adhesome genes (blue) and the randomly selected set of active non-adhesome genes (red). We used a two-sample test based on the number of nearest neighbor type co-incidences, as described in ^63^. We created a 1-Nearest Neighbor graph among genes from both groups using the Euclidean distance to determine neighboring relationships. Then, we computed the number of edges between genes of the same group. We compared this statistic to a simulated null distribution where we computed the same statistic under random permutation of gene labels, for 100 different permutations (see *SI Appendix*, Supplementary Fig. 9E). We reported the one-sided *p*-value for that null distribution.

*Hypergeometric test*: In Fig. 4C, we plotted each TF of interest based on the percentage of genes it targets among all active genes (x-axis) and the percentage of genes it targets among all active adhesome genes (y-axis). To identify important TFs, we adopted a null hypergeometric model for each TF, conditionally on the number of genes it targets among all active genes. Under that null model, the number of active adhesome genes targeted by the TF follows a Hypergeometric(M,n,N) distribution where M is the total number of active genes, n is the total number of active adhesome genes and N is the number of active genes targeted by the TFs. For each TF, we computed the corresponding one-sided *p*-value. In Fig. 4C, we also shaded in grey the area corresponding to non-significant TFs at the 0.05 significance level, after applying a Bonferroni correction to control the Family-Wise Error Rate (FWER).

*Cosine similarity*: For hierarchical clustering, we can measure affinity between two genes using the cosine similarity between their feature vectors. The cosine similarity between two vectors u and v is equal to <u,v>/(||u||·||v||). It is a scoring metric to measure the similarity between u and v, ranging from −1 (vectors in opposite directions) to 1 (vectors aligned in the same direction); a value of 0 corresponds to orthogonal vectors.

### Large Average Submatrices (LAS) algorithm

The LAS algorithm ^53^ takes a real-valued data matrix X (of size m-by-n) as input and outputs contiguous submatrices U (of size k-by-l) that have a high average, τ, when compared to independent standard Gaussian entries. Therefore, the Hi-C contact frequencies were log-transformed and standardized using the mean and standard deviation of the non-zero contact frequencies across the entire genome. The resulting distribution of non-zero transformed Hi-C contact values in the entire genome is reported in Supplementary Fig. 24A (*SI Appendix*); we note that the LAS algorithm is robust to slight violations of the normality assumption.

We used a version of the LAS algorithm implemented in ^26^, which specifically detects contiguous submatrices rather than arbitrary submatrices. A detailed description of the algorithm is provided in the Supplementary Information of ^26^. In our work, the LAS algorithm search space was limited to contiguous submatrices of at most 2Mb-by-2Mb in size, i.e., 8-by-8 submatrices (at 250kb resolution). For each chromosome pair, each iteration of the search procedure was initialized at a random contiguous k-by-l submatrix in the log-transformed and standardized interchromosomal Hi-C map, such that the initial submatrix contained at least one active adhesome gene or adhesome TF gene. There are 3,990,387,344 submatrices of size at most 8-by-8 in the collection of interchromosomal Hi-C maps corresponding to all pairs of autosomes. To control the Family-Wise Error Rate (FWER) of the procedure at 0.0001, we used a Bonferroni correction and identified submatrices whose LAS *p*-value was below 0.0001/3,990,387,344 = 2.5e-14. To be even more conservative, we applied a *p*-value cutoff of 1e-15, as shown in Supplementary Fig. 24B (*SI Appendix*).

The resulting distribution of average Hi-C contacts of the submatrices selected by the LAS algorithm is displayed in Supplementary Fig. 24C (*SI Appendix*), where we also show a Hi-C cutoff at 2 (dashed red vertical line). We used this threshold to discard submatrices detected by LAS whose average Hi-C contact value was low. For example, the LAS submatrices identified for the chromosome pair 19-20 before and after applying the Hi-C threshold are reported in Figs. S9D-E. Thresholding based on the average Hi-C value alone is slightly different from thresholding based on the *p*-value, because the *p*-value depends on both the average Hi-C value and the dimension of the submatrix.

### Gene ontology (GO) enrichment analysis

GO enrichment analyses were performed using the GSEApy Enrichr ^64–66^ module with the GO_Biological_Process_2018 gene set in humans.

## Data availability

All data used in this study are publicly available. General genome data was retrieved from the UCSC Genome Browser ^40,42^ Table Browser. Adhesome data was released online (http://adhesome.org/) by ^36,37^. Data on IMR-90 and GM12878 gene expression were obtained from the GEO (references GSE16256 and GSE90276, respectively); data on regulatory marks were obtained from ^39^, where it was gathered from public sources and deposited on GitHub (https://github.com/shwhalen/targetfinder). Hi-C data from ^20^ was downloaded from the GEO (reference GSE63525). TF-target relationships were retrieved from the hTFtarget portal (http://bioinfo.life.hust.edu.cn/hTFtarget#!/) developed by ^16^. The DNA-MERFISH data used for validation was obtained from ^22^ who deposited it on Zenodo. Finally, the PPI data used to analyze physical interactions of adhesome loci was retrieved from the STRING v11 interactome ^57^ (https://string-db.org/). See ^41^ for more details on data availability and processing.

## Code availability

We used Python 3.7 and relied on open-source Python libraries for the code in this work. In particular, we used scikit-learn 0.24.2, NetworkX 2.5, SciPy 1.7.0, pybedtools 0.8.1, Requests 2.22.0 and GSEApy 0.10.3. Our code is available on GitHub (https://github.com/uhlerlab/adhesome_clustering/).

## Acknowledgments

LVC was supported by a Bok Center Fellowship at Harvard University. CU was partially supported by the National Center for Complementary and Integrative Health/National Institutes of Health (Grant 1DP2AT012345), the Office of Naval Research (Grant N00014-22-1-2116), the National Science Foundation (Grant DMS-1651995), the MIT-IBM Watson AI Lab, the MIT J-Clinic for Machine Learning and Health, the Eric and Wendy Schmidt Center at the Broad Institute and a Simons Investigator Award. GVS was partially supported by the Swiss National Foundation award (310030_208046) and he also thanks ETH Zurich and the Paul Scherrer Institute for funding.

## Author Contributions

Conceptualization: LVC, CU, GVS; Methodology: LVC, CU, GVS; Software: LVC; Formal analysis: LVC; Writing (original draft): LVC, CU, GVS.

## Competing Interest Statement

The authors declare no conflict of interest.

## Supporting Information for

**Supplementary Fig. 1:**
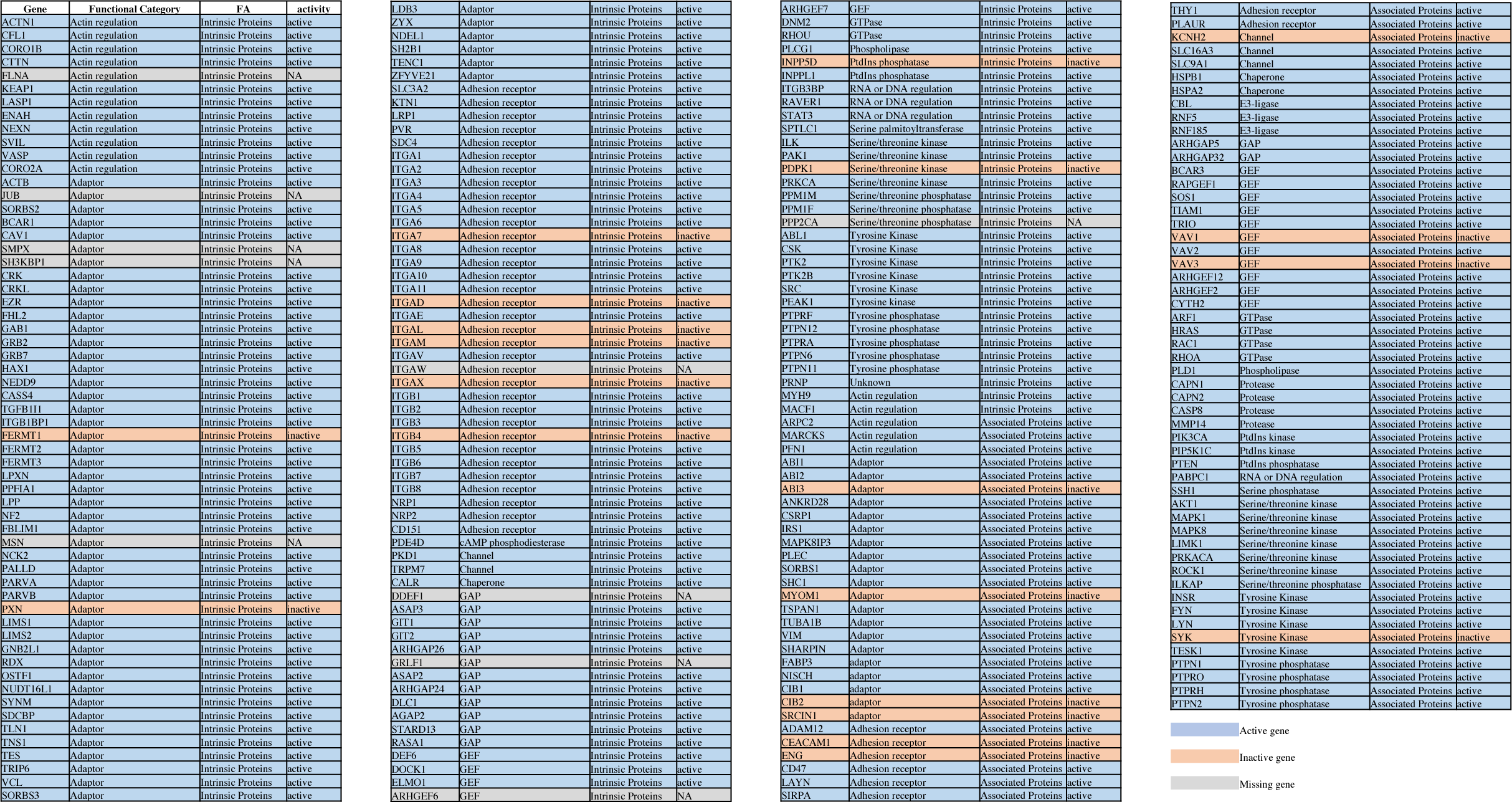
List of adhesome genes and their activity. Intrinsic and associated adhesome genes from ^1^ colored by activity level.

**Supplementary Fig. 2:**
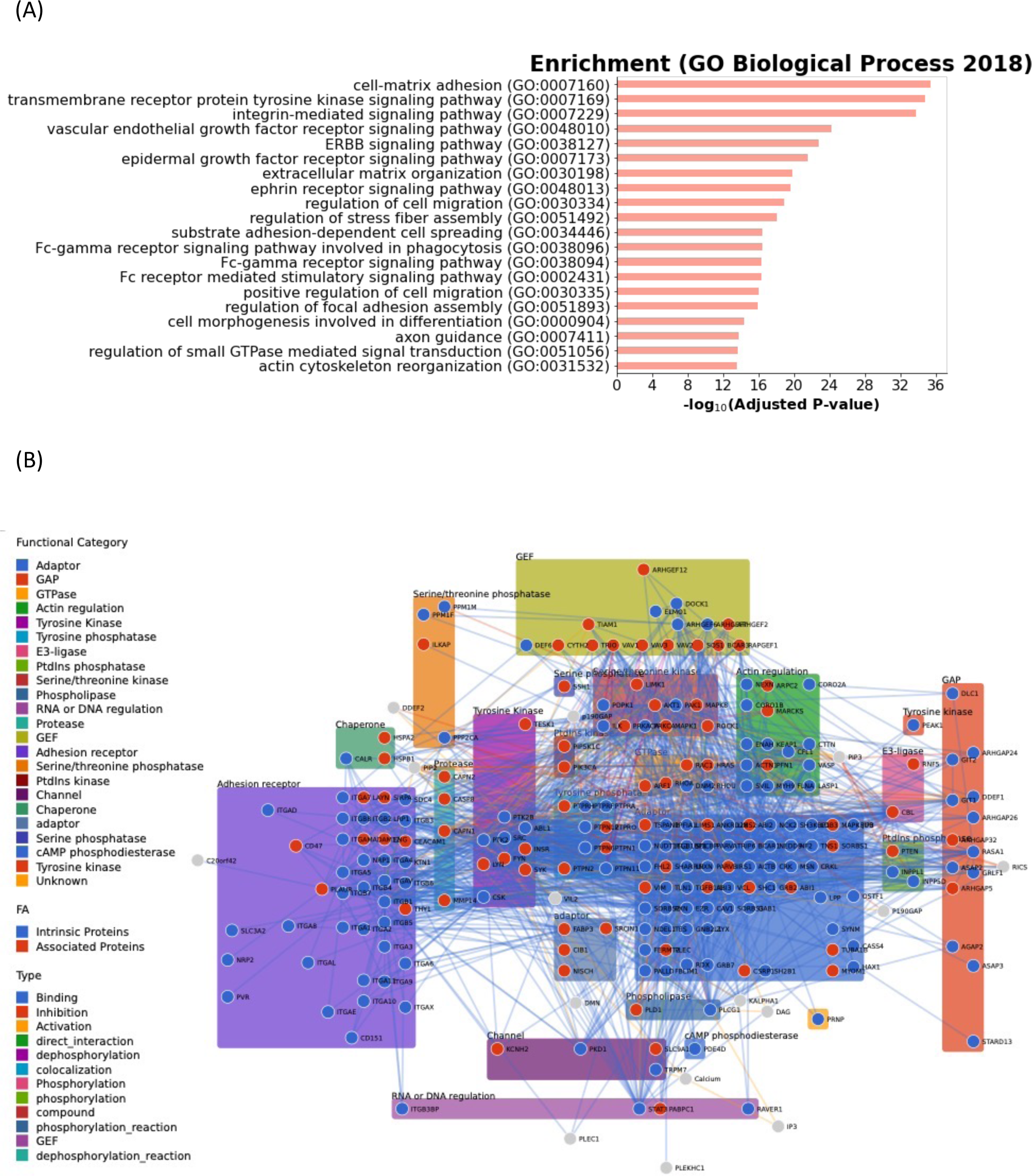
Function and structure of the adhesome. (**A**) Gene Set Enrichment Analysis of intrinsic and associated adhesome components using the 2018 Gene Ontology database for biological processes. (**B**) Network of interactions among adhesome proteins. Nodes are colored in blue if they are intrinsic adhesome proteins, and in red if they are associated adhesome proteins. Nodes are grouped by functional categories. Links between nodes are colored based on the type of interaction.

**Supplementary Fig. 3:**
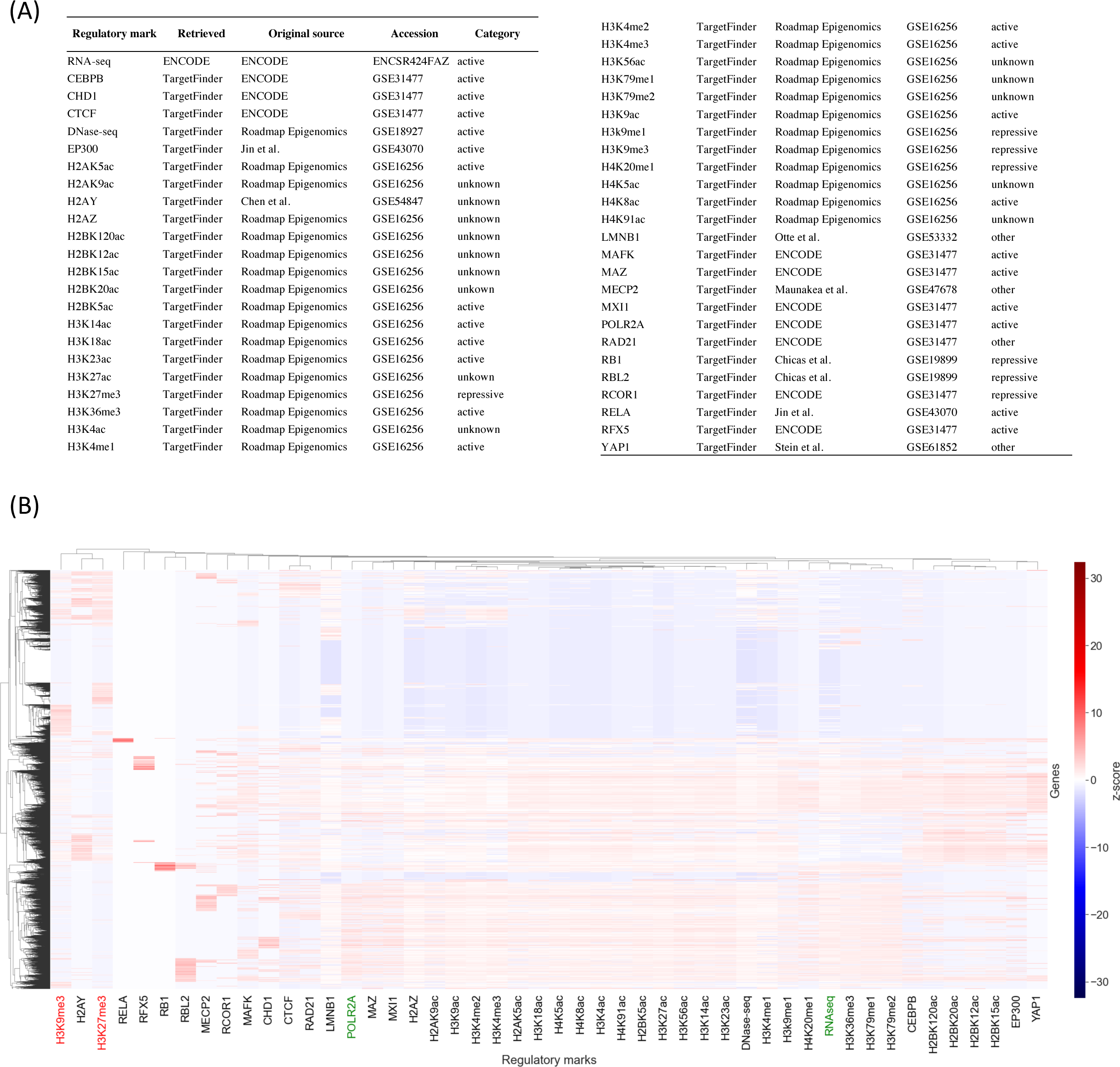
Genome-wide regulatory marks in IMR-90 cells. (**A**) Table describing the 48 regulatory features used in this study, including bulk RNA-seq and 47 cistromic marks (TF and histone ChIP-seq, DNAse-seq). The table also reports the source and accession ID of each data set. Likely annotations on the regulatory activity of each mark are provided when available in the literature (see ^2^ and ^3^). (**B**) Genome-wide heatmap in IMR-90 cells of the 48 regulatory marks in (A). Genes (rows) and regulatory marks (columns) were clustered using hierarchical clustering with average linkage and Pearson correlation metric. The feature matrix was normalized and scaled prior to clustering ^3^. The labels of two typical active marks (RNA-seq, POLR2A ChIP-seq) are colored in green and the labels of two typical repressive marks (H3K9me3, H3K27me3 ChIP-seq) are colored in red.

**Supplementary Fig. 4:**
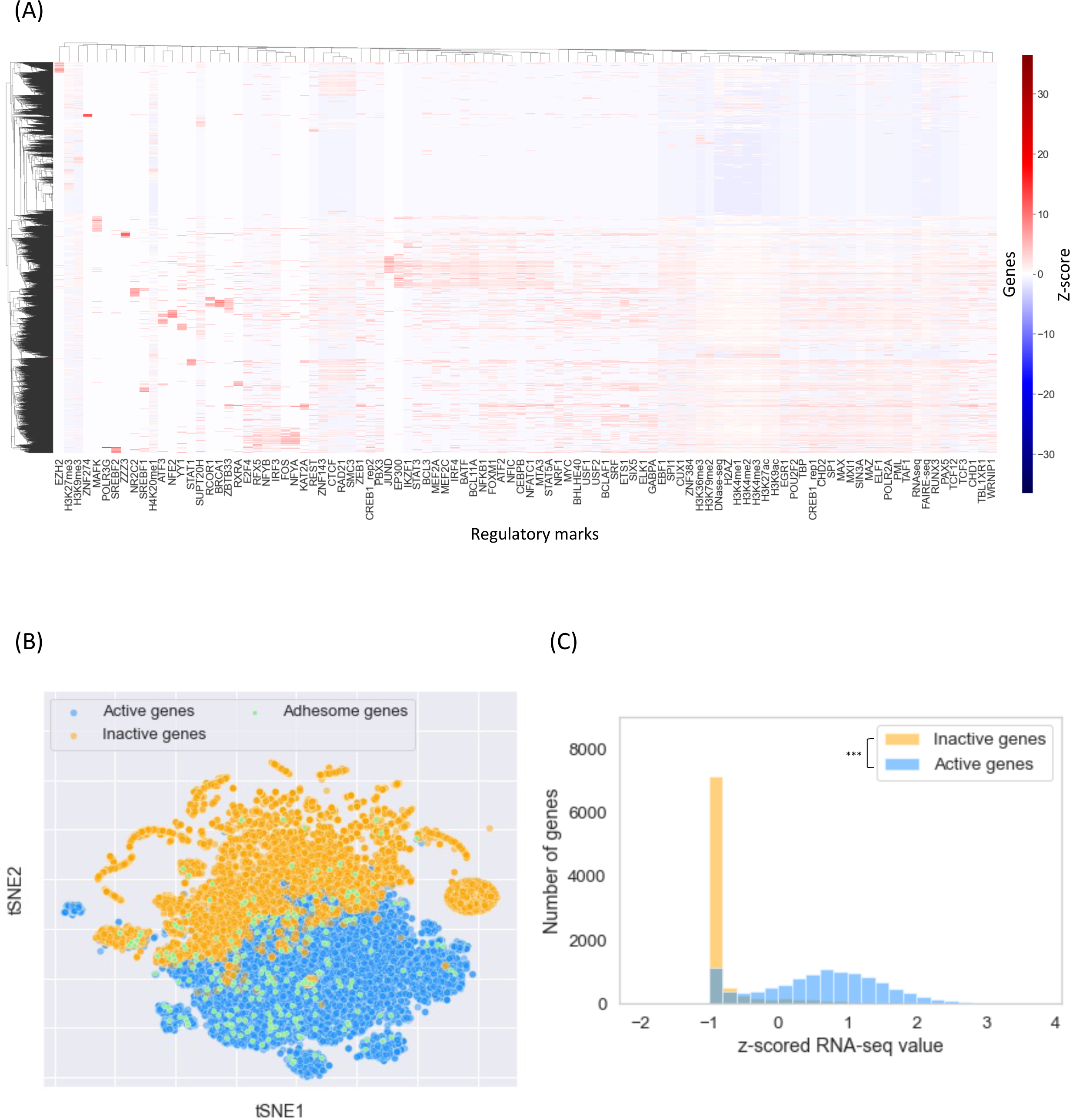
Activity of adhesome genes in GM12878. (**A**) Heatmap of 100 regulatory marks used to determine the activity of genes in GM12878 ^3^. Genes (rows) and regulatory marks (columns) were clustered using hierarchical clustering with average linkage and Pearson correlation metric. The feature matrix was normalized and scaled prior to clustering ^3^. (**B**) tSNE representation of all 25,959 genes based on the 100 GM12878 regulatory marks. Orange dots correspond to inactive genes (9,107 genes, as determined by the clustering in (A)), blue dots correspond to active genes (10,984 genes), and green dots correspond to adhesome genes. Out of the 222 adhesome genes studied (green dots), 172 are active (in contrast to 202 in IMR-90 cells). (**C**) Histogram of bulk RNA-seq expression in GM12878 for all active genes (blue) and all inactive genes (orange). Active genes have a significantly higher expression than inactive genes (*p*-value < 3e-308, Wilcoxon rank-sums test).

**Supplementary Fig. 5:**
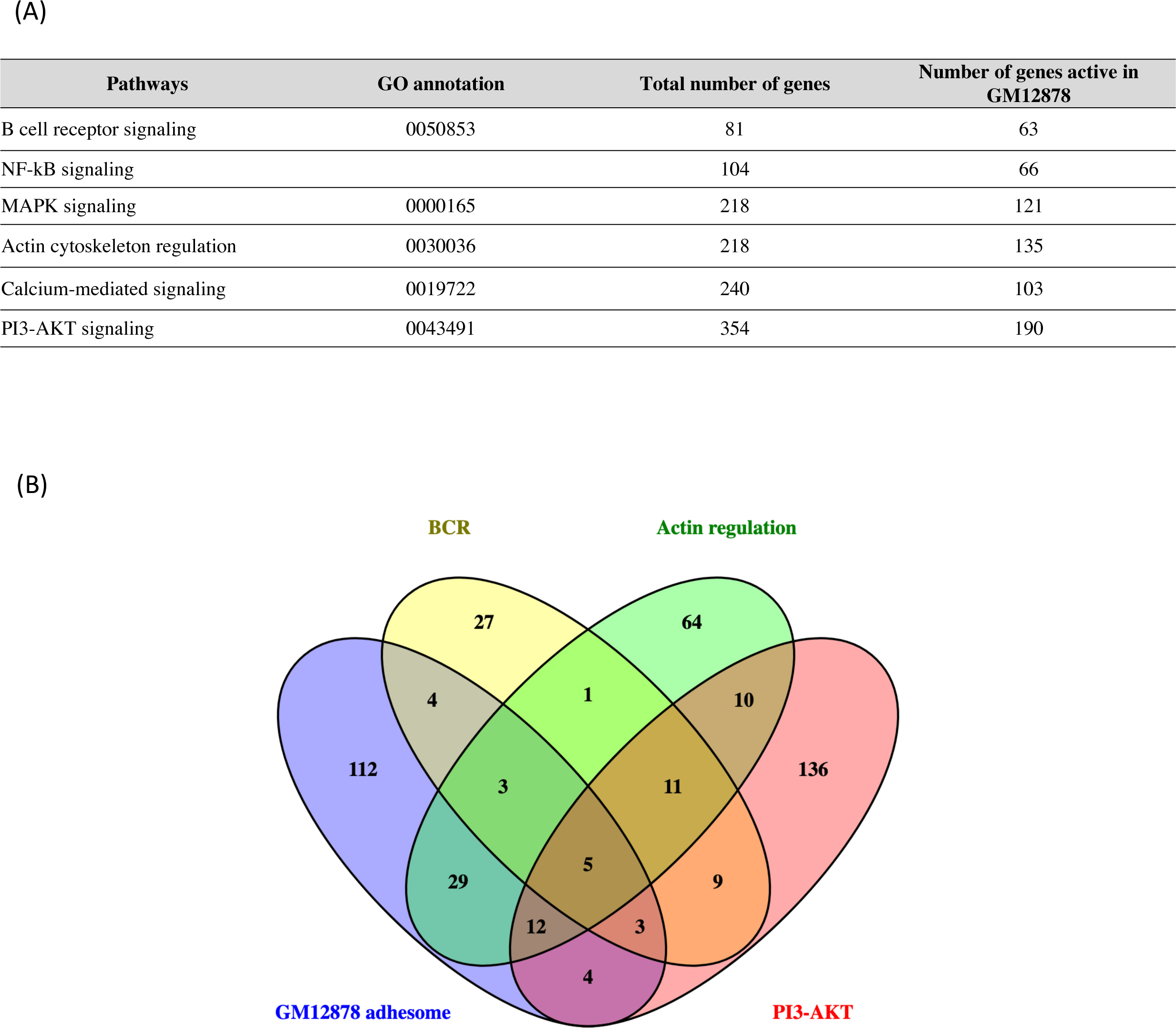
Active adhesome genes intersect with B Cell Receptor signaling pathways in GM12878. (**A**) Table of signaling pathways related to the B Cell Receptor (BCR) signaling pathway. We intersect genes in each pathway with the active genes in GM12878 (as determined in Supplementary Fig. 4) to obtain the corresponding number of active pathway genes. (**B**) Venn diagram of four gene lists: active adhesome genes in GM12878 (blue), active BCR signaling genes (yellow), active actin cytoskeleton regulation genes (green), and active PI3- AKT signaling pathway genes (red). The Venn diagram was drawn using Venny ^4^.

**Supplementary Fig. 6:**
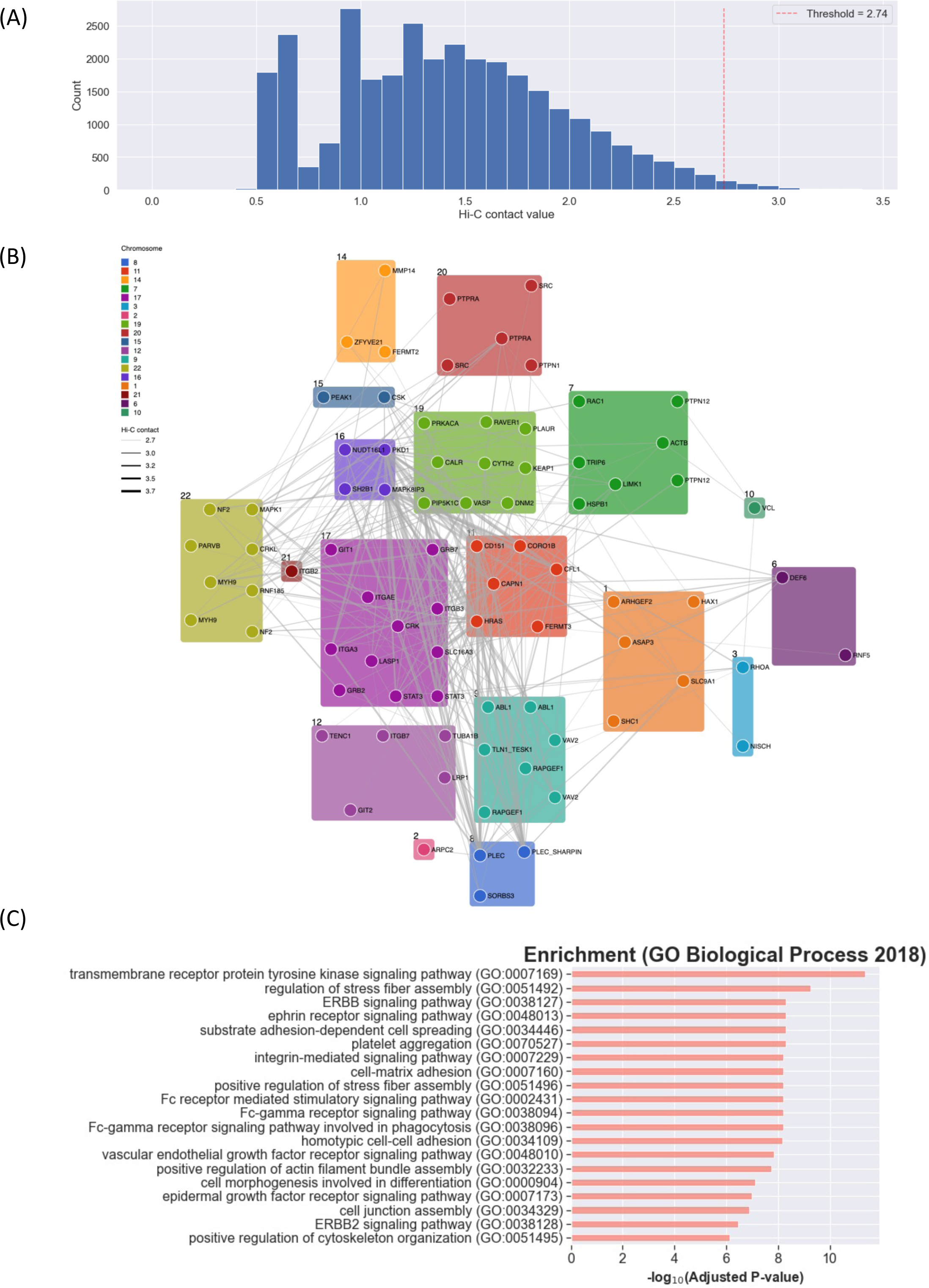
Active adhesome genes with high Hi-C contacts. (**A**) Distribution of processed interchromosomal Hi-C contact values among active adhesome gene loci. A lower threshold materialized by the dashed red vertical line at 2.74 (99th percentile) is applied to select the strongest loci interactions, which are shown in the network in B. (**B**) Subnetwork of active adhesome gene loci, colored by chromosomal location and labeled with the active adhesome genes they harbor. Only non-isolated nodes engaging in high Hi-C contacts (Hi-C > 2.74, see A) are shown in the plot; edges are sized according to the corresponding Hi-C contact value. The network contains 82 active adhesome loci (74 adhesome genes) and 321 edges (average degree = 7.8). (**C**) Gene set enrichment analysis of active adhesome genes present in the subnetwork in B using the GO Biological Process 2018 gene sets. Only GO terms with FDR-corrected enrichment *p*-value smaller than 0.0001 are reported.

**Supplementary Fig. 7:**
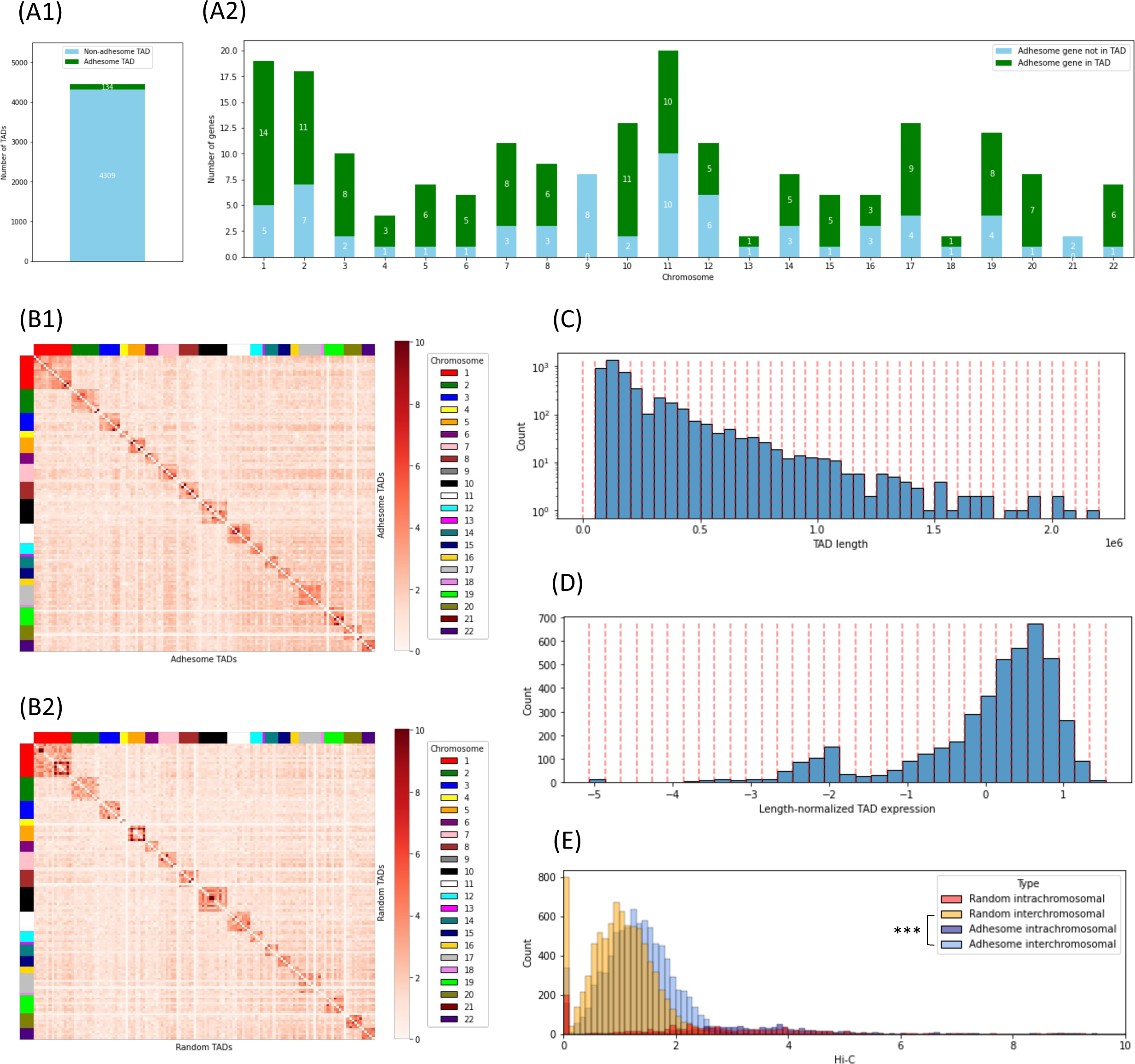
TAD-based interchromosomal proximity analysis. (**A**) 134 TADs out of 4,443 TADs called by Arrowhead (at resolution 5 kb with Knight-Ruiz normalization) contain at least one active adhesome gene (A1). The number of active adhesome genes either in TADs or not in TADs per chromosome is shown in A2. (**B**) Hi-C contact map between active adhesome TADs (C1) and random active non-adhesome TADs (C2). Each row/column corresponds to a TAD. The contact value between two TADs is obtained by averaging the Hi-C contact values associated with these TADs. TADs are grouped by chromosomal location, as shown by the side color bars. The selection of random active adhesome TADs is explained in D and E. (**C**) Distribution of TAD length for all 4,443 TADs called by Arrowhead at resolution 5 kb with Knight-Ruiz normalization. TADs are stratified into buckets of length 50 kb. These buckets are used to sample background TADs on a per-chromosome basis. (**D**) Distribution of length-normalized TAD gene expression for all 4,443 TADs called by Arrowhead at resolution 5 kb with Knight-Ruiz normalization. TADs are stratified into buckets of size 0.2. These buckets are used to sample background TADs on a per-chromosome basis. (**E**) Distribution of interchromosomal and intrachromosomal contacts between active adhesome TADs and between random TADs. Random TADs were selected such that they are similar in size to adhesome TADs, they are similarly distributed over the chromosomes, and they have a similar expression level. The distribution of interchromosomal contacts between active adhesome TADs significantly dominates the distribution of interchromosomal contacts between random TADs (Wilcoxon Rank-Sum test, *p*-value < 5e-124).

**Supplementary Fig. 8:**
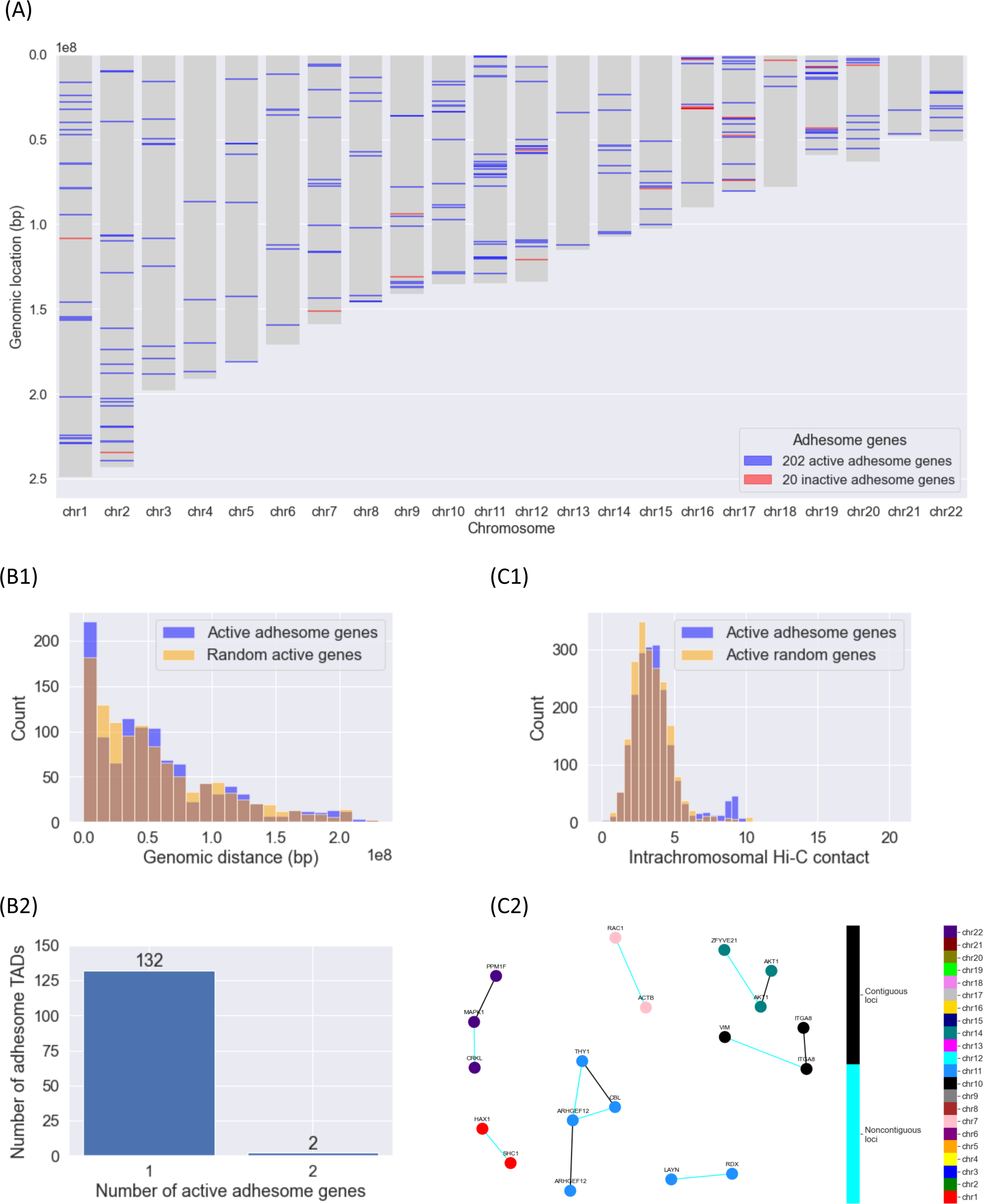
Intrachromosomal proximity analysis. (**A**) Localization of active and inactive adhesome genes on the chromosomes (using the hg19 reference genome). (**B**) B1: Distribution of 1D genomic distances (in bp) between active adhesome genes and between a set of random active genes located on the same chromosomes; the distance between 2 genes (located on the same chromosome) is defined as the minimum distance between |start2-end1| and |start1-end2|. B2: Number of active adhesome genes per active adhesome TAD, using TADs called by Arrowhead at resolution 5 kb with Knight-Ruiz normalization. (**C**) C1: Distribution of intrachromosomal Hi-C contacts among active adhesome gene loci and random active non-adhesome loci located on the same chromosomes. C2: Subnetwork of active adhesome loci whose intrachromosomal Hi-C contact values are above 7 in C1; only non-degenerate connected components are shown. Adhesome gene loci are colored by their chromosomal location and labeled with the adhesome genes they harbor. Contiguous loci are linked by black edges, while non-contiguous loci are linked by cyan edges.

**Supplementary Fig. 9:**
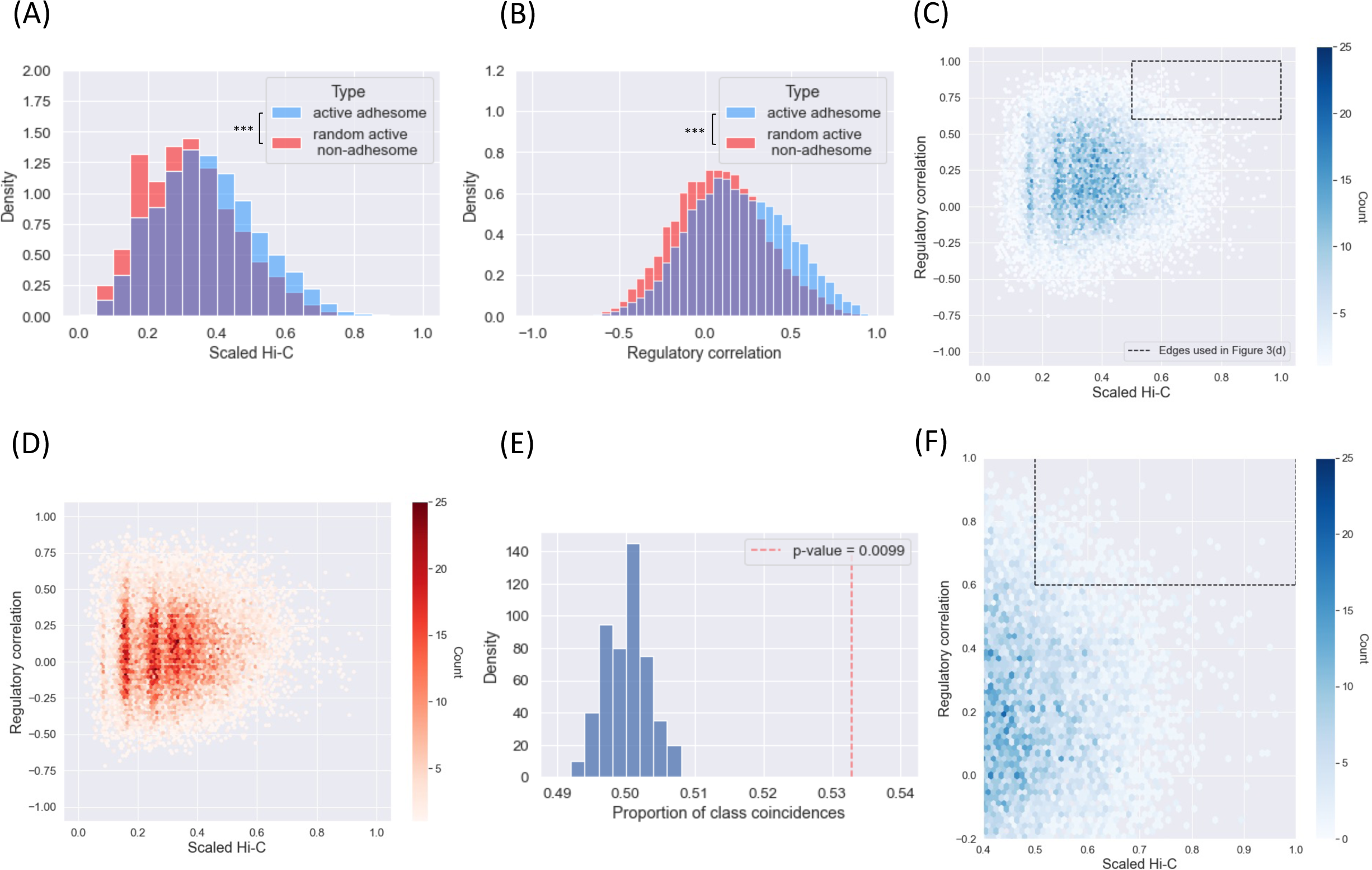
Joint distribution of scaled Hi-C contact values and regulatory correlation among active adhesome genes in IMR-90 cells. (**A**) Histogram of scaled Hi-C contact values between active adhesome genes (blue) and a random set of active non-adhesome genes (red). There are significantly more contacts between adhesome genes compared to random non-adhesome genes (Wilcoxon Rank-Sums test, *p*-value = 1e-85). (**B**) Histogram of regulatory Pearson correlation (based on the 48 regulatory marks of Supplementary Fig. 3A) between active adhesome genes (blue) and a random set of active non-adhesome genes (red). There is significantly more coregulation among adhesome genes compared to random non-adhesome genes (Wilcoxon Rank-Sums test, *p*-value < 3e-308). (**C**) 2D histogram of scaled Hi-C contact values and Pearson regulatory correlation for active adhesome genes. The dashed box shows the thresholds used to build the network in Fig. 3D. (**D**) 2D histogram of scaled Hi-C contact values and Pearson regulatory correlation for random active non-adhesome genes. (**E**) Simulated null distribution for the permutation test used to assess the difference between the joint distribution of scaled Hi-C contact values and Pearson regulatory correlation for the two gene groups of interest: active adhesome genes (blue, see (C)) and the randomly selected set of active non-adhesome genes (red, see (D)). The test is a two-sample test based on the number of nearest neighbor type co-incidences ^3^. The red vertical dashed line corresponds to the actual value of the statistic. (**F**) Zoom into the region around the dashed box of (C).

**Supplementary Fig. 10:**
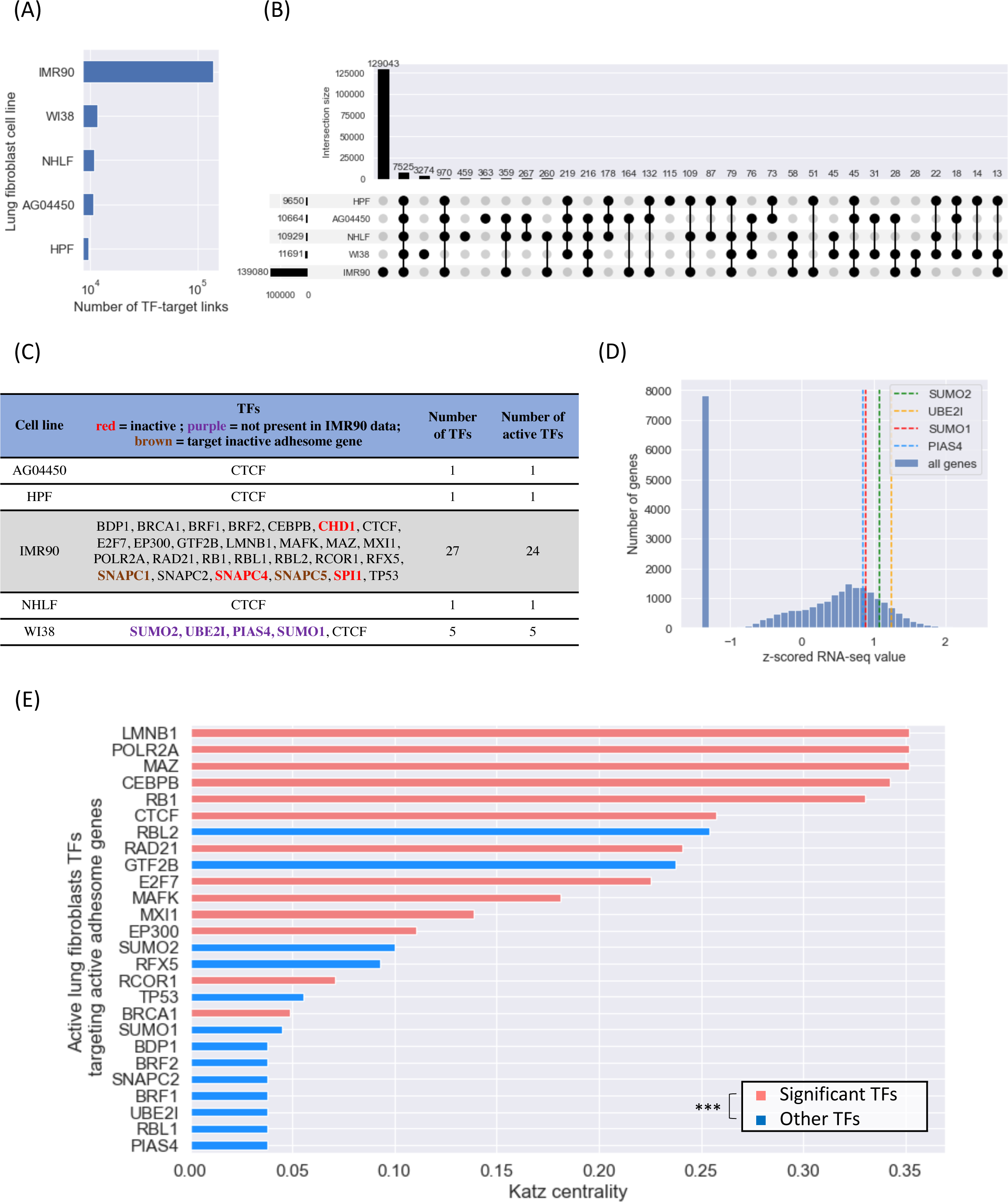
TF-target data in normal human lung fibroblast cell lines. (**A**) Bar plot of the number of TF-target links from the hTFtarget data set in five human lung fibroblast cell lines (AG04450, HPF, NHLF, WI38, IMR-90). Only targets that are active in IMR-90 are considered. (**B**) Set overlap plot for the TF-target relationships in cell lines AG04450, HPF, NHLF, WI38, and IMR-90. Only targets that are active in IMR-90 are considered. While most identified TF-target interactions come from the IMR-90 data, other human lung fibroblast cell lines contribute a few additional TF-target interactions that are not present in the IMR-90 data. (**C**) Table of TFs present for all normal human lung fibroblast cell lines. Red font indicates inactive genes, purple font indicates TFs that are not present in IMR-90, and brown font indicates TFs that target inactive adhesome genes. (**D**) Histogram of z-scored RNA-seq expression for all genes in IMR-90. The expression of SUMO2, SUMO1, UBE2I and PIAS4 (transcription factors found in WI38 only) are represented with colored vertical dashed lines. (**E**) Bar plot of Katz centrality of all active lung fibroblast TFs in the network of Fig. 4E. The parameter alpha of Katz centrality was chosen to be 1/s-0.01, where s is the largest eigenvalue (in magnitude) of the adjacency matrix of the network in Fig. 4E. Bars are colored in red if they correspond to significant TFs from Fig. 4D, otherwise they are colored in blue. There is a significant enrichment of significant TFs at the top of the list (XL-minimal hypergeometric test ^5^, *p*-value < 6e-4 and optimal cutoff value at 17, which corresponds to BRCA1). The parameters X and L were chosen to be 10% of the number of significant TFs and 150% of the number of significant TFs, respectively.

**Supplementary Fig. 11:**
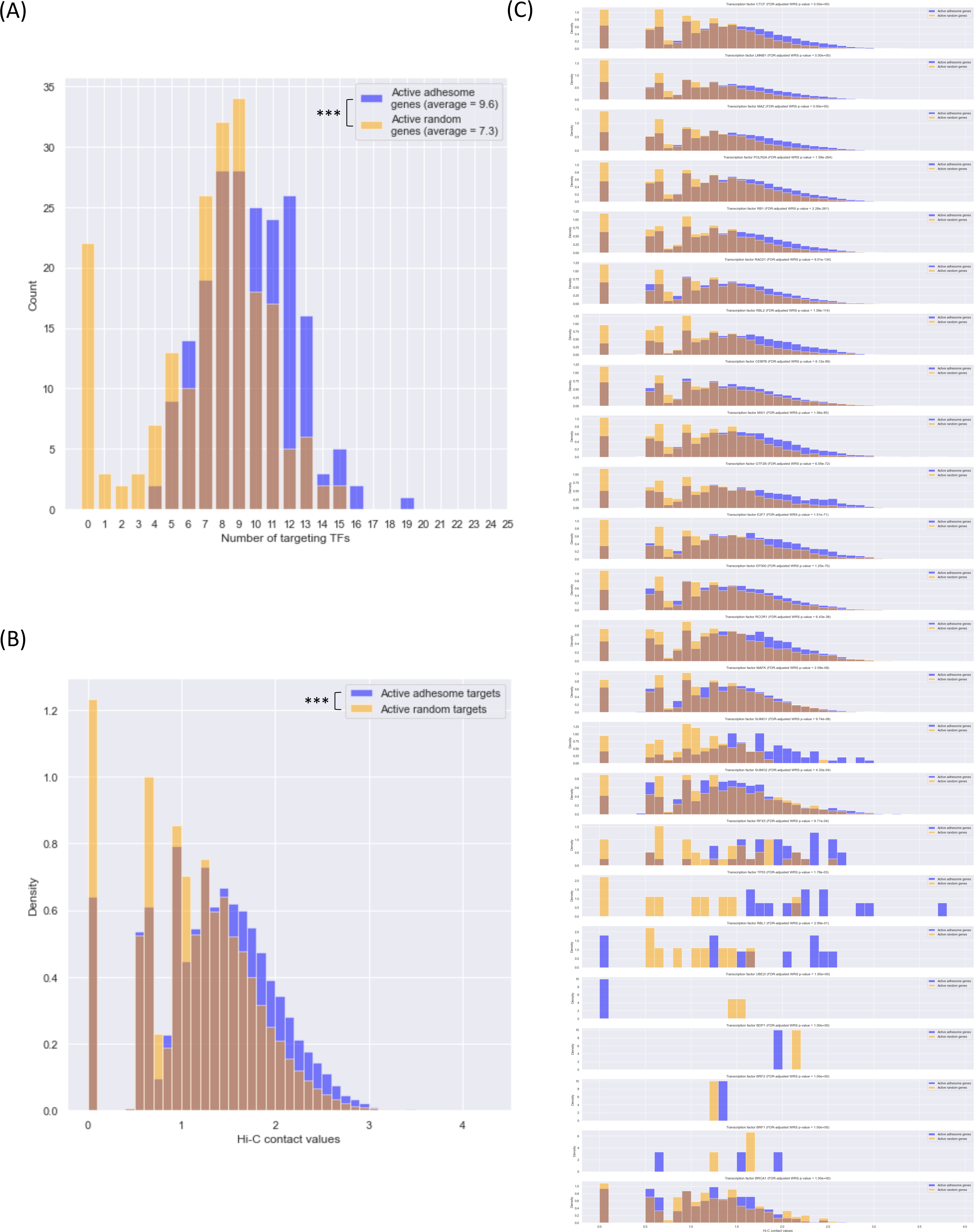
Spatial proximity analysis of transcription factor targets. (**A**) Histograms of the number of TFs targeting active adhesome genes and the number of TFs targeting a set of random active non-adhesome genes (Wilcoxon Rank-Sum *p*-value < 2e-10). (**B**) Histograms of interchromosomal Hi-C contacts among active adhesome TF targets and random active non-adhesome non-TF target genes located on the same chromosomes (Wilcoxon Rank-Sum test, *p*-value < 3e-308). (**C**) Histograms of interchromosomal Hi-C contacts among active adhesome TF targets and among random active non-adhesome non-TF target genes located on the same chromosomes, separately for each TF. The plots are organized in ascending order of Wilcoxon Rank-Sum *p*-value.

**Supplementary Fig. 12:**
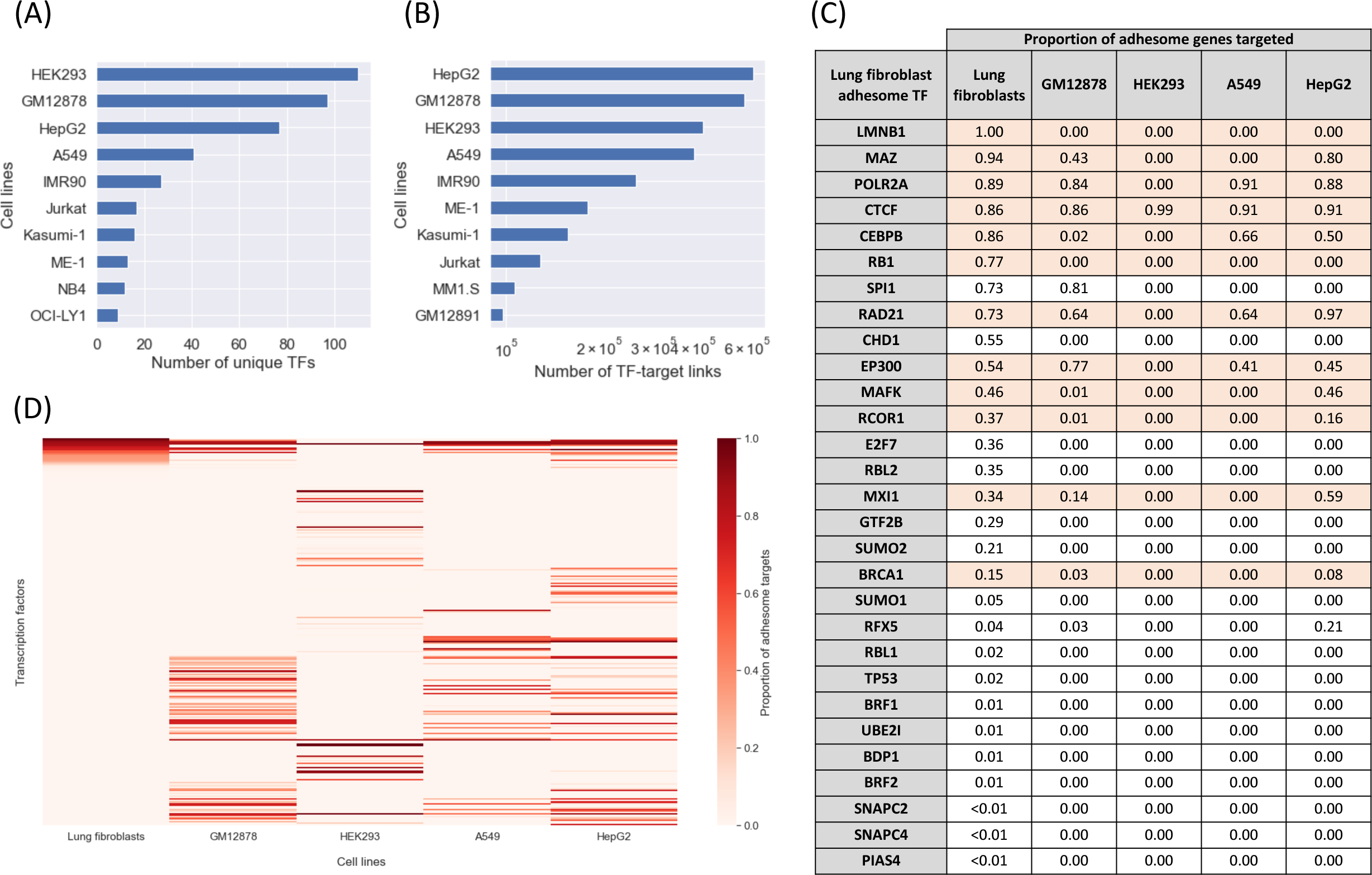
The identified active human lung fibroblast TFs target fewer adhesome genes in other cell lines and are quite specific to human lung fibroblasts. (**A**) Number of unique TFs present in the hTFtarget data set for different adherent and non-adherent cell lines. (**B**) Number of unique TF-target links present in the hTFtarget data set for different cell lines. The cell lines HepG2, GM12878, HEK293, A549 and normal lung fibroblasts (combination of IMR-90, WI38, AG04450, HPF and NHLF) have the highest amount of data. (**C**) Table showing the proportion of adhesome genes targeted by selected lung fibroblast TFs in different cell lines, including lung fibroblasts, HepG2, GM12878, HEK293, and A549. Rows shaded in red correspond to the 13 significant TFs identified in lung fibroblasts. (**D**) Heatmap of the proportion of adhesome genes targeted by TFs in different cell lines compared to the proportion of adhesome genes targeted in normal lung fibroblasts. These include cell lines with adherence properties (A549, HepG2, HEK293) and a non-adherent cell line (GM12878). TFs that target (active and/or inactive) adhesome genes in IMR-90 are in the top 29 rows. The remaining TFs do not target adhesome genes in IMR-90 but target at least one adhesome gene in other cell lines.

**Supplementary Fig. 13:**
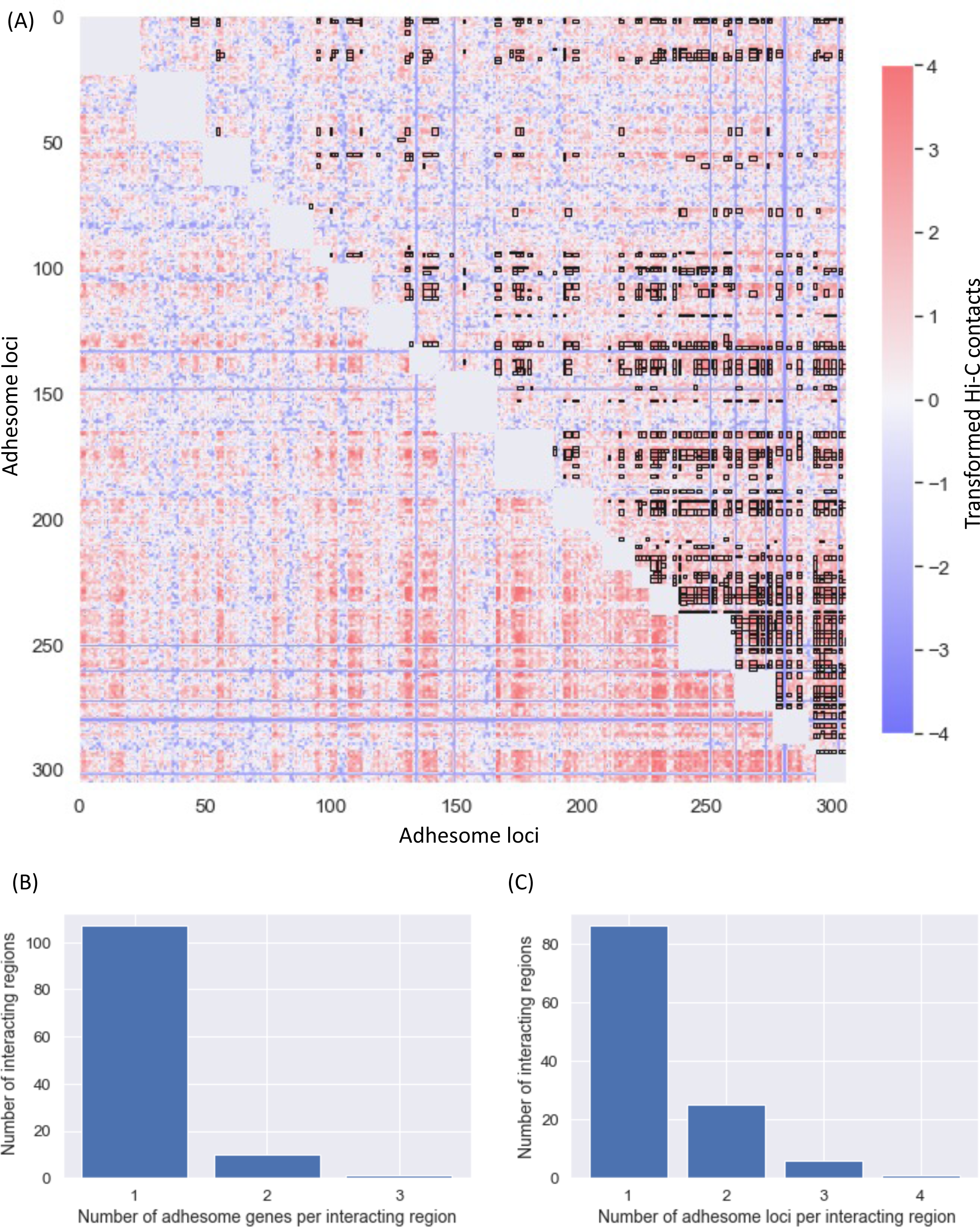
Projection of LAS submatrices onto adhesome loci. (**A**) Heatmap of transformed Hi-C contact frequencies among active adhesome and adhesome TF loci (frequencies were log-transformed and centerdized using the mean and standard deviation of contact values across the entire genome). Significant submatrices identified by the LAS algorithm and whose average Hi-C value is higher than 2 are shown as black boxes. This heatmap is obtained by combining LAS interacting regions containing adhesome genes from all interchromosomal Hi-C contact matrices (such as, for example, the matrix in Fig. 5A). (**B**) Bar plot of the number of adhesome genes per adhesome interacting region. Adhesome LAS interacting regions contain at most 3 adhesome genes. (**C**) Bar plot of the number of adhesome loci per adhesome interacting region. Adhesome LAS interacting regions contain at most 4 adhesome loci. To relate this to (B), note that an adhesome gene may span multiple 250 kb loci.

**Supplementary Fig. 14:**
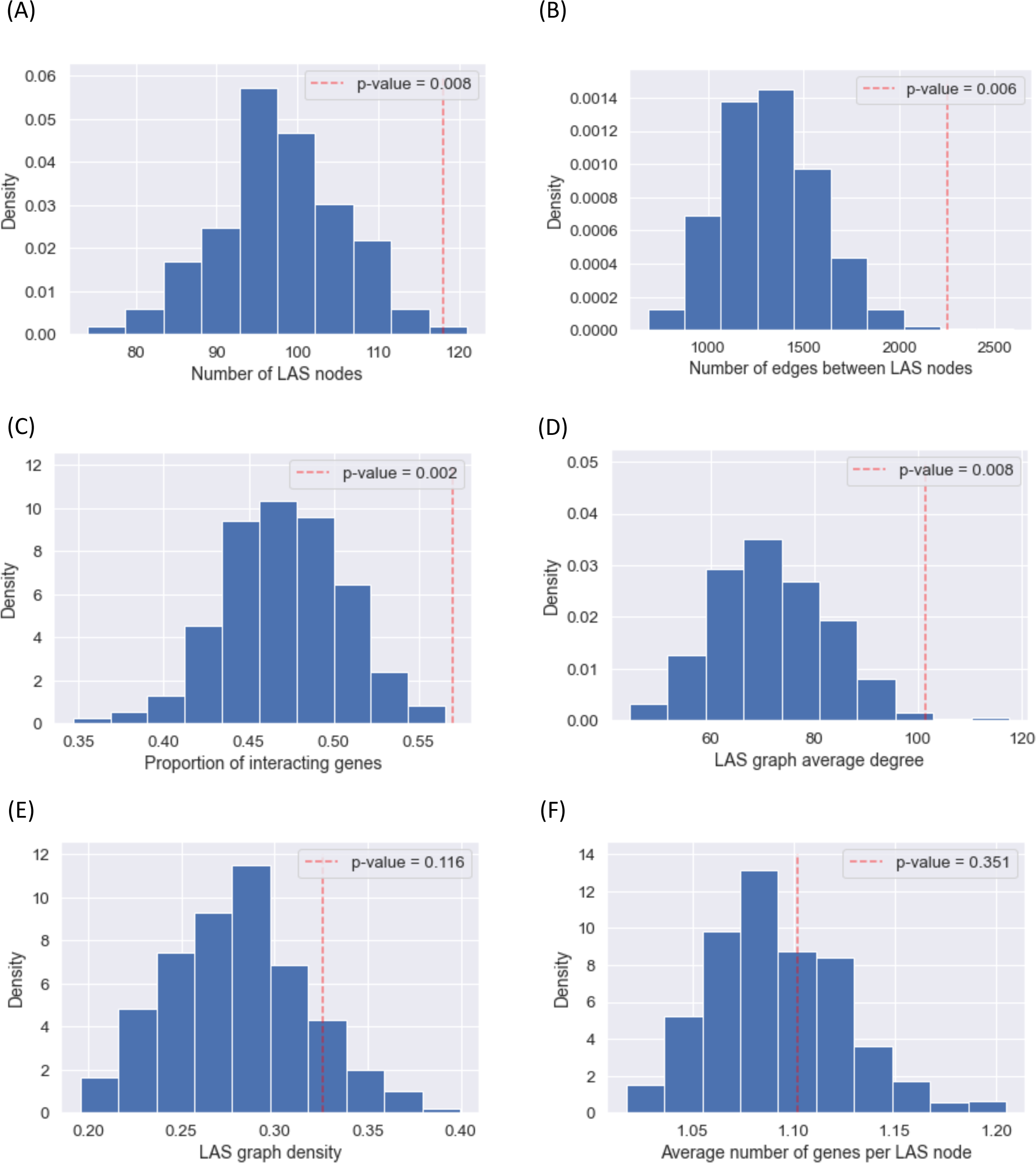
Statistical significance of the LAS adhesome network. (**A**) Histogram of the number of LAS nodes containing at least one gene from a collection of 500 random gene sets. Each random gene set is sampled uniformly without replacement among all active non-adhesome genes. The number of nodes containing active adhesome and adhesome TF genes is shown with a red dashed vertical line, corresponding to a *p*-value of 0.008. This shows that a significantly high number of nodes contain adhesome genes under the 1% significance level. (**B**) Histogram of the number of interactions among LAS nodes containing at least one gene from the collection of 500 random gene sets in (A). The number of interactions among adhesome LAS nodes is shown with a red dashed vertical line, corresponding to a *p*-value of 0.006. This shows that there is a significantly high number of interactions between adhesome LAS nodes at the 1% significance level. (**C**) Histogram of the proportion of genes belonging to LAS nodes from the collection of 500 random gene sets in (A). The proportion of adhesome genes belonging to adhesome LAS nodes is shown with a red dashed vertical line, corresponding to a *p*-value of 0.002. This shows that there is a significantly high proportion of adhesome genes that belong to adhesome LAS nodes at the 1% significance level. (**D**) Histogram of average degree computed on networks of LAS nodes containing genes from the collection of 500 random gene sets in (A). The average degree of the adhesome LAS network is shown with a red dashed vertical line, corresponding to a *p*-value of 0.008. This shows that, on average, the degree of adhesome LAS nodes is significantly higher than the degree of LAS nodes containing random genes at the 1% significance level. (**E**) Histogram of graph density computed on networks of LAS nodes containing genes from the collection of 500 random gene sets in (A). The graph density of the adhesome LAS network is shown with a red dashed vertical line, corresponding to a *p*-value of 0.116. (**F**) Histogram of the average number of genes per LAS node from the collection of 500 random gene sets in (A); only LAS nodes containing at least one gene from the random gene set at hand are considered. The average number of adhesome genes per adhesome LAS node is shown with a red dashed vertical line, corresponding to a *p*-value of 0.351.

**Supplementary Fig. 15:**
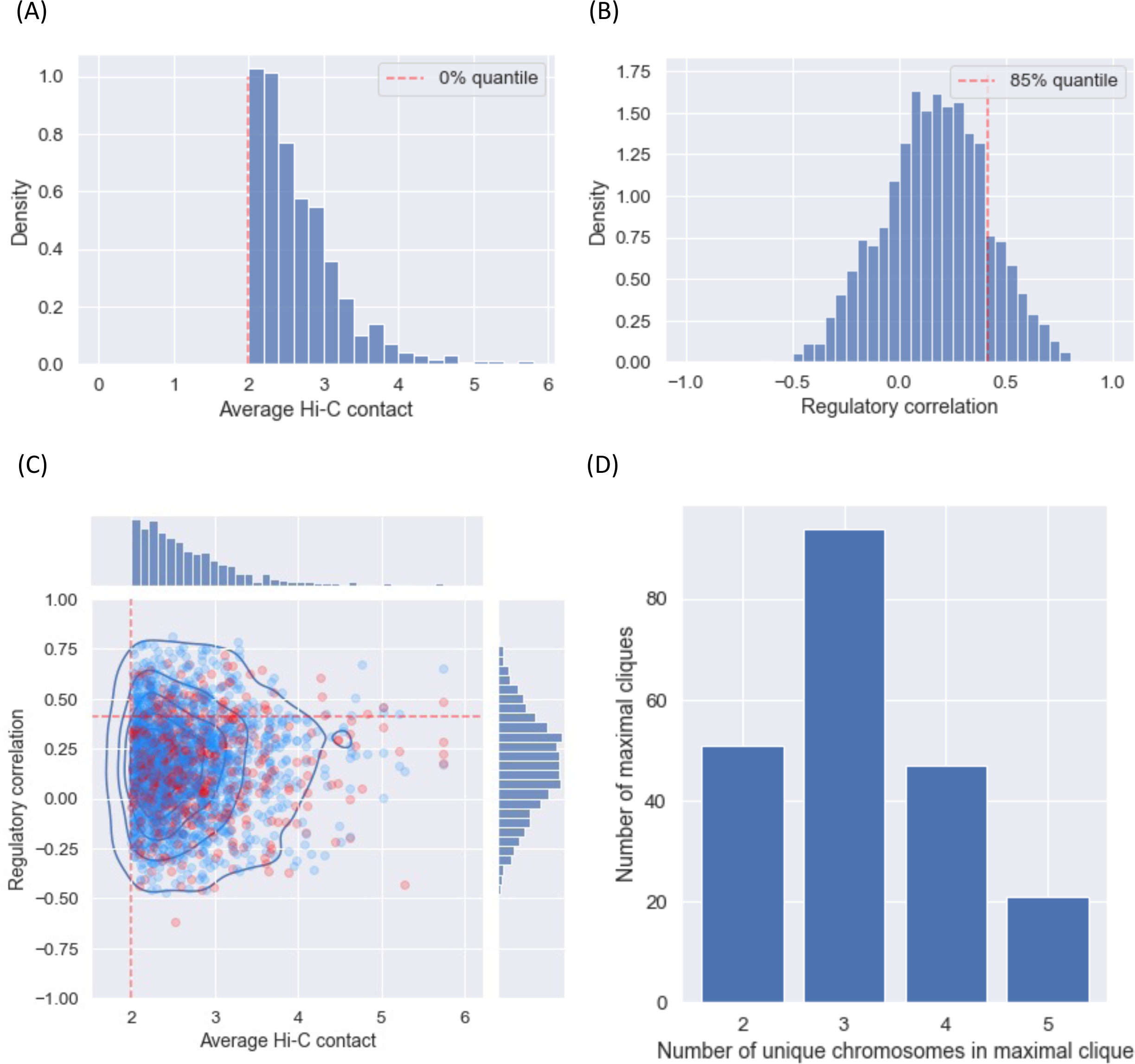
Regulatory correlation of adhesome LAS nodes. (**A**) Histogram of average transformed Hi-C contact values among all adhesome LAS nodes. The red dashed vertical line corresponds to the minimum value, which is equal to 2 (by design from the LAS thresholding we performed in a previous step, see Supplementary Fig. 9C). (**B**) Histogram of average Spearman regulatory correlation among all adhesome LAS nodes. The average Spearman regulatory correlation between two adhesome LAS nodes is computed by taking the average of Spearman regulatory correlations between all tuples of adhesome genes in the two nodes, using the 48 regulatory marks used throughout this article ^3^. The red dashed vertical line corresponds to the 85% quantile, used as a lower regulatory threshold to select interactions among adhesome LAS nodes for the adhesome LAS network of Fig. 5C. (**C**) Scatterplot and 2D contour plot of pairs of adhesome LAS nodes. The location of a given pair (u, v) on the scatterplot is determined by the average Hi-C contact value (x-axis) and the regulatory correlation (y-axis) associated with the edge u–v of the adhesome LAS network. Pairs that involve at least one TF targeting active adhesome genes are shown in red. The red dashed lines correspond to marginal thresholds on the distribution of average Hi-C contact values (vertical, 0% quantile) and regulatory correlations (horizontal, 85% quantile). (**D**) Bar plot of the number of unique chromosomes per maximal clique in the adhesome LAS network obtained after dropping interactions whose average Spearman regulatory correlation is below the threshold in (B). Maximal cliques contain at most 5 unique chromosomes, which is consistent with the hypothesis that no more than 5 chromosomes can physically interact at a given location.

**Supplementary Fig. 16:**
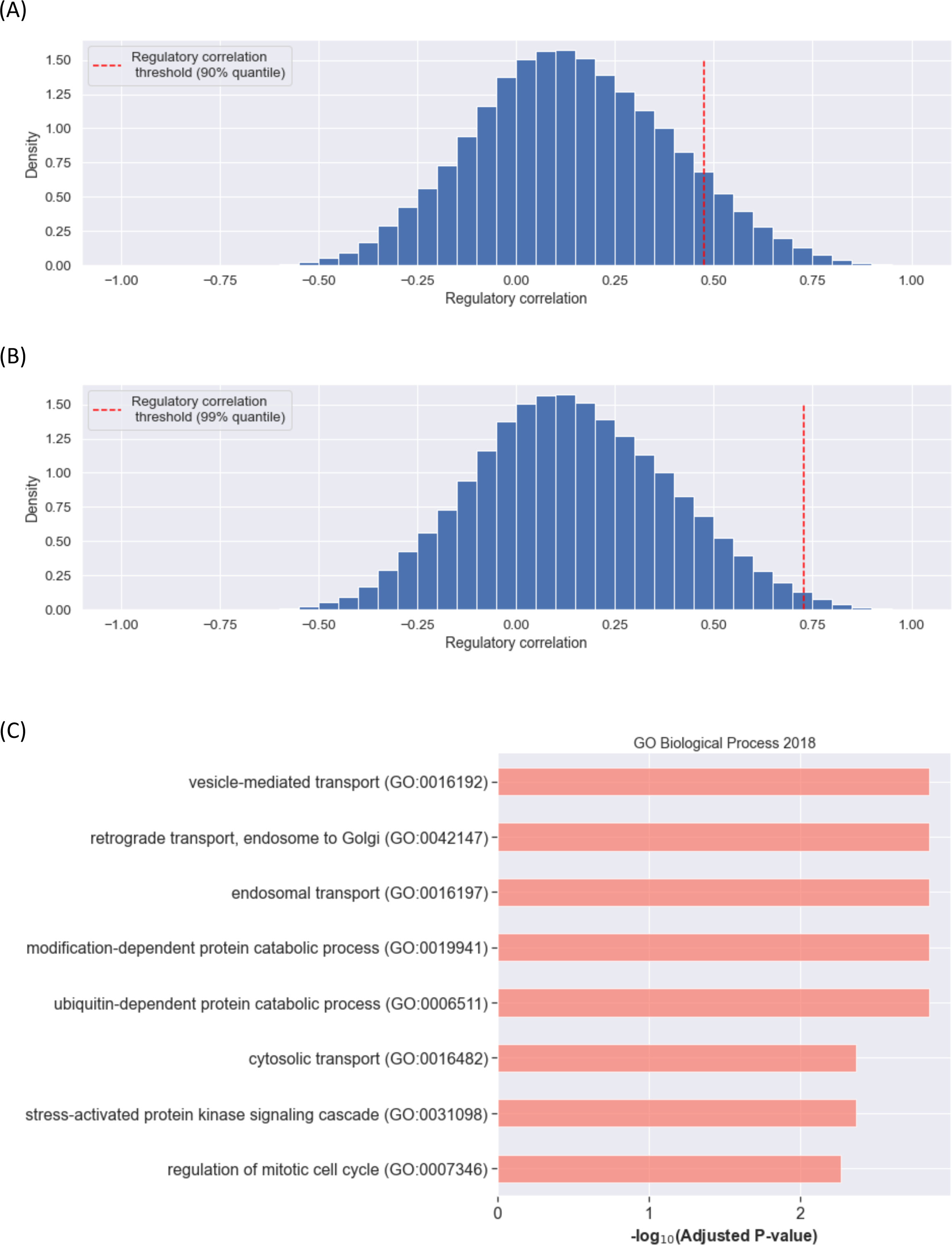
Gene set enrichment analysis of active non-adhesome genes interacting with active adhesome genes. (**A**) Histogram of Spearman regulatory correlation between (i) active non-adhesome genes in adhesome interacting regions and (ii) the active adhesome genes with which these genes interact interchromosomally. Regulatory correlation is computed from the 48 regulatory marks used throughout this article ^3^. The red dashed vertical threshold line marks the upper decile of the distribution. (**B**) Same as A, with the red dashed vertical threshold line marking the upper percentile of the distribution. (**C**) Significant Gene Ontology (GO) terms associated to active non-adhesome genes belonging to genomic regions that interact with adhesome interacting regions. Only genes having a strong positive regulatory correlation (upper percentile in B) with the active adhesome genes they interact with (in LAS) are considered. A GO term is reported as significant if its FDR-adjusted *p*-value is lower than 1e-2.

**Supplementary Fig. 17:**
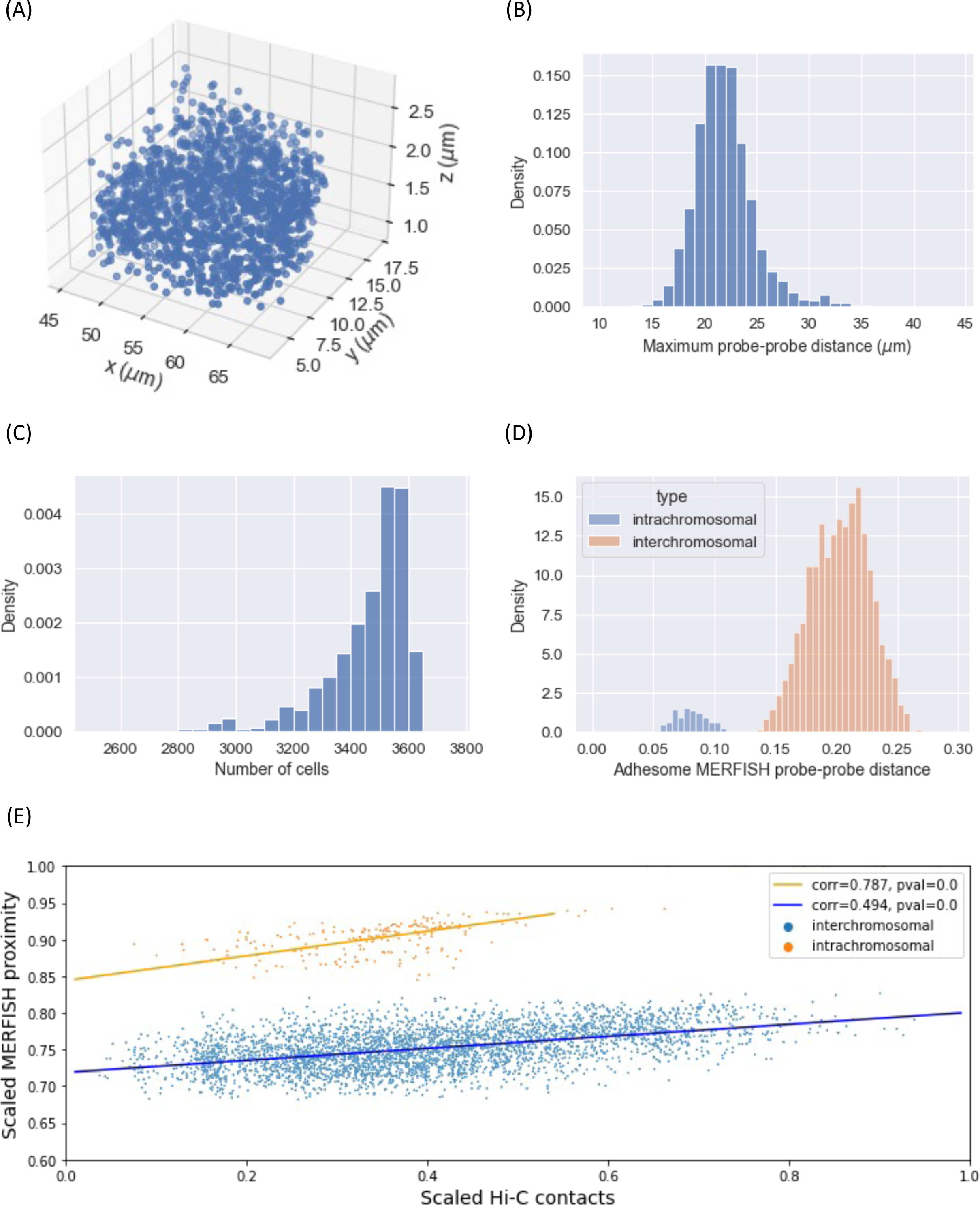
Normalization of Euclidean distances between DNA-MERFISH probes. (**A**) 3D scatterplot of all 959 DNA-MERFISH probes in the nucleus of a given cell (cell ID 1788). The shape of the nucleus is approximately an ellipsoid with major axis length ∼20μm, which is consistent with the expected nucleus size of IMR-90 cells. (**B**) Histogram of the maximum distance between pairs of DNA-MERFISH probes for all 3,668 cells in the DNA-MERFISH data set. The maximum distance for each cell is used to normalize distances between probes in that cell. (**C**) Histogram of the number of cells in which a pair of DNA-MERFISH probes is found. The distance metric we use in the main text, i.e., the average cell-normalized Euclidean distance between two probes, corresponds to normalizing the distance between two probes in a cell by the maximum probe-probe distance in the cell in which the two probes are found, then taking the average over all cells that contain these two probes. The histogram shows that this average is taken on at least 2,800 cells and should thus enjoy a high statistical accuracy. (**D**) Histogram of average cell-normalized Euclidean distances between adhesome DNA-MERFISH probes. Intrachromosomal probe-probe distances are shown in blue, while interchromosomal probe-probe distances are shown in orange. (**E**) Scatter plot of max-min scaled Hi-C contacts and cell-normalized DNA-MERFISH Euclidean proximity (defined as (1-Euclidean distance)) for interchromosomal adhesome gene pairs (blue, *p*-value < 3e-308, see ^3^) and intrachromosomal adhesome gene pairs (orange, *p*-value < 3e-308).

**Supplementary Fig. 18:**
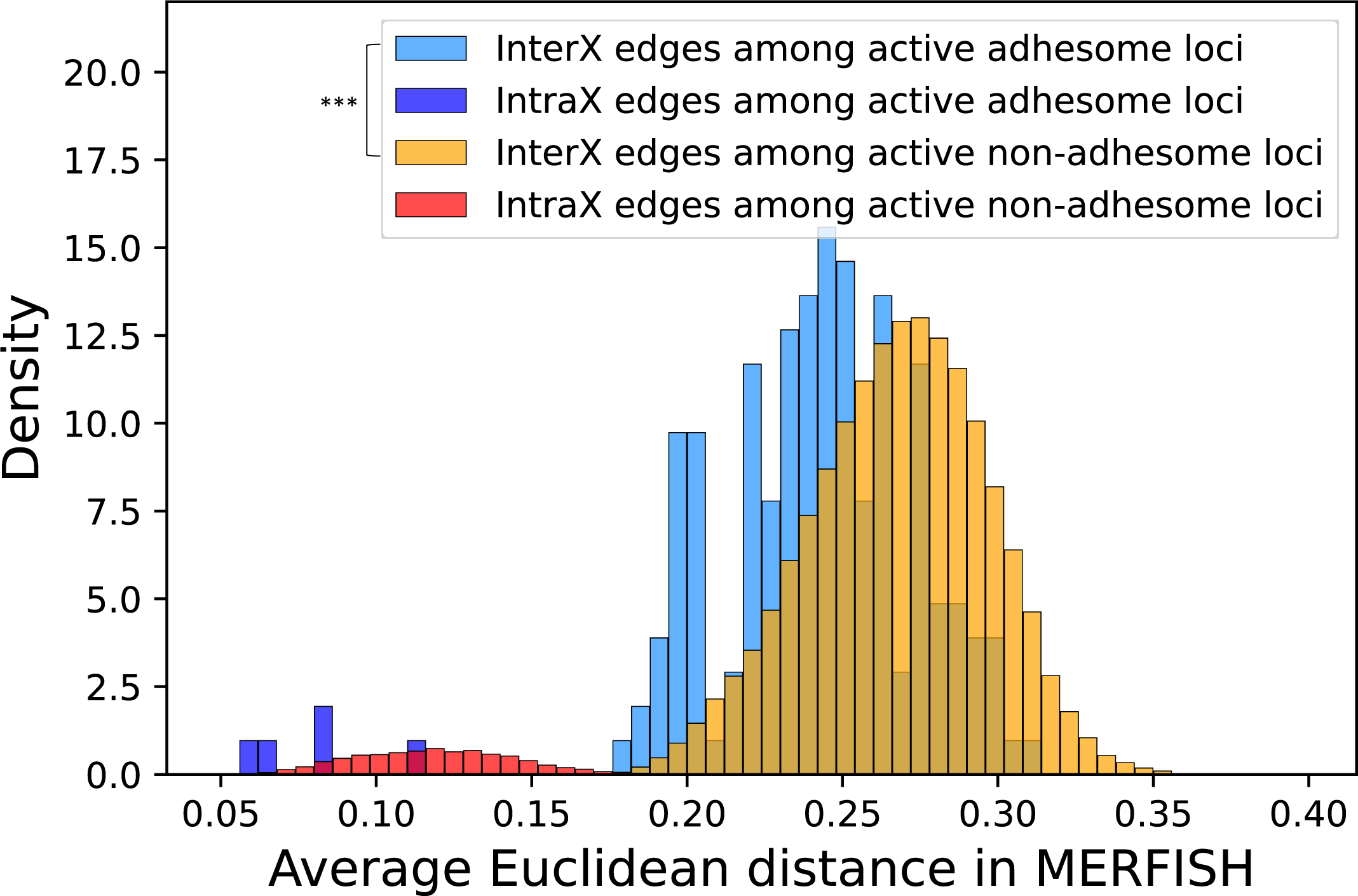
Extended MERFISH analysis. Distribution of average normalized Euclidean distances between active adhesome gene probes and active non-adhesome gene probes (a MERFISH probe is associated with a gene if it overlaps with the gene). The average interchromosomal Euclidean distance between active adhesome probes is smaller than the average interchromosomal Euclidean distance among active non-adhesome probes (Wilcoxon Rank-Sum *p*-value < 1e-18).

**Supplementary Fig. 19:**
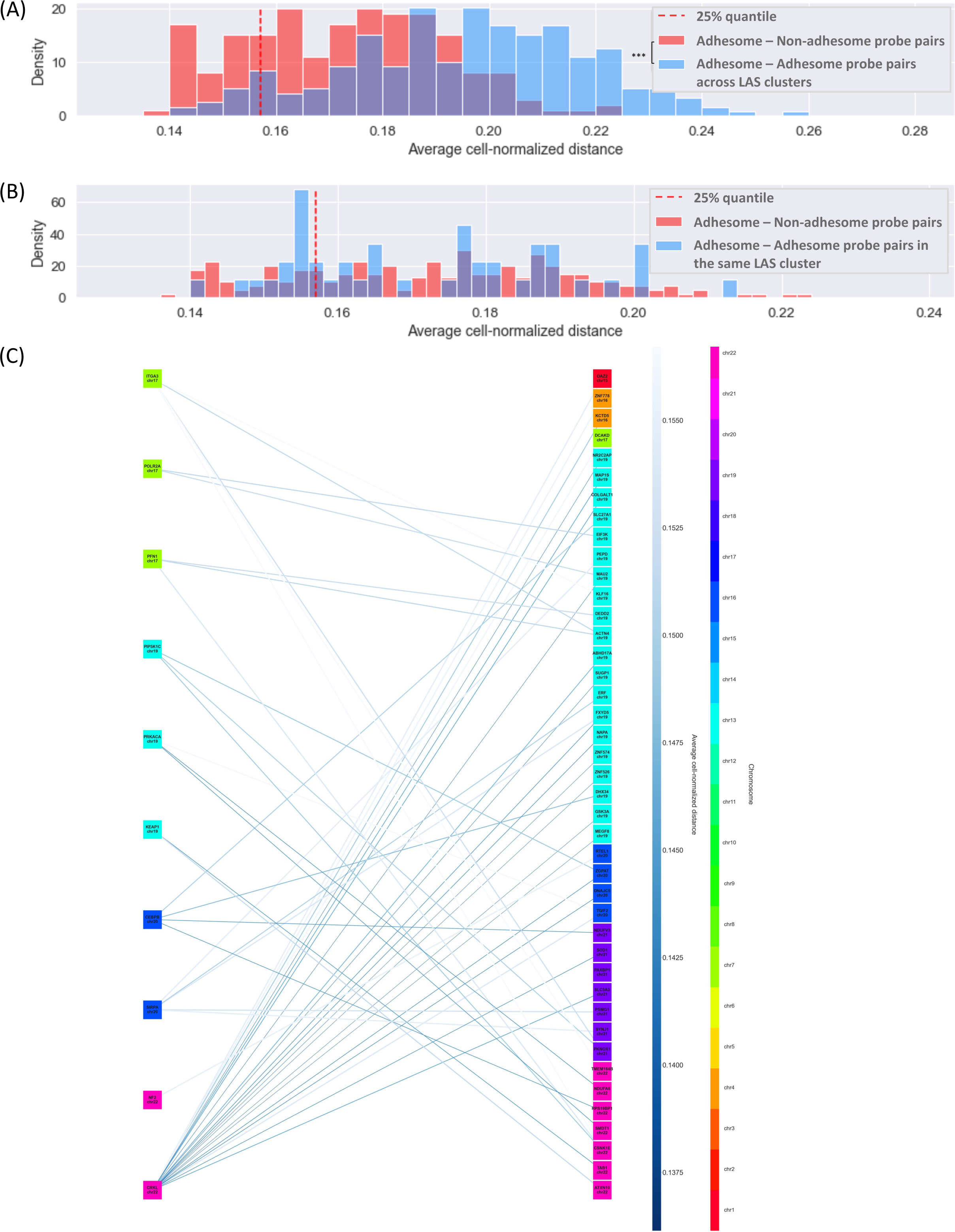
Bipartite network of active non-adhesome genes interacting with active adhesome genes. (**A**) Red histogram of the average cell-normalized Euclidean distance between 135 active non-adhesome genes (61 MERFISH probes) and the 17 active adhesome genes (16 MERFISH probes) they interact with. MERFISH probes are associated with a gene if they are less than 100 kb upstream or downstream of the gene. Only gene pairs that interact in LAS and have a strong regulatory correlation (upper decile in Supplementary Fig. 10A) are considered. The red vertical dashed line corresponds to an upper threshold (first quartile) on the distance used to select edges for (C). Blue histogram of the average cell-normalized Euclidean distance between 23 active adhesome MERFISH probes across LAS regions, used for comparison with the red histogram (Wilcoxon Rank Sums test, *p*-value<2e-29). (**B**) Red histogram as in (A). Blue histogram of the average cell-normalized Euclidean distance between 13 active adhesome MERFISH probes (only LAS interactions are reported) (Wilcoxon Rank Sums test, *p*-value=0.75). (**C**) Bipartite network of active adhesome genes (10 genes, left) and active non-adhesome genes they interact with (42 genes, right). Only genes with an associated MERFISH probe (less than 100 kb upstream or downstream) are considered; more details on how this network is constructed can be found in the main text. Gene colors indicate the corresponding chromosomes. Edge colors and thicknesses reflect the physical distance between genes. Edges with an average cell-normalized distance above the first quartile threshold of (A) are not drawn.

**Supplementary Fig. 20:**
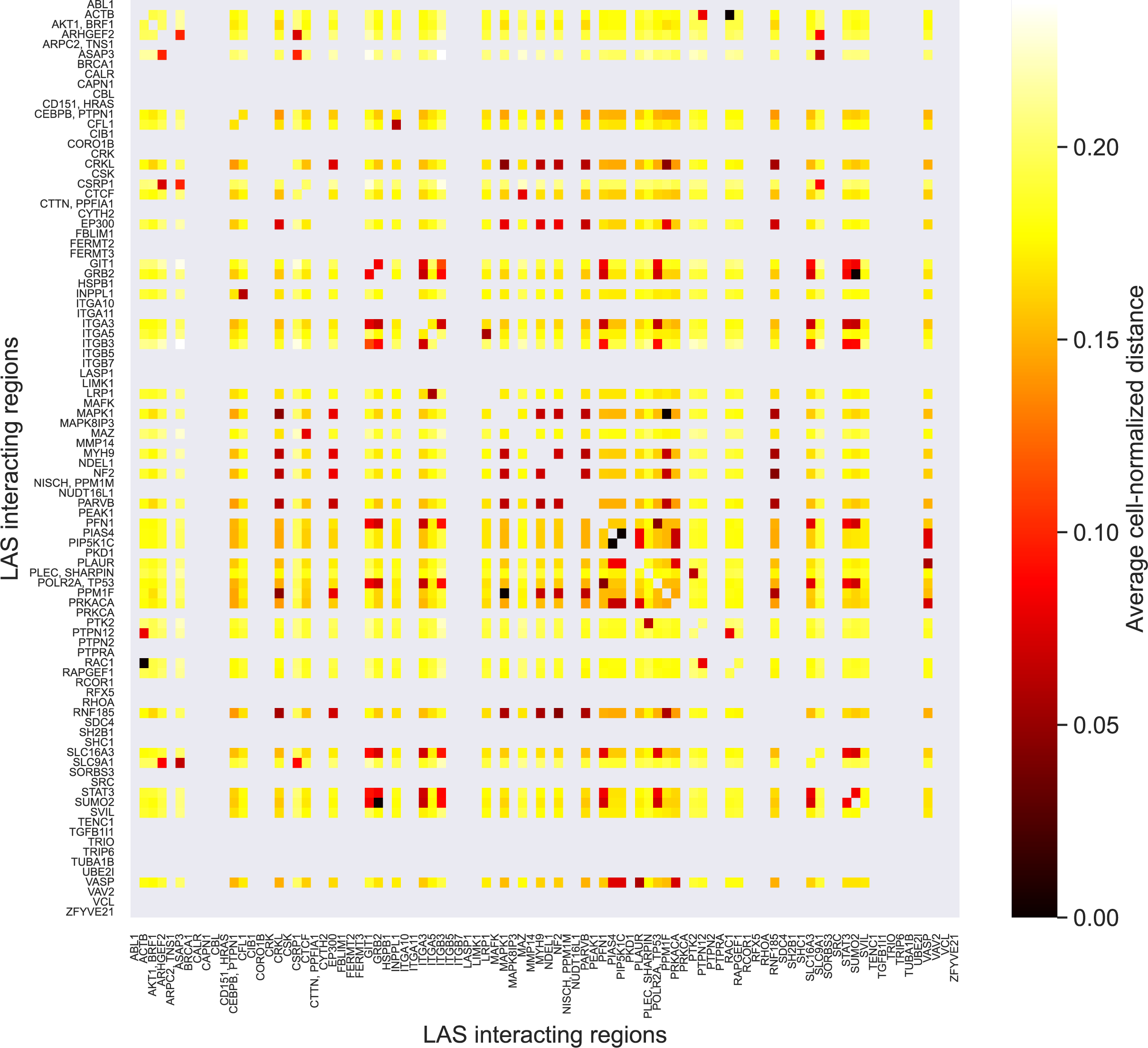
Relationship between Hi-C contact frequency and DNA-MERFISH Euclidean distance between adhesome loci. Heatmap of average cell-normalized DNA-MERFISH Euclidean distance between adhesome LAS nodes. The distance between two nodes is defined by averaging the distances between all DNA-MERFISH probes associated to the first node and all DNA-MERFISH probes associated to the second node. A DNA-MERFISH probe is associated to a given adhesome LAS node if it is located at most 500kb upstream or downstream of an adhesome gene within that node. Adhesome LAS nodes that do not have an associated DNA-MERFISH probe are represented as grey rows/columns in the heatmap (missing values). Self-distances are also shown in grey. Dark entries (very low distance) typically correspond to pairs of adhesome LAS nodes on the same chromosome.

**Supplementary Fig. 21:**
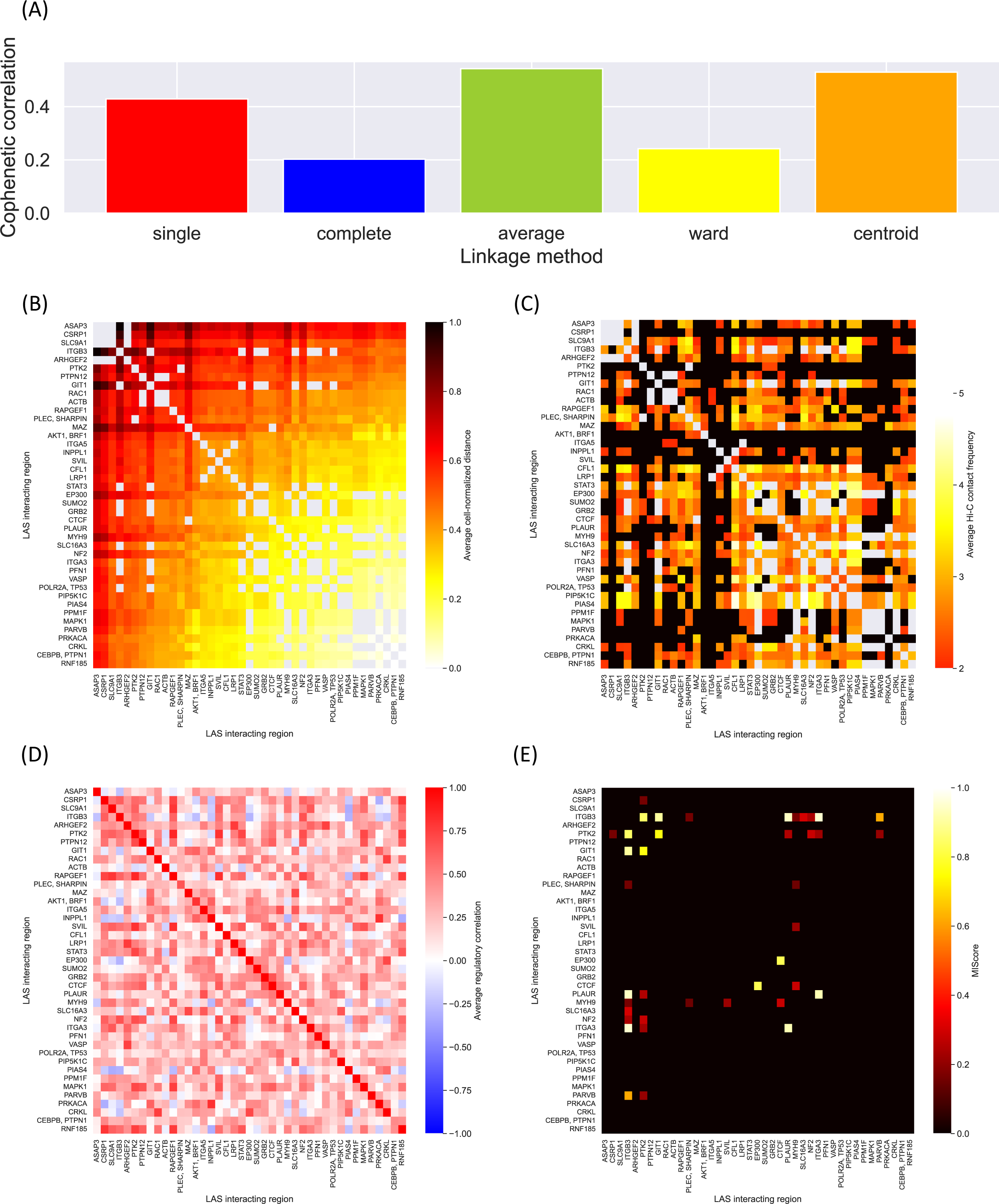
Joint analysis of DNA-MERFISH data, Hi-C data, regulatory data, and protein-protein interaction data identifies a cluster of highly co-localized and co-regulated adhesome genes. (**A**) Bar plot of cophenetic correlation for different linkage functions applied to the average cell-normalized DNA-MERFISH Euclidean distance matrix between adhesome LAS nodes having at least one associated DNA-MERFISH probe. (**B**) Heatmap of average cell-normalized DNA-MERFISH Euclidean distance between adhesome LAS nodes having at least one associated DNA-MERFISH probe. Rows and columns are ordered using hierarchical clustering with single linkage. Single linkage produces a dendrogram that is reasonably faithful to the original Euclidean distances (cophenetic correlation ∼0.4, see (A)) and yields an interpretable ordering of the heatmap. Entries of the heatmap that correspond to self-interactions and intrachromosomal interactions are shown in grey. (**C**) Heatmap of average transformed Hi-C contact frequency between adhesome LAS nodes having at least one associated DNA-MERFISH probe. Rows and columns are ordered as in (B). Entries of the heatmap that correspond to self-interactions and intrachromosomal interactions are shown in grey. Entries corresponding to interactions that were not identified by the LAS algorithm are shown in black. (**D**) Heatmap of average Spearman regulatory correlation between adhesome LAS nodes having at least one associated DNA-MERFISH probe. Rows and columns are ordered as in (B). (**E**) Heatmap of protein-protein physical interaction confidence score between genes in adhesome LAS nodes having at least one associated DNA-MERFISH probe. Rows and columns are ordered as in (B). Protein-protein physical interactions are retrieved from the physical STRING network. The confidence score for the interaction of two proteins is based on the number and type of sources reporting this interaction. A score of 0 indicates that two proteins are not known to physically interact.

**Supplementary Fig. 22:**
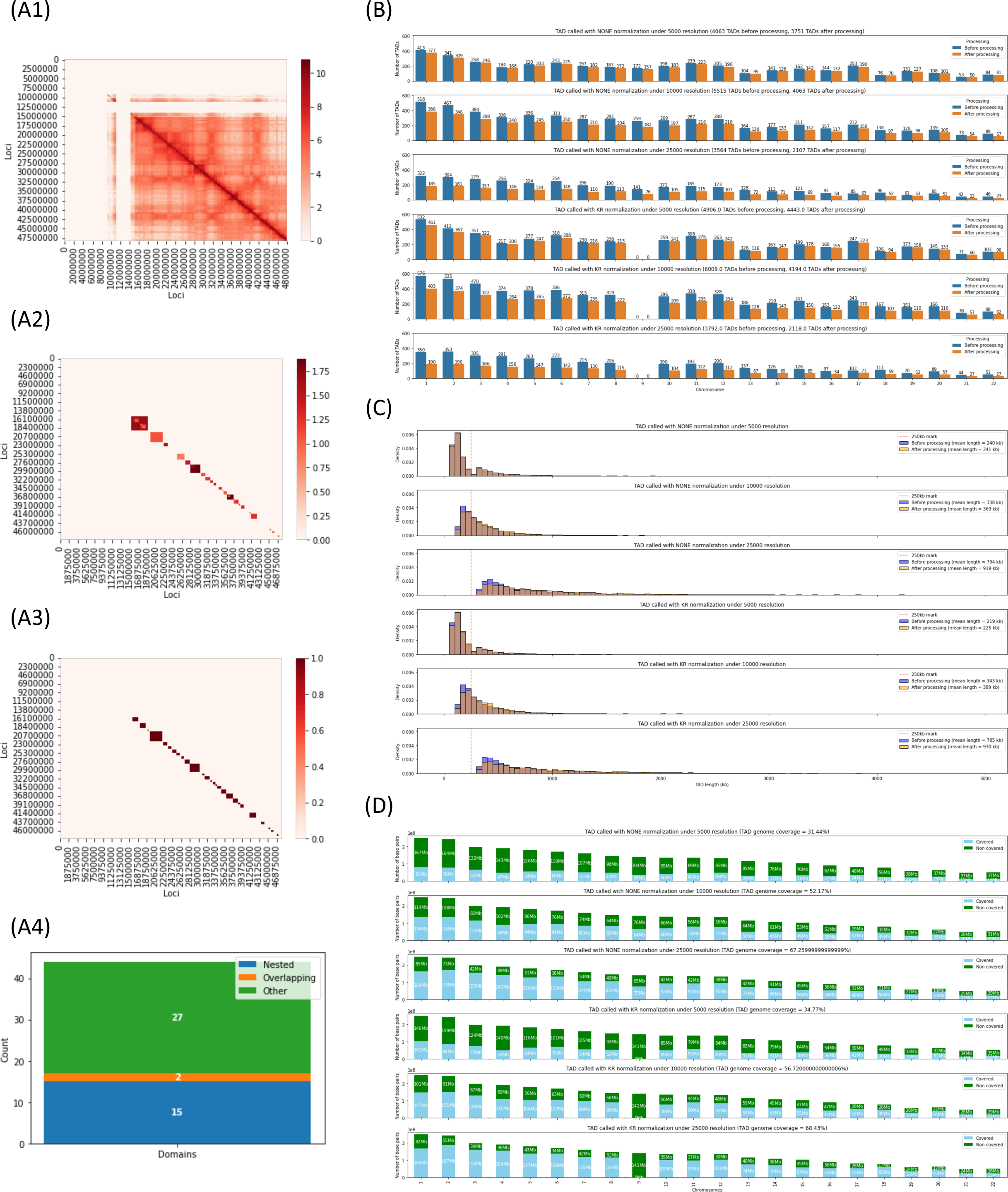
TAD calling. (**A**) TAD calling pipeline shown for chromosome 21 at resolution 25 kb with Knight-Ruiz normalization. The Arrowhead software (see Methods section) is applied to the normalized intrachromosomal Hi-C contact matrix of chromosome 21 at a given resolution (A1), resulting in a list of potentially overlapping and nested TADs (A2). This TAD list is then processed to remove nested TADs and merge overlapping TADs (A3). Most TADs are disjoint, yet a minority of TADs are nested and only very few TADs are overlapping (A4). (**B**) Number of TADs called per chromosome before and after post-processing for TAD lists called by Arrowhead at different resolutions (5 kb, 10 kb, 25 kb), with and without Knight-Ruiz normalization. (**C**) Distribution of TAD length before and after post-processing for TAD lists called by Arrowhead at different resolutions (5 kb, 10 kb, 25 kb), with and without Knight-Ruiz normalization. (**D**) Number of base pairs covered per chromosome by processed TADs for TAD lists called by Arrowhead at different resolutions (5 kb, 10 kb, 25 kb), with and without Knight-Ruiz normalization.

**Supplementary Fig. 23:**
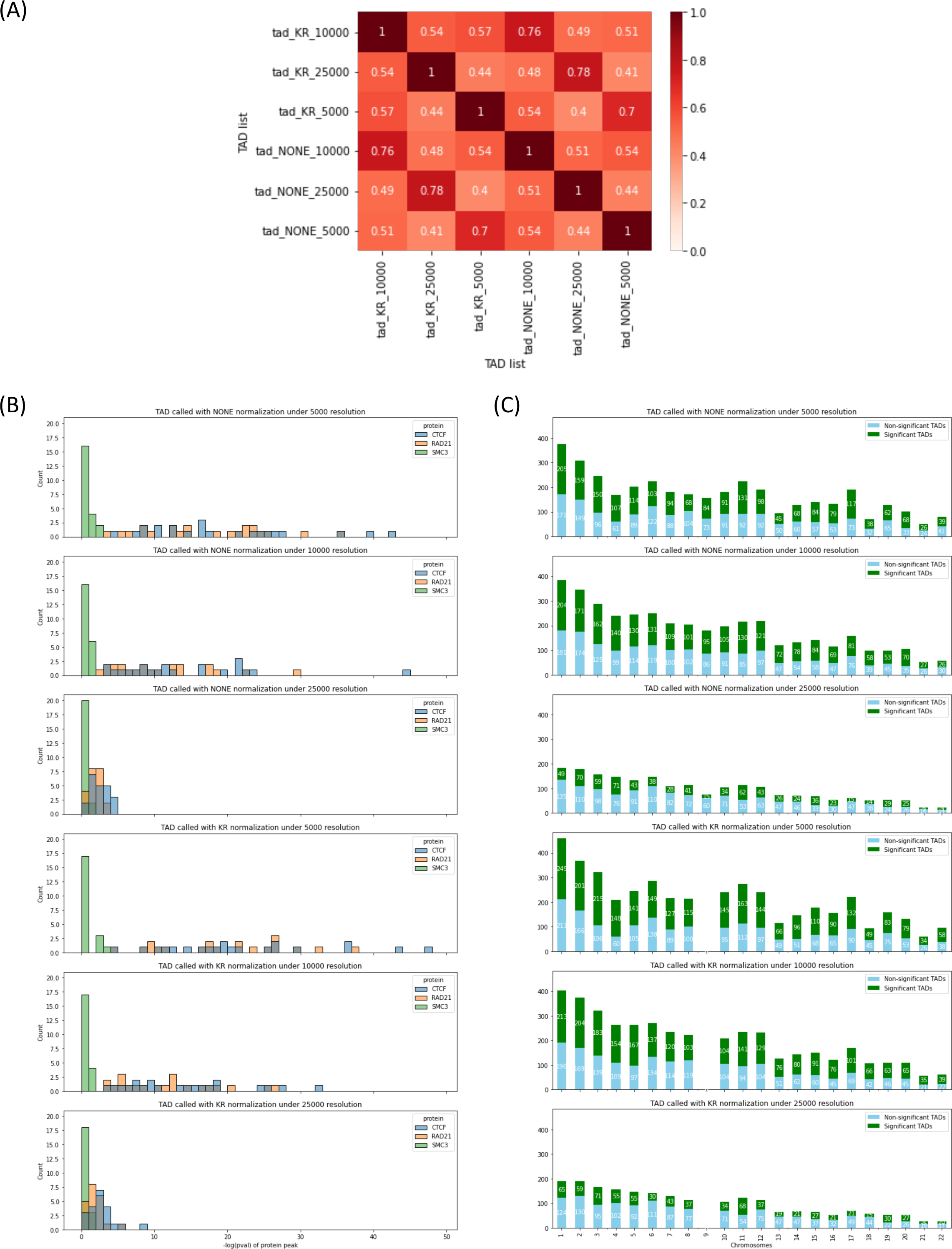
TAD quality. (**A**) Measure of Concordance (MoC) heatmap for TAD lists called by Arrowhead at different resolutions (5 kb, 10 kb, 25 kb), with and without Knight-Ruiz normalization. An MoC of 1 indicates that 2 TAD lists are identical, while an MoC of 0 indicates that 2 TAD lists are non-overlapping (see Methods). (**B**) Negative log-*p*-value of the structural protein profiles (see Methods) of CTCF, RAD21, and SMC3 for TAD lists called by Arrowhead at different resolutions (5 kb, 10 kb, 25 kb), with and without Knight-Ruiz normalization. (**C**) Number of TADs per chromosome significantly enriched for either transcriptional activating (H3K36me3) or repressing (H3K27me3) histone marks for TAD lists called by Arrowhead at different resolutions (5 kb, 10 kb, 25 kb), with and without Knight-Ruiz normalization. Enrichment is estimated as the log10-ratio between the ChIP-seq signals of H3K36me3 and H3K27me3 and compared with a null distribution derived from random permutations (see Methods).

**Supplementary Fig. 24:**
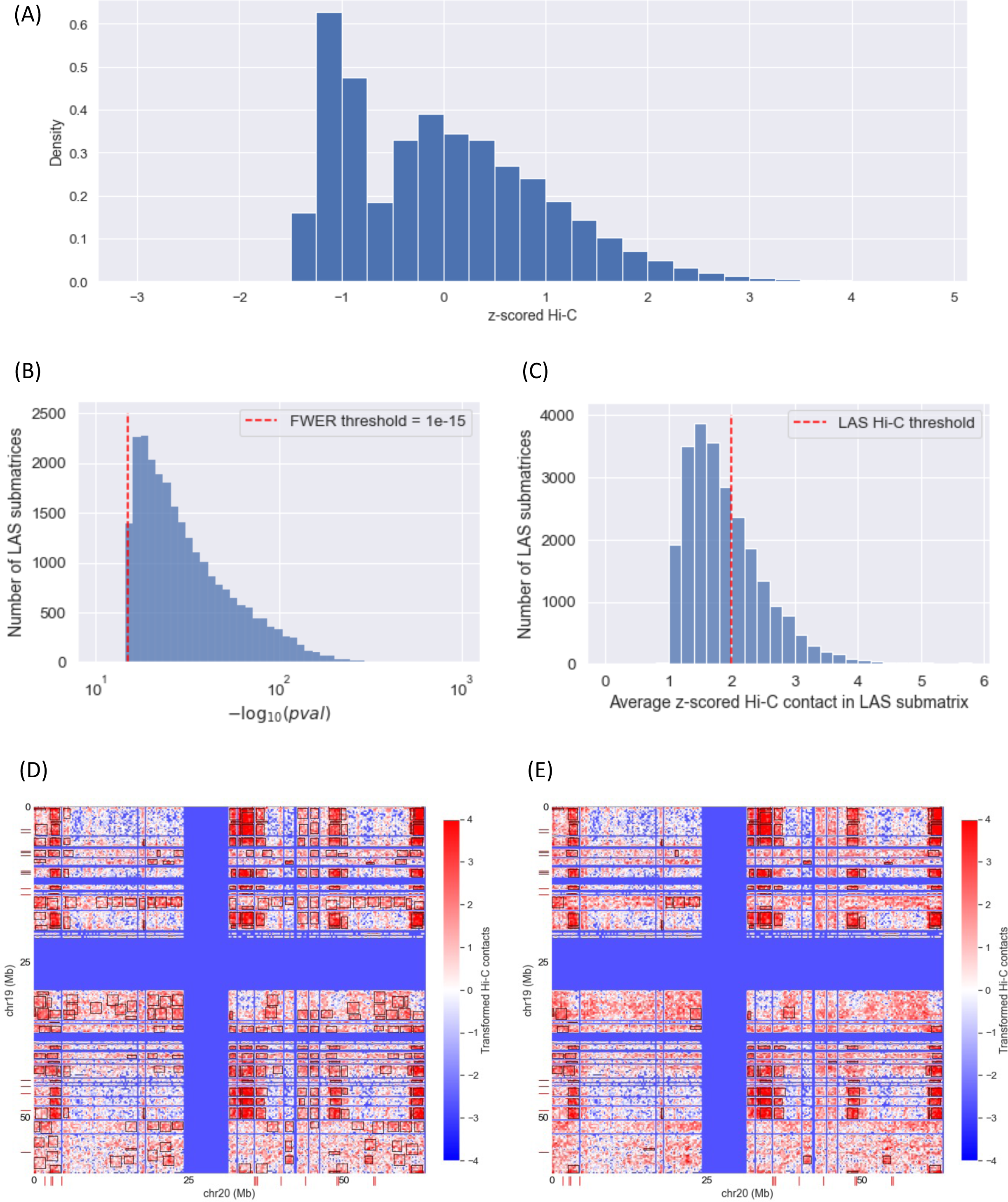
Clustering of transformed Hi-C contact frequencies using LAS algorithm. (**A**) Histogram of non-zero log-transformed and centerdized Hi-C contact values in the whole genome. Centerdization was performed using the mean and standard deviation of the non-zero contact values across the whole genome. (**B**) Histogram of negative log10-transformed *p*- values of submatrices identified by the LAS algorithm ^3^ in all interchromosomal Hi-C contact matrices. The *p*-value threshold of 1e-15 used for signal detection is represented as a red dashed vertical line. This *p*-value threshold controls the FWER at <0.0001. (**C**) Histogram of average transformed Hi-C contact values in all LAS submatrices. The Hi-C threshold of 2 used to discard low-average submatrices is represented as a red dashed vertical line. (**D**) Heatmap of transformed Hi-C contact frequencies between genomic loci on chromosome 19 and chromosome 20. Adhesome loci are indicated by red ticks on the x-axis (for chromosome 20) and the y-axis (for chromosome 19). Significant submatrices identified by the LAS algorithm are shown in black boxes. (**E**) Same as (D), except that only significant submatrices identified by the LAS algorithm with an average Hi-C contact value higher than 2 are shown in black boxes.

## Notes

### Competing Interest Statement

The authors have declared no competing interest.

