## Supplementary material for "Adhesome Receptor Clustering is Accompanied by the Co- localization of the Associated Genes in the Cell Nucleus": SI Appendix

**This PDF file includes:**

SI Figures S1 to S24

SI References

**SI Figures**

| Gene | Functional Category | FA | activity |
|---|---|---|---|
| ACTN1 | Actin regulation | Intrinsic Proteins | active |
| CFL1 | Actin regulation | Intrinsic Proteins | active |
| CORO1B | Actin regulation | Intrinsic Proteins | active |
| CTTN | Actin regulation | Intrinsic Proteins | active |
| FLNA | Actin regulation | Intrinsic Proteins | NA |
| KEAP1 | Actin regulation | Intrinsic Proteins | active |
| LASP1 | Actin regulation | Intrinsic Proteins | active |
| ENAH | Actin regulation | Intrinsic Proteins | active |
| NEXN | Actin regulation | Intrinsic Proteins | active |
| SVIL | Actin regulation | Intrinsic Proteins | active |
| VASP | Actin regulation | Intrinsic Proteins | active |
| CORO2A | Actin regulation | Intrinsic Proteins | active |
| ACTB | Adaptor | Intrinsic Proteins | active |
| JUB | Adaptor | Intrinsic Proteins | NA |
| SORBS2 | Adaptor | Intrinsic Proteins | active |
| BCAR1 | Adaptor | Intrinsic Proteins | active |
| CAV1 | Adaptor | Intrinsic Proteins | active |
| SMPX | Adaptor | Intrinsic Proteins | NA |
| SH3KBP1 | Adaptor | Intrinsic Proteins | NA |
| CRK | Adaptor | Intrinsic Proteins | active |
| CRKL | Adaptor | Intrinsic Proteins | active |
| EZR | Adaptor | Intrinsic Proteins | active |
| FHL2 | Adaptor | Intrinsic Proteins | active |
| GAB1 | Adaptor | Intrinsic Proteins | active |
| GRB2 | Adaptor | Intrinsic Proteins | active |
| GRB7 | Adaptor | Intrinsic Proteins | active |
| HAX1 | Adaptor | Intrinsic Proteins | active |
| NEDD9 | Adaptor | Intrinsic Proteins | active |
| CASS4 | Adaptor | Intrinsic Proteins | active |
| TGFB1I1 | Adaptor | Intrinsic Proteins | active |
| ITGB1BP1 | Adaptor | Intrinsic Proteins | active |
| FERMT1 | Adaptor | Intrinsic Proteins | inactive |
| FERMT2 | Adaptor | Intrinsic Proteins | active |
| FERMT3 | Adaptor | Intrinsic Proteins | active |
| LPXN | Adaptor | Intrinsic Proteins | active |
| PPFIA1 | Adaptor | Intrinsic Proteins | active |
| LPP | Adaptor | Intrinsic Proteins | active |
| NF2 | Adaptor | Intrinsic Proteins | active |
| FBLIM1 | Adaptor | Intrinsic Proteins | active |
| MSN | Adaptor | Intrinsic Proteins | NA |
| NCK2 | Adaptor | Intrinsic Proteins | active |
| PALLD | Adaptor | Intrinsic Proteins | active |
| PARVA | Adaptor | Intrinsic Proteins | active |
| PARVB | Adaptor | Intrinsic Proteins | active |
| PXN | Adaptor | Intrinsic Proteins | inactive |
| LIMS1 | Adaptor | Intrinsic Proteins | active |
| LIMS2 | Adaptor | Intrinsic Proteins | active |
| GNB2L1 | Adaptor | Intrinsic Proteins | active |
| RDX | Adaptor | Intrinsic Proteins | active |
| OSTF1 | Adaptor | Intrinsic Proteins | active |
| NUDT16L1 | Adaptor | Intrinsic Proteins | active |
| SYNM | Adaptor | Intrinsic Proteins | active |
| SDCBP | Adaptor | Intrinsic Proteins | active |
| TLN1 | Adaptor | Intrinsic Proteins | active |
| TNS1 | Adaptor | Intrinsic Proteins | active |
| TES | Adaptor | Intrinsic Proteins | active |
| TRIP6 | Adaptor | Intrinsic Proteins | active |
| VCL | Adaptor | Intrinsic Proteins | active |
| SORBS3 | Adaptor | Intrinsic Proteins | active |
| LDB3 | Adaptor | Intrinsic Proteins | active |
| ZYX | Adaptor | Intrinsic Proteins | active |
| NDEL1 | Adaptor | Intrinsic Proteins | active |
| SH2B1 | Adaptor | Intrinsic Proteins | active |
| TENC1 | Adaptor | Intrinsic Proteins | active |
| ZFYVE21 | Adaptor | Intrinsic Proteins | active |
| SLC3A2 | Adhesion receptor | Intrinsic Proteins | active |
| KTN1 | Adhesion receptor | Intrinsic Proteins | active |
| LRP1 | Adhesion receptor | Intrinsic Proteins | active |
| PVR | Adhesion receptor | Intrinsic Proteins | active |
| SDC4 | Adhesion receptor | Intrinsic Proteins | active |
| ITGA1 | Adhesion receptor | Intrinsic Proteins | active |
| ITGA2 | Adhesion receptor | Intrinsic Proteins | active |
| ITGA3 | Adhesion receptor | Intrinsic Proteins | active |
| ITGA4 | Adhesion receptor | Intrinsic Proteins | active |
| ITGA5 | Adhesion receptor | Intrinsic Proteins | active |
| ITGA6 | Adhesion receptor | Intrinsic Proteins | active |
| ITGA7 | Adhesion receptor | Intrinsic Proteins | inactive |
| ITGA8 | Adhesion receptor | Intrinsic Proteins | active |
| ITGA9 | Adhesion receptor | Intrinsic Proteins | active |
| ITGA10 | Adhesion receptor | Intrinsic Proteins | active |
| ITGA11 | Adhesion receptor | Intrinsic Proteins | active |
| ITGAD | Adhesion receptor | Intrinsic Proteins | inactive |
| ITGAE | Adhesion receptor | Intrinsic Proteins | active |
| ITGAL | Adhesion receptor | Intrinsic Proteins | inactive |
| ITGAM | Adhesion receptor | Intrinsic Proteins | inactive |
| ITGAV | Adhesion receptor | Intrinsic Proteins | active |
| ITGAW | Adhesion receptor | Intrinsic Proteins | NA |
| ITGAX | Adhesion receptor | Intrinsic Proteins | inactive |
| ITGB1 | Adhesion receptor | Intrinsic Proteins | active |
| ITGB2 | Adhesion receptor | Intrinsic Proteins | active |
| ITGB3 | Adhesion receptor | Intrinsic Proteins | active |
| ITGB4 | Adhesion receptor | Intrinsic Proteins | inactive |
| ITGB5 | Adhesion receptor | Intrinsic Proteins | active |
| ITGB6 | Adhesion receptor | Intrinsic Proteins | active |
| ITGB7 | Adhesion receptor | Intrinsic Proteins | active |
| ITGB8 | Adhesion receptor | Intrinsic Proteins | active |
| NRP1 | Adhesion receptor | Intrinsic Proteins | active |
| NRP2 | Adhesion receptor | Intrinsic Proteins | active |
| CD151 | Adhesion receptor | Intrinsic Proteins | active |
| PDE4D | cAMP phosphodiesterase | Intrinsic Proteins | active |
| PKD1 | Channel | Intrinsic Proteins | active |
| TRPM7 | Channel | Intrinsic Proteins | active |
| CALR | Chaperone | Intrinsic Proteins | active |
| DDEF1 | GAP | Intrinsic Proteins | NA |
| ASAP3 | GAP | Intrinsic Proteins | active |
| GIT1 | GAP | Intrinsic Proteins | active |
| GIT2 | GAP | Intrinsic Proteins | active |
| ARHGAP26 | GAP | Intrinsic Proteins | active |
| GRLF1 | GAP | Intrinsic Proteins | NA |
| ASAP2 | GAP | Intrinsic Proteins | active |
| ARHGAP24 | GAP | Intrinsic Proteins | active |
| DLC1 | GAP | Intrinsic Proteins | active |
| AGAP2 | GAP | Intrinsic Proteins | active |
| STARD13 | GAP | Intrinsic Proteins | active |
| RASA1 | GAP | Intrinsic Proteins | active |
| DEF6 | GEF | Intrinsic Proteins | active |
| DOCK1 | GEF | Intrinsic Proteins | active |
| ELMO1 | GEF | Intrinsic Proteins | active |
| ARHGEF6 | GEF | Intrinsic Proteins | NA |
| ARHGEF7 | GEF | Intrinsic Proteins | active |
| DNM2 | GTPase | Intrinsic Proteins | active |
| RHOU | GTPase | Intrinsic Proteins | active |
| PLCG1 | Phospholipase | Intrinsic Proteins | active |
| INPP5D | PtdIns phosphatase | Intrinsic Proteins | inactive |
| INPPL1 | PtdIns phosphatase | Intrinsic Proteins | active |
| ITGB3BP | RNA or DNA regulation | Intrinsic Proteins | active |
| RAVER1 | RNA or DNA regulation | Intrinsic Proteins | active |
| STAT3 | RNA or DNA regulation | Intrinsic Proteins | active |
| SPTLC1 | Serine palmitoyltransferase | Intrinsic Proteins | active |
| ILK | Serine/threonine kinase | Intrinsic Proteins | active |
| PAK1 | Serine/threonine kinase | Intrinsic Proteins | active |
| PDPK1 | Serine/threonine kinase | Intrinsic Proteins | inactive |
| PRKCA | Serine/threonine kinase | Intrinsic Proteins | active |
| PPM1M | Serine/threonine phosphatase | Intrinsic Proteins | active |
| PPM1F | Serine/threonine phosphatase | Intrinsic Proteins | active |
| PPP2CA | Serine/threonine phosphatase | Intrinsic Proteins | NA |
| ABL1 | Tyrosine Kinase | Intrinsic Proteins | active |
| CSK | Tyrosine Kinase | Intrinsic Proteins | active |
| PTK2 | Tyrosine Kinase | Intrinsic Proteins | active |
| PTK2B | Tyrosine Kinase | Intrinsic Proteins | active |
| SRC | Tyrosine Kinase | Intrinsic Proteins | active |
| PEAK1 | Tyrosine kinase | Intrinsic Proteins | active |
| PTPRF | Tyrosine phosphatase | Intrinsic Proteins | active |
| PTPN12 | Tyrosine phosphatase | Intrinsic Proteins | active |
| PTPRA | Tyrosine phosphatase | Intrinsic Proteins | active |
| PTPN6 | Tyrosine phosphatase | Intrinsic Proteins | active |
| PTPN11 | Tyrosine phosphatase | Intrinsic Proteins | active |
| PRNP | Unknown | Intrinsic Proteins | active |
| MYH9 | Actin regulation | Intrinsic Proteins | active |
| MACF1 | Actin regulation | Intrinsic Proteins | active |
| ARPC2 | Actin regulation | Associated Proteins | active |
| MARCKS | Actin regulation | Associated Proteins | active |
| PFN1 | Actin regulation | Associated Proteins | active |
| ABI1 | Adaptor | Associated Proteins | active |
| ABI2 | Adaptor | Associated Proteins | active |
| ABI3 | Adaptor | Associated Proteins | inactive |
| ANKRD28 | Adaptor | Associated Proteins | active |
| CSRP1 | Adaptor | Associated Proteins | active |
| IRS1 | Adaptor | Associated Proteins | active |
| MAPK8IP3 | Adaptor | Associated Proteins | active |
| PLEC | Adaptor | Associated Proteins | active |
| SORBS1 | Adaptor | Associated Proteins | active |
| SHC1 | Adaptor | Associated Proteins | active |
| MYOM1 | Adaptor | Associated Proteins | inactive |
| TSPAN1 | Adaptor | Associated Proteins | active |
| TUBA1B | Adaptor | Associated Proteins | active |
| VIM | Adaptor | Associated Proteins | active |
| SHARPIN | Adaptor | Associated Proteins | active |
| FABP3 | adaptor | Associated Proteins | active |
| NISCH | adaptor | Associated Proteins | active |
| CIB1 | adaptor | Associated Proteins | active |
| CIB2 | adaptor | Associated Proteins | inactive |
| SRCIN1 | adaptor | Associated Proteins | inactive |
| ADAM12 | Adhesion receptor | Associated Proteins | active |
| CEACAM1 | Adhesion receptor | Associated Proteins | inactive |
| ENG | Adhesion receptor | Associated Proteins | inactive |
| CD47 | Adhesion receptor | Associated Proteins | active |
| LAYN | Adhesion receptor | Associated Proteins | active |
| SIRPA | Adhesion receptor | Associated Proteins | active |
| THY1 | Adhesion receptor | Associated Proteins | active |
| PLAUR | Adhesion receptor | Associated Proteins | active |
| KCNH2 | Channel | Associated Proteins | inactive |
| SLC16A3 | Channel | Associated Proteins | active |
| SLC9A1 | Channel | Associated Proteins | active |
| HSPB1 | Chaperone | Associated Proteins | active |
| HSPA2 | Chaperone | Associated Proteins | active |
| CBL | E3-ligase | Associated Proteins | active |
| RNF5 | E3-ligase | Associated Proteins | active |
| RNF185 | E3-ligase | Associated Proteins | active |
| ARHGAP5 | GAP | Associated Proteins | active |
| ARHGAP32 | GAP | Associated Proteins | active |
| BCAR3 | GEF | Associated Proteins | active |
| RAPGEF1 | GEF | Associated Proteins | active |
| SOS1 | GEF | Associated Proteins | active |
| TIAM1 | GEF | Associated Proteins | active |
| TRIO | GEF | Associated Proteins | active |
| VAV1 | GEF | Associated Proteins | inactive |
| VAV2 | GEF | Associated Proteins | active |
| VAV3 | GEF | Associated Proteins | inactive |
| ARHGEF12 | GEF | Associated Proteins | active |
| ARHGEF2 | GEF | Associated Proteins | active |
| CYTH2 | GEF | Associated Proteins | active |
| ARF1 | GTPase | Associated Proteins | active |
| HRAS | GTPase | Associated Proteins | active |
| RAC1 | GTPase | Associated Proteins | active |
| RHOA | GTPase | Associated Proteins | active |
| PLD1 | Phospholipase | Associated Proteins | active |
| CAPN1 | Protease | Associated Proteins | active |
| CAPN2 | Protease | Associated Proteins | active |
| CASP8 | Protease | Associated Proteins | active |
| MMP14 | Protease | Associated Proteins | active |
| PIK3CA | PtdIns kinase | Associated Proteins | active |
| PIP5K1C | PtdIns kinase | Associated Proteins | active |
| PTEN | PtdIns phosphatase | Associated Proteins | active |
| PABPC1 | RNA or DNA regulation | Associated Proteins | active |
| SSH1 | Serine phosphatase | Associated Proteins | active |
| AKT1 | Serine/threonine kinase | Associated Proteins | active |
| MAPK1 | Serine/threonine kinase | Associated Proteins | active |
| MAPK8 | Serine/threonine kinase | Associated Proteins | active |
| LIMK1 | Serine/threonine kinase | Associated Proteins | active |
| PRKACA | Serine/threonine kinase | Associated Proteins | active |
| ROCK1 | Serine/threonine kinase | Associated Proteins | active |
| ILKAP | Serine/threonine phosphatase | Associated Proteins | active |
| INSR | Tyrosine Kinase | Associated Proteins | active |
| FYN | Tyrosine Kinase | Associated Proteins | active |
| LYN | Tyrosine Kinase | Associated Proteins | active |
| SYK | Tyrosine Kinase | Associated Proteins | inactive |
| TESK1 | Tyrosine Kinase | Associated Proteins | active |
| PTPN1 | Tyrosine phosphatase | Associated Proteins | active |
| PTPRO | Tyrosine phosphatase | Associated Proteins | active |
| PTPRH | Tyrosine phosphatase | Associated Proteins | active |
| PTPN2 | Tyrosine phosphatase | Associated Proteins | active |

Active gene

Inactive gene

Missing gene

### Supplementary Figure 1

**Supplementary Fig. 1: List of adhesome genes and their activity.** Intrinsic and associated adhesome genes from [1] colored by activity level.

(A)

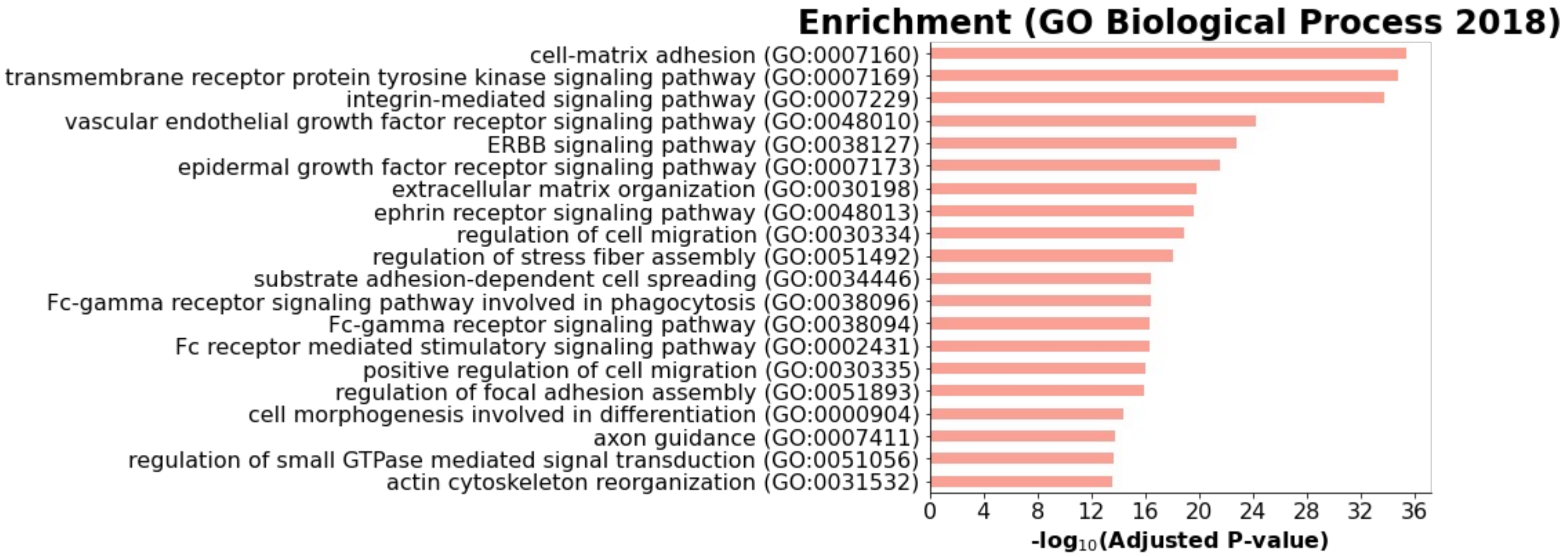


(B)

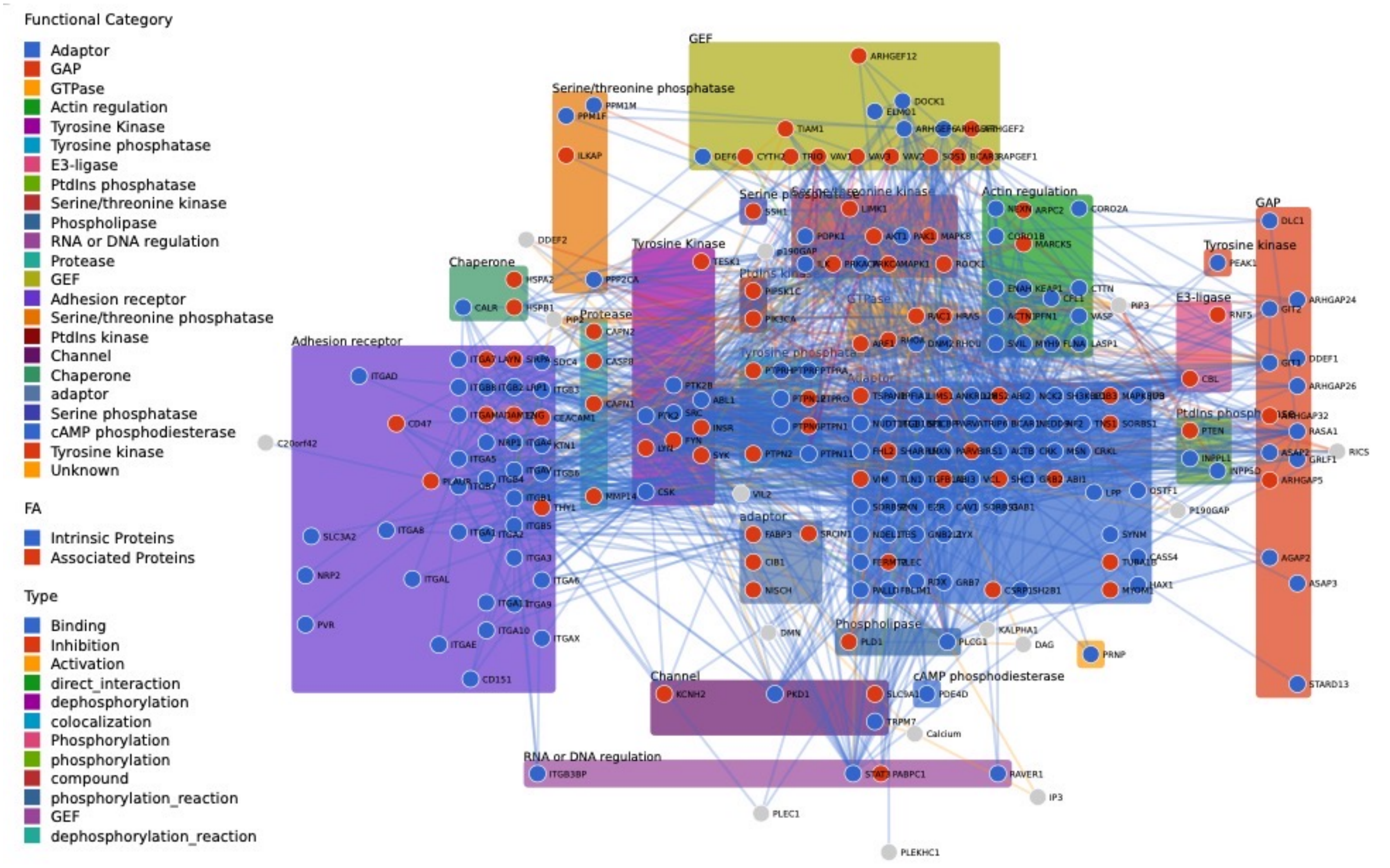


**Supplementary Figure 2**

**Supplementary Fig. 2: Function and structure of the adhesome.** (**A**) Gene Set Enrichment Analysis of intrinsic and associated adhesome components using the 2018 Gene Ontology database for biological processes. (**B**) Network of interactions among adhesome proteins. Nodes are colored in blue if they are intrinsic adhesome proteins, and in red if they are associated adhesome proteins. Nodes are grouped by functional categories. Links between nodes are colored based on the type of interaction.

(A)

| Regulatory mark | Retrieved | Original source | Accession | Category |
|---|---|---|---|---|
| RNA-seq | ENCODE | ENCODE | ENCSR424FAZ | active |
| CEBPB | TargetFinder | ENCODE | GSE31477 | active |
| CHD1 | TargetFinder | ENCODE | GSE31477 | active |
| CTCF | TargetFinder | ENCODE | GSE31477 | active |
| DNase-seq | TargetFinder | Roadmap Epigenomics | GSE18927 | active |
| EP300 | TargetFinder | Jin et al. | GSE43070 | active |
| H2AK5ac | TargetFinder | Roadmap Epigenomics | GSE16256 | active |
| H2AK9ac | TargetFinder | Roadmap Epigenomics | GSE16256 | unknown |
| H2AY | TargetFinder | Chen et al. | GSE54847 | unknown |
| H2AZ | TargetFinder | Roadmap Epigenomics | GSE16256 | unknown |
| H2BK120ac | TargetFinder | Roadmap Epigenomics | GSE16256 | unknown |
| H2BK12ac | TargetFinder | Roadmap Epigenomics | GSE16256 | unknown |
| H2BK15ac | TargetFinder | Roadmap Epigenomics | GSE16256 | unknown |
| H2BK20ac | TargetFinder | Roadmap Epigenomics | GSE16256 | unkown |
| H2BK5ac | TargetFinder | Roadmap Epigenomics | GSE16256 | active |
| H3K14ac | TargetFinder | Roadmap Epigenomics | GSE16256 | active |
| H3K18ac | TargetFinder | Roadmap Epigenomics | GSE16256 | active |
| H3K23ac | TargetFinder | Roadmap Epigenomics | GSE16256 | active |
| H3K27ac | TargetFinder | Roadmap Epigenomics | GSE16256 | unkown |
| H3K27me3 | TargetFinder | Roadmap Epigenomics | GSE16256 | repressive |
| H3K36me3 | TargetFinder | Roadmap Epigenomics | GSE16256 | active |
| H3K4ac | TargetFinder | Roadmap Epigenomics | GSE16256 | unknown |
| H3K4me1 | TargetFinder | Roadmap Epigenomics | GSE16256 | active |
| H3K4me2 | TargetFinder | Roadmap Epigenomics | GSE16256 | active |
| H3K4me3 | TargetFinder | Roadmap Epigenomics | GSE16256 | active |
| H3K56ac | TargetFinder | Roadmap Epigenomics | GSE16256 | unknown |
| H3K79me1 | TargetFinder | Roadmap Epigenomics | GSE16256 | unknown |
| H3K79me2 | TargetFinder | Roadmap Epigenomics | GSE16256 | unknown |
| H3K9ac | TargetFinder | Roadmap Epigenomics | GSE16256 | active |
| H3k9me1 | TargetFinder | Roadmap Epigenomics | GSE16256 | repressive |
| H3K9me3 | TargetFinder | Roadmap Epigenomics | GSE16256 | repressive |
| H4K20me1 | TargetFinder | Roadmap Epigenomics | GSE16256 | repressive |
| H4K5ac | TargetFinder | Roadmap Epigenomics | GSE16256 | unknown |
| H4K8ac | TargetFinder | Roadmap Epigenomics | GSE16256 | active |
| H4K91ac | TargetFinder | Roadmap Epigenomics | GSE16256 | unknown |
| LMNB1 | TargetFinder | Otte et al. | GSE53332 | other |
| MAFK | TargetFinder | ENCODE | GSE31477 | active |
| MAZ | TargetFinder | ENCODE | GSE31477 | active |
| MECP2 | TargetFinder | Maunakea et al. | GSE47678 | other |
| MXI1 | TargetFinder | ENCODE | GSE31477 | active |
| POLR2A | TargetFinder | ENCODE | GSE31477 | active |
| RAD21 | TargetFinder | ENCODE | GSE31477 | other |
| RB1 | TargetFinder | Chicas et al. | GSE19899 | repressive |
| RBL2 | TargetFinder | Chicas et al. | GSE19899 | repressive |
| RCOR1 | TargetFinder | ENCODE | GSE31477 | repressive |
| RELA | TargetFinder | Jin et al. | GSE43070 | active |
| RFX5 | TargetFinder | ENCODE | GSE31477 | active |
| YAP1 | TargetFinder | Stein et al. | GSE61852 | other |

(B)

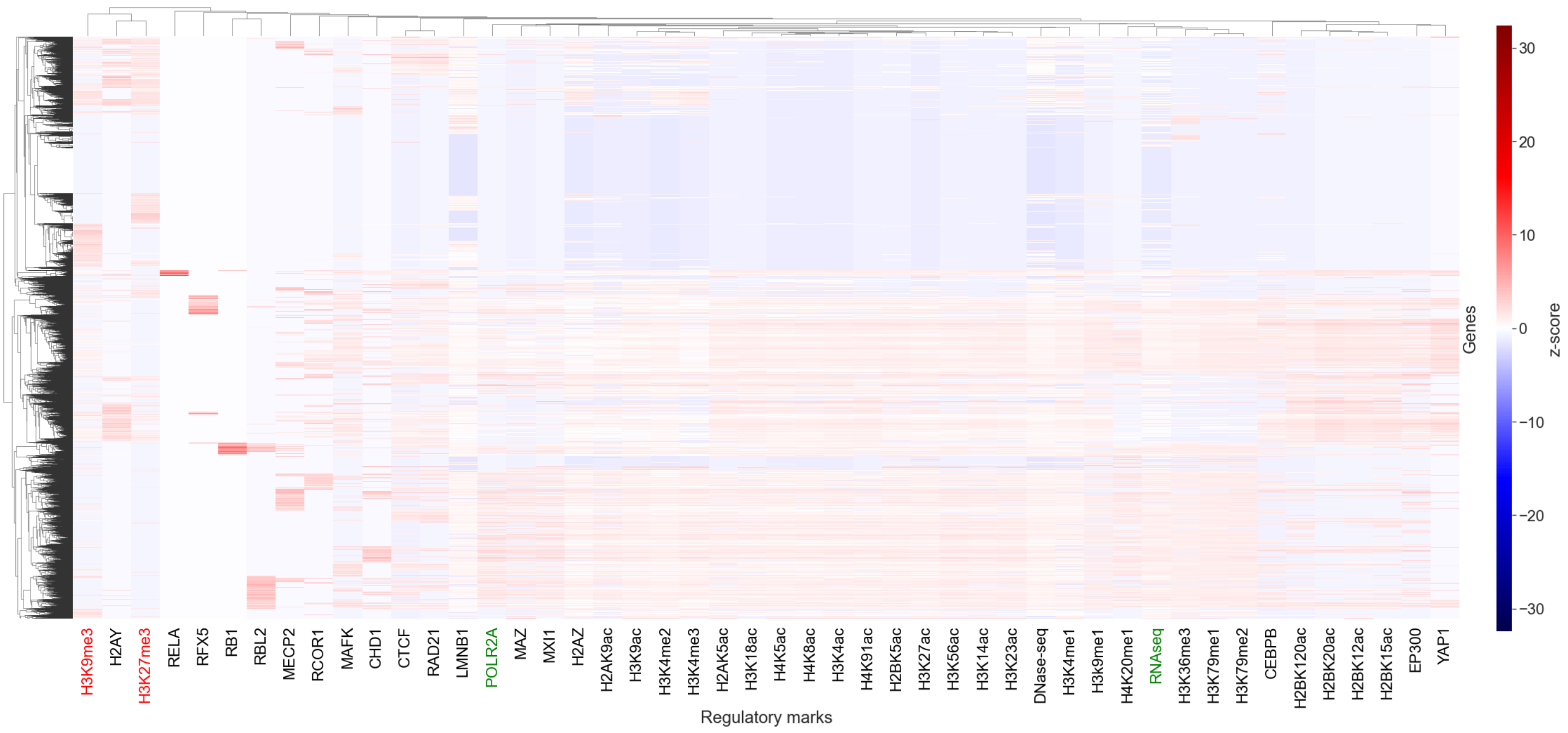


### Supplementary Figure 3

**Supplementary Fig. 3: Genome-wide regulatory marks in IMR-90 cells.** (**A**) Table describing the 48 regulatory features used in this study, including bulk RNA-seq and 47 cistromic marks (TF and histone ChIP-seq, DNAse-seq). The table also reports the source and accession ID of each data set. Likely annotations on the regulatory activity of each mark are provided when available in the literature (see [2] and [3]). (**B**) Genome-wide heatmap in IMR-90 cells of the 48 regulatory marks in (A). Genes (rows) and regulatory marks (columns) were clustered using hierarchical clustering with average linkage and Pearson correlation metric. The feature matrix was normalized and scaled prior to clustering [3]. The labels of two typical active marks (RNA-seq, POLR2A ChIP-seq) are colored in green and the labels of two typical repressive marks (H3K9me3, H3K27me3 ChIP-seq) are colored in red.

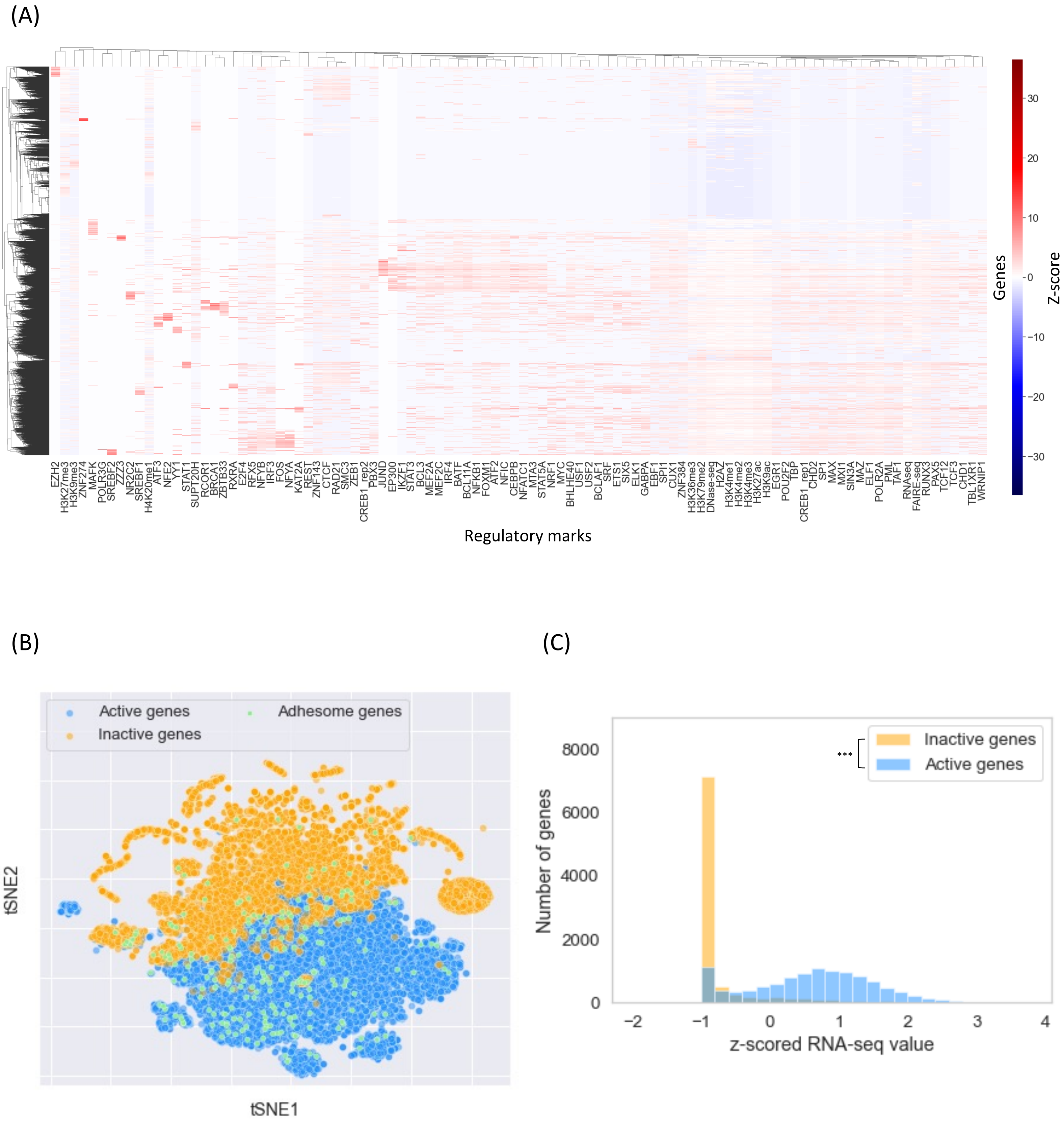


### Supplementary Figure 4

**Supplementary Fig. 4: Activity of adhesome genes in GM12878.** (**A**) Heatmap of 100 regulatory marks used to determine the activity of genes in GM12878 [3]. Genes (rows) and regulatory marks (columns) were clustered using hierarchical clustering with average linkage and Pearson correlation metric. The feature matrix was normalized and scaled prior to clustering [3]. (**B**) tSNE representation of all 25,959 genes based on the 100 GM12878 regulatory marks. Orange dots correspond to inactive genes (9,107 genes, as determined by the clustering in (A)), blue dots correspond to active genes (10,984 genes), and green dots correspond to adhesome genes. Out of the 222 adhesome genes studied (green dots), 172 are active (in contrast to 202 in IMR-90 cells). (**C**) Histogram of bulk RNA-seq expression in GM12878 for all active genes (blue) and all inactive genes (orange). Active genes have a significantly higher expression than inactive genes ($p$-value < 3e-308, Wilcoxon rank-sums test).

(A)

| Pathways | GO annotation | Total number of genes | Number of genes active in GM12878 |
|---|---|---|---|
| B cell receptor signaling | 0050853 | 81 | 63 |
| NF-kB signaling | | 104 | 66 |
| MAPK signaling | 0000165 | 218 | 121 |
| Actin cytoskeleton regulation | 0030036 | 218 | 135 |
| Calcium-mediated signaling | 0019722 | 240 | 103 |
| PI3-AKT signaling | 0043491 | 354 | 190 |

(B)

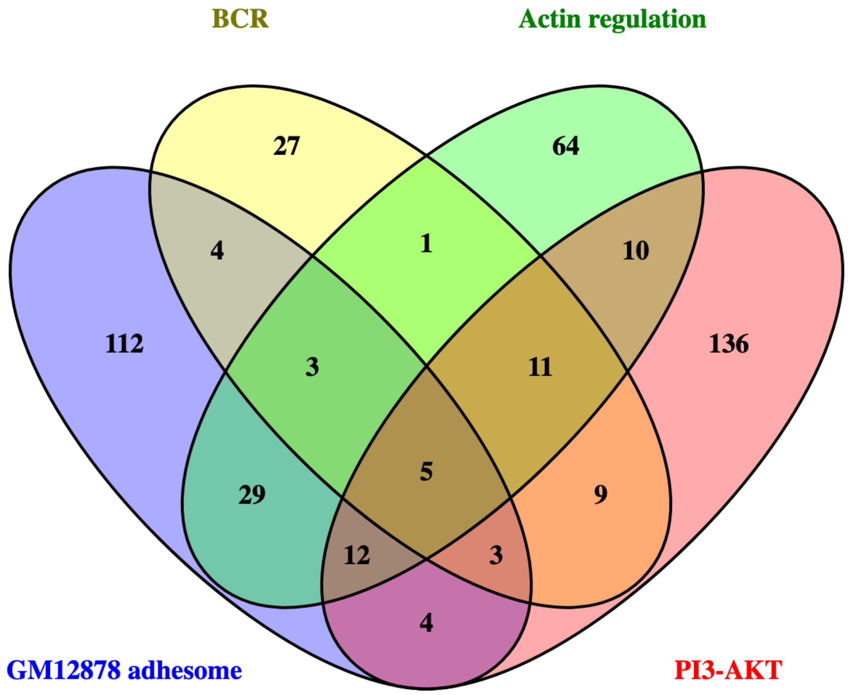


**Supplementary Figure 5**

**Supplementary Fig. 5: Active adhesome genes intersect with B Cell Receptor signaling pathways in GM12878.** (**A**) Table of signaling pathways related to the B Cell Receptor (BCR) signaling pathway. We intersect genes in each pathway with the active genes in GM12878 (as determined in Supplementary Fig. 4) to obtain the corresponding number of active pathway genes. (**B**) Venn diagram of four gene lists: active adhesome genes in GM12878 (blue), active BCR signaling genes (yellow), active actin cytoskeleton regulation genes (green), and active PI3-AKT signaling pathway genes (red). The Venn diagram was drawn using Venny [4].

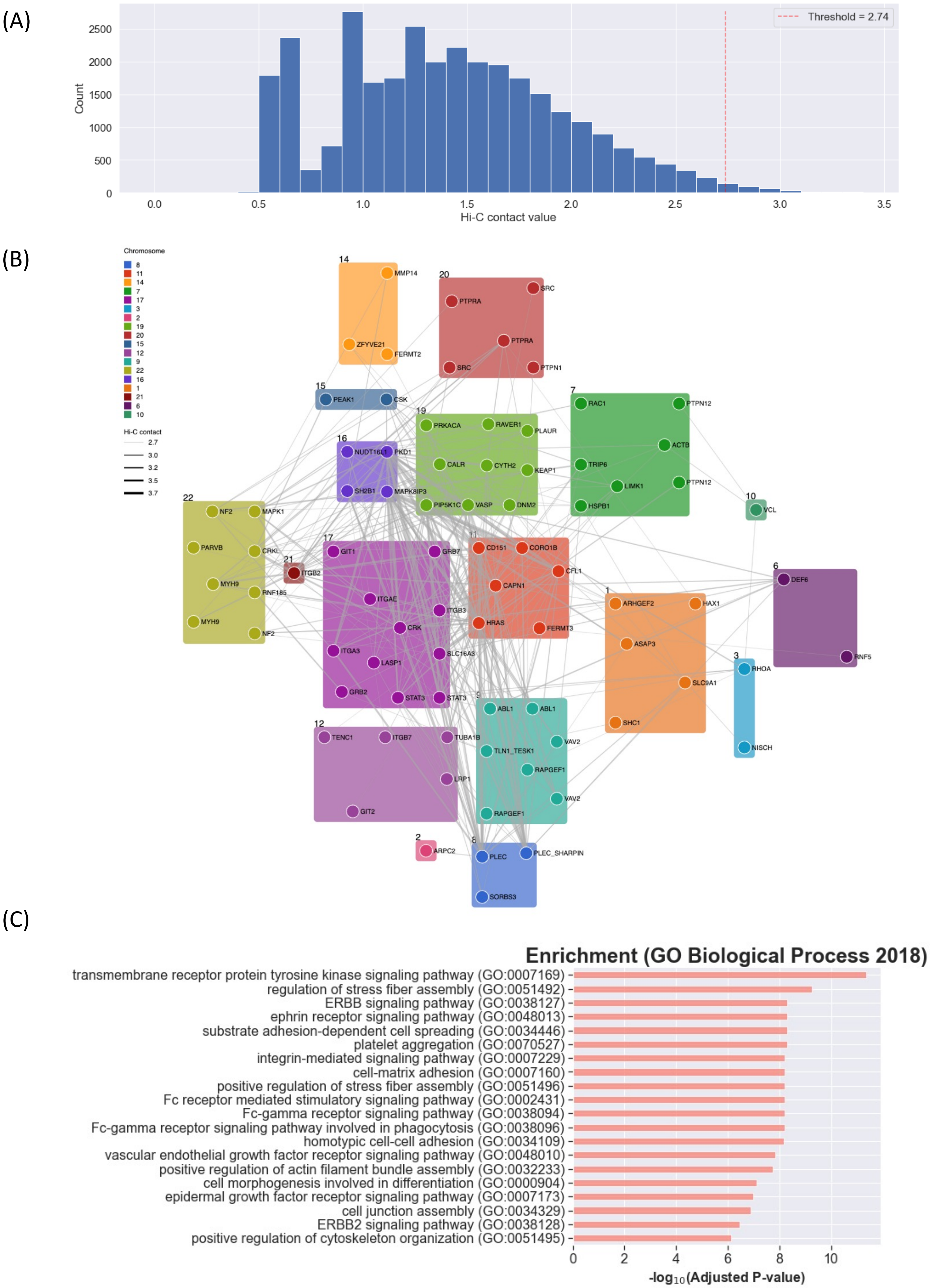


**Supplementary Figure 6**

**Supplementary Fig. 6: Active adhesome genes with high Hi-C contacts.** (**A**) Distribution of processed interchromosomal Hi-C contact values among active adhesome gene loci. A lower threshold materialized by the dashed red vertical line at 2.74 (99th percentile) is applied to select the strongest loci interactions, which are shown in the network in B. (**B**) Subnetwork of active adhesome gene loci, colored by chromosomal location and labeled with the active adhesome genes they harbor. Only non-isolated nodes engaging in high Hi-C contacts (Hi-C > 2.74, see A) are shown in the plot; edges are sized according to the corresponding Hi-C contact value. The network contains 82 active adhesome loci (74 adhesome genes) and 321 edges (average degree = 7.8). (**C**) Gene set enrichment analysis of active adhesome genes present in the subnetwork in B using the GO Biological Process 2018 gene sets. Only GO terms with FDR-corrected enrichment *p*-value smaller than 0.0001 are reported.

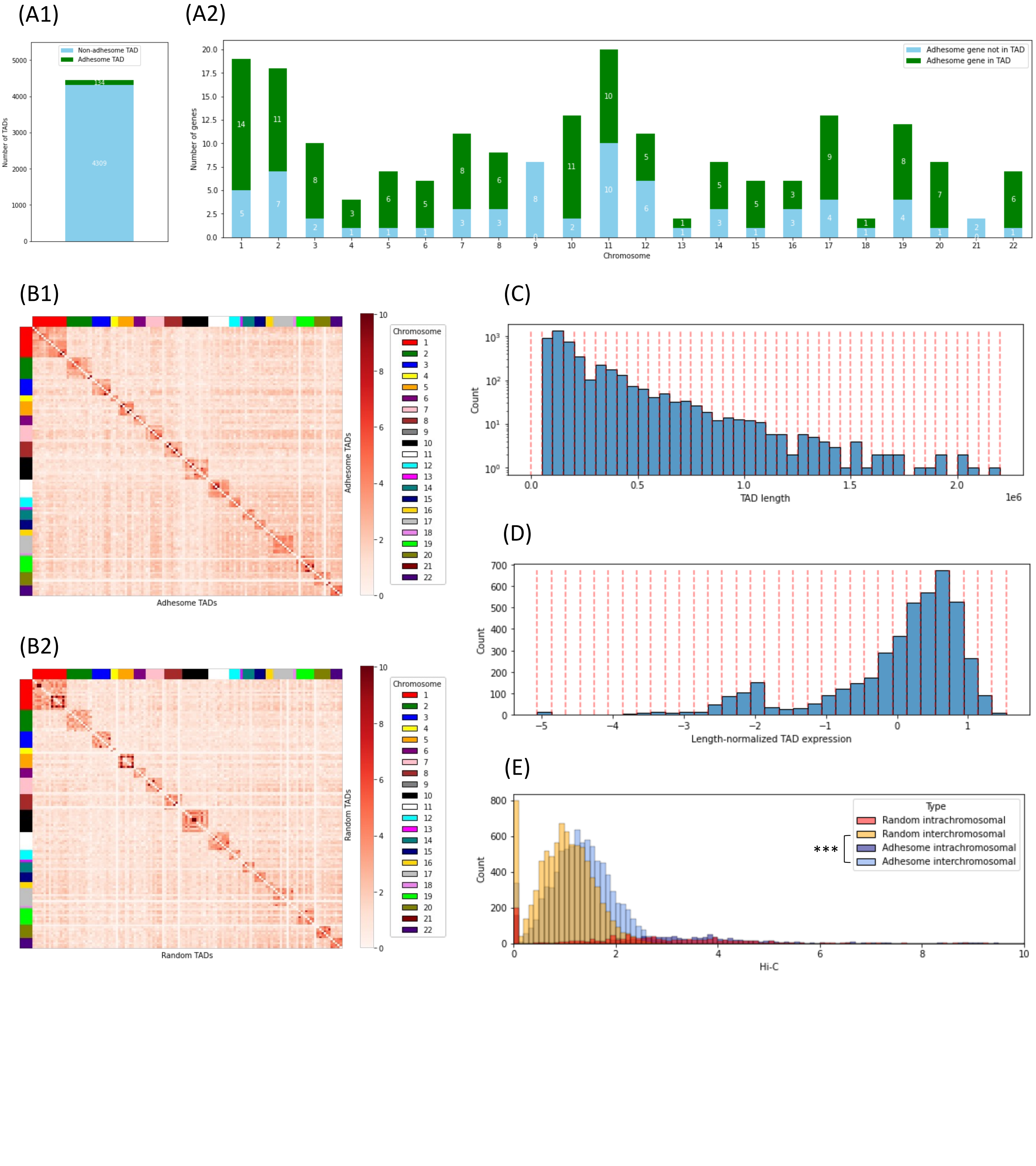


**Supplementary Figure 7**

**Supplementary Fig. 7: TAD-based interchromosomal proximity analysis.** (**A**) 134 TADs out of 4,443 TADs called by Arrowhead (at resolution 5 kb with Knight-Ruiz normalization) contain at least one active adhesome gene (A1). The number of active adhesome genes either in TADs or not in TADs per chromosome is shown in A2. (**B**) Hi-C contact map between active adhesome TADs (C1) and random active non-adhesome TADs (C2). Each row/column corresponds to a TAD. The contact value between two TADs is obtained by averaging the Hi-C contact values associated with these TADs. TADs are grouped by chromosomal location, as shown by the side color bars. The selection of random active adhesome TADs is explained in D and E. (**C**) Distribution of TAD length for all 4,443 TADs called by Arrowhead at resolution 5 kb with Knight-Ruiz normalization. TADs are stratified into buckets of length 50 kb. These buckets are used to sample background TADs on a per-chromosome basis. (**D**) Distribution of length-normalized TAD gene expression for all 4,443 TADs called by Arrowhead at resolution 5 kb with Knight-Ruiz normalization. TADs are stratified into buckets of size 0.2. These buckets are used to sample background TADs on a per-chromosome basis. (**E**) Distribution of interchromosomal and intrachromosomal contacts between active adhesome TADs and between random TADs. Random TADs were selected such that they are similar in size to adhesome TADs, they are similarly distributed over the chromosomes, and they have a similar expression level. The distribution of interchromosomal contacts between active adhesome TADs significantly dominates the distribution of interchromosomal contacts between random TADs (Wilcoxon Rank-Sum test, $p$-value < 5e-124).

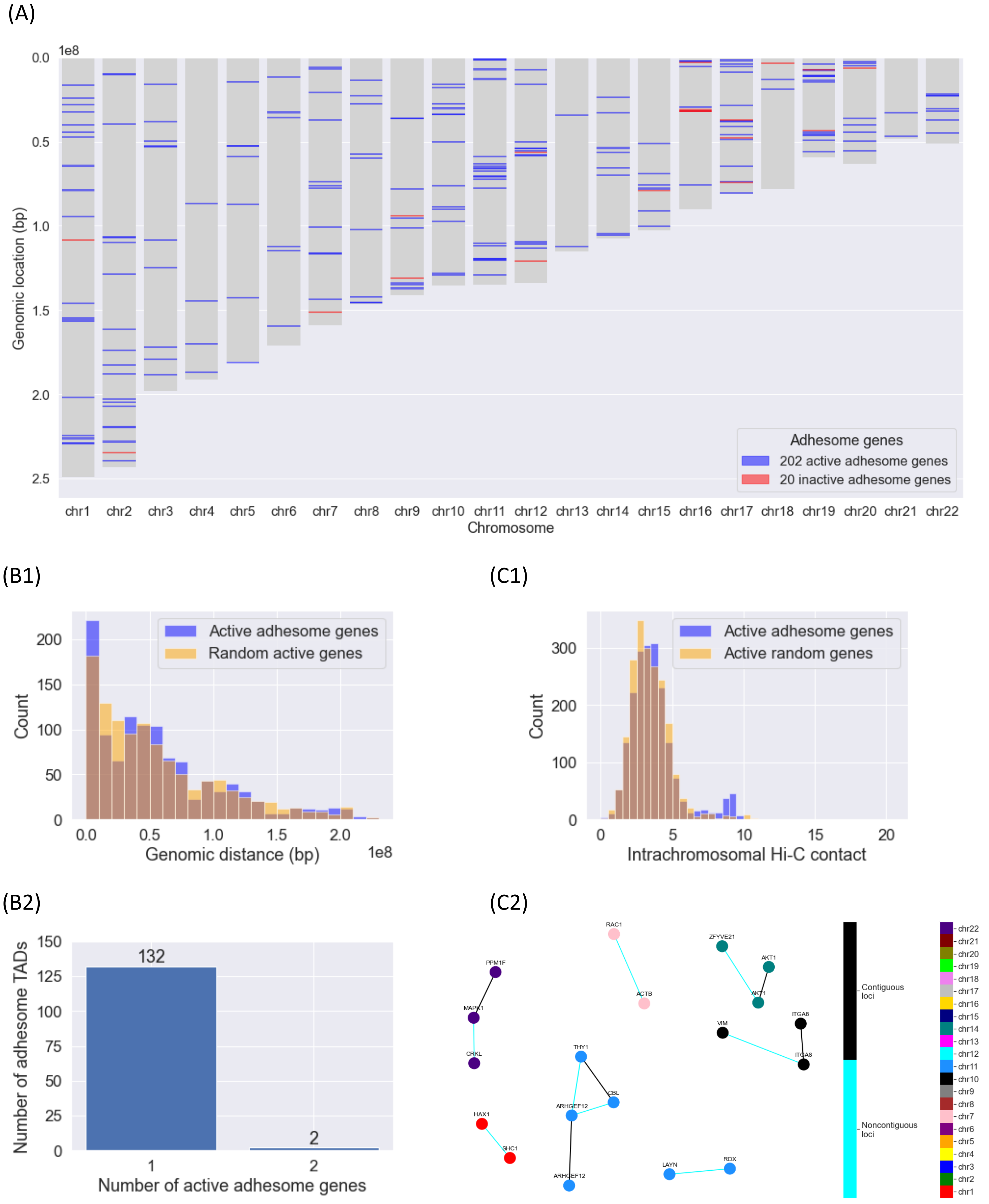


**Supplementary Figure 8**

**Supplementary Fig. 8: Intrachromosomal proximity analysis.** (**A**) Localization of active and inactive adhesome genes on the chromosomes (using the hg19 reference genome). (**B**) B1: Distribution of 1D genomic distances (in bp) between active adhesome genes and between a set of random active genes located on the same chromosomes; the distance between 2 genes (located on the same chromosome) is defined as the minimum distance between |start2-end1| and |start1-end2|. B2: Number of active adhesome genes per active adhesome TAD, using TADs called by Arrowhead at resolution 5 kb with Knight-Ruiz normalization. (**C**) C1: Distribution of intrachromosomal Hi-C contacts among active adhesome gene loci and random active non-adhesome loci located on the same chromosomes. C2: Subnetwork of active adhesome loci whose intrachromosomal Hi-C contact values are above 7 in C1; only non-degenerate connected components are shown. Adhesome gene loci are colored by their chromosomal location and labeled with the adhesome genes they harbor. Contiguous loci are linked by black edges, while non-contiguous loci are linked by cyan edges.

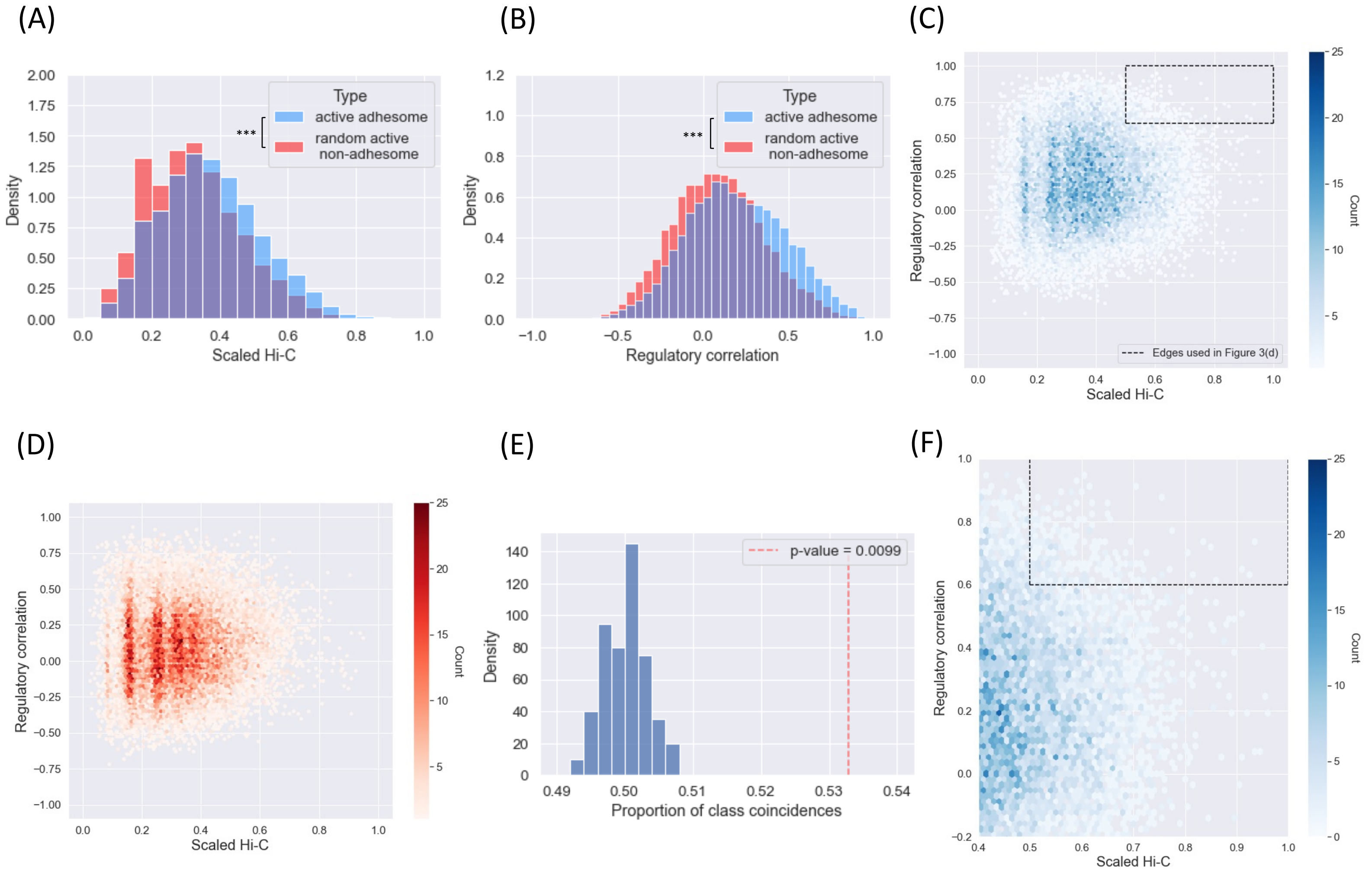


**Supplementary Figure 9**

**Supplementary Fig. 9: Joint distribution of scaled Hi-C contact values and regulatory correlation among active adhesome genes in IMR-90 cells.** (**A**) Histogram of scaled Hi-C contact values between active adhesome genes (blue) and a random set of active non-adhesome genes (red). There are significantly more contacts between adhesome genes compared to random non-adhesome genes (Wilcoxon Rank-Sums test, *p*-value = 1e-85). (**B**) Histogram of regulatory Pearson correlation (based on the 48 regulatory marks of Supplementary Fig. 3A) between active adhesome genes (blue) and a random set of active non-adhesome genes (red). There is significantly more coregulation among adhesome genes compared to random non-adhesome genes (Wilcoxon Rank-Sums test, *p*-value < 3e-308). (**C**) 2D histogram of scaled Hi-C contact values and Pearson regulatory correlation for active adhesome genes. The dashed box shows the thresholds used to build the network in Fig. 3D. (**D**) 2D histogram of scaled Hi-C contact values and Pearson regulatory correlation for random active non-adhesome genes. (**E**) Simulated null distribution for the permutation test used to assess the difference between the joint distribution of scaled Hi-C contact values and Pearson regulatory correlation for the two gene groups of interest: active adhesome genes (blue, see (C)) and the randomly selected set of active non-adhesome genes (red, see (D)). The test is a two-sample test based on the number of nearest neighbor type co-incidences [3]. The red vertical dashed line corresponds to the actual value of the statistic. (**F**) Zoom into the region around the dashed box of (C).

(A)

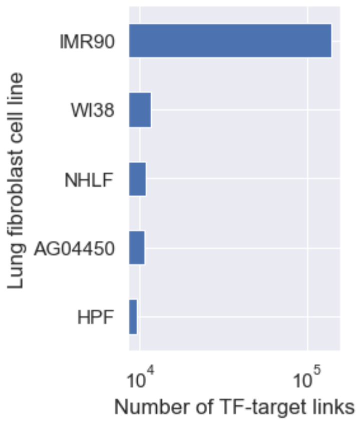


(B)

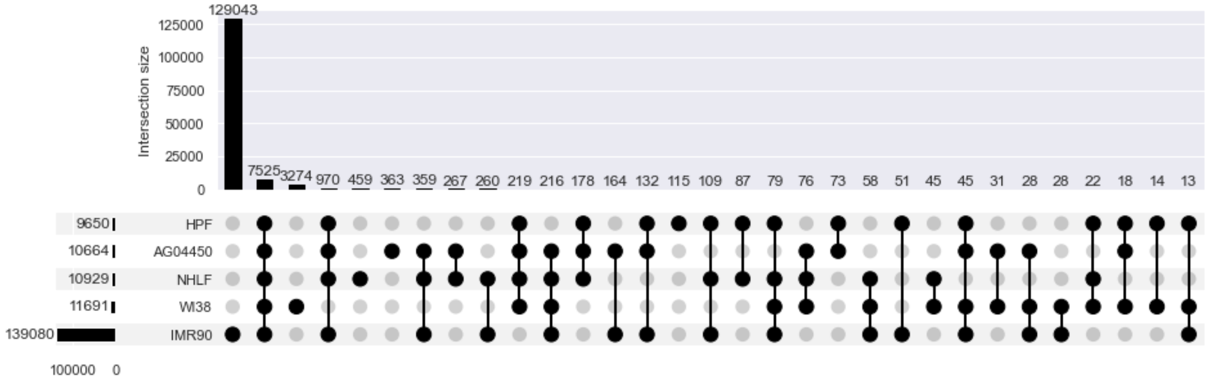


(C)

| Cell line | TFs<br>red = inactive ; purple = not present in IMR90 data;<br>brown = target inactive adhesome gene | Number of TFs | Number of active TFs |
|---|---|---|---|
| AG04450 | CTCF | 1 | 1 |
| HPF | CTCF | 1 | 1 |
| IMR90 | BDP1, BRCA1, BRF1, BRF2, CEBPB, **CHD1**, CTCF, E2F7, EP300, GTF2B, LMNB1, MAFK, MAZ, MXI1, POLR2A, RAD21, RB1, RBL1, RBL2, RCOR1, RFX5, **SNAPC1**, SNAPC2, **SNAPC4**, **SNAPC5**, **SPI1**, TP53 | 27 | 24 |
| NHLF | CTCF | 1 | 1 |
| WI38 | **SUMO2, UBE2I, PIAS4, SUMO1**, CTCF | 5 | 5 |

(D)

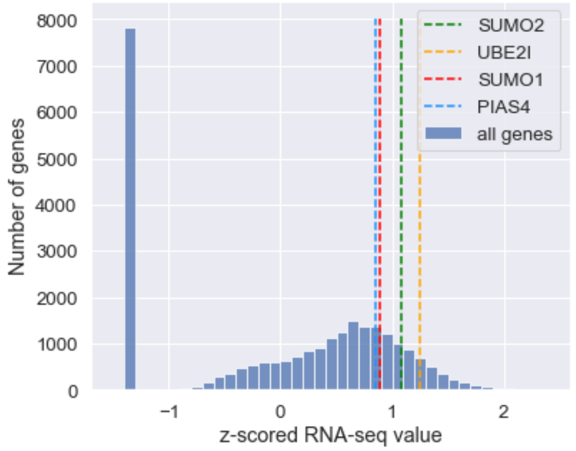


(E)

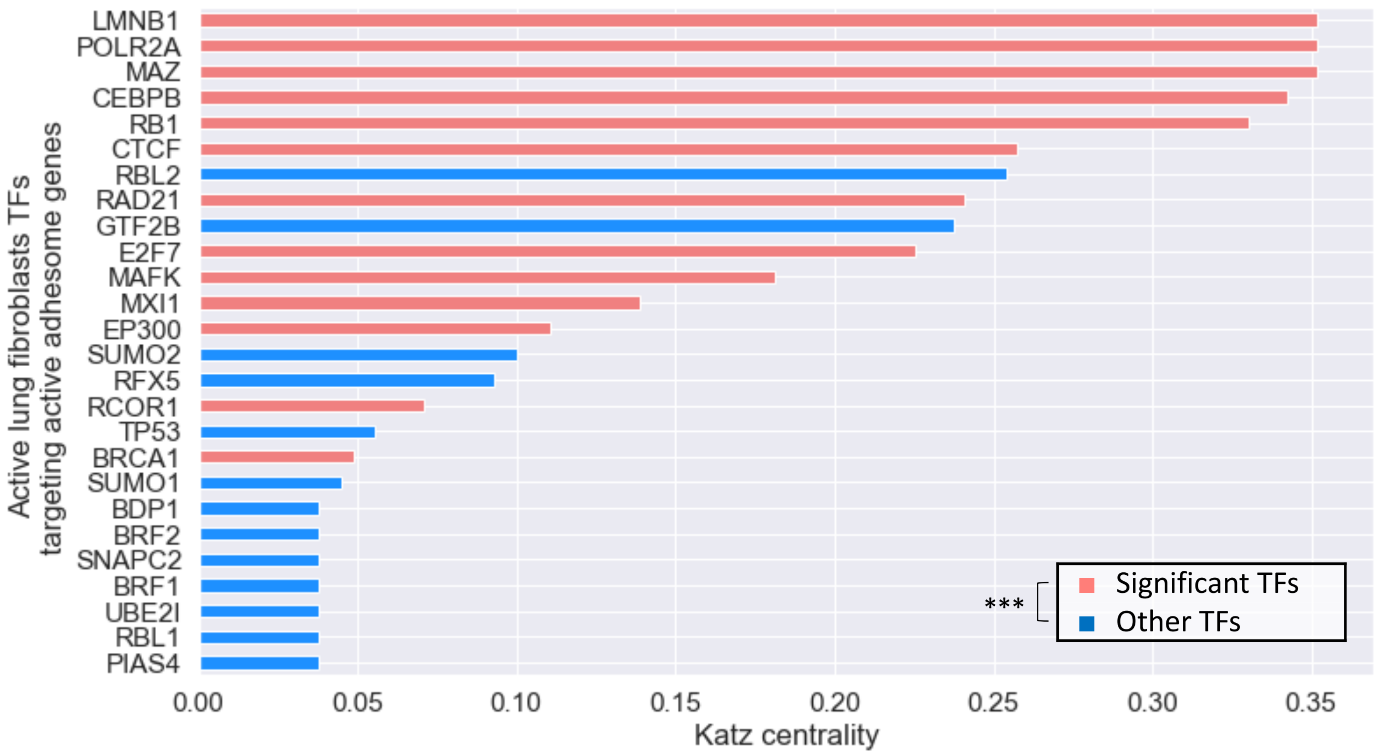


**Supplementary Figure 10**

**Supplementary Fig. 10: TF-target data in normal human lung fibroblast cell lines.** (**A**) Bar plot of the number of TF-target links from the hTFtarget data set in five human lung fibroblast cell lines (AG04450, HPF, NHLF, WI38, IMR-90). Only targets that are active in IMR-90 are considered. (**B**) Set overlap plot for the TF-target relationships in cell lines AG04450, HPF, NHLF, WI38, and IMR-90. Only targets that are active in IMR-90 are considered. While most identified TF-target interactions come from the IMR-90 data, other human lung fibroblast cell lines contribute a few additional TF-target interactions that are not present in the IMR-90 data. (**C**) Table of TFs present for all normal human lung fibroblast cell lines. Red font indicates inactive genes, purple font indicates TFs that are not present in IMR-90, and brown font indicates TFs that target inactive adhesome genes. (**D**) Histogram of z-scored RNA-seq expression for all genes in IMR-90. The expression of SUMO2, SUMO1, UBE2I and PIAS4 (transcription factors found in WI38 only) are represented with colored vertical dashed lines. (**E**) Bar plot of Katz centrality of all active lung fibroblast TFs in the network of Fig. 4E. The parameter alpha of Katz centrality was chosen to be 1/s-0.01, where s is the largest eigenvalue (in magnitude) of the adjacency matrix of the network in Fig. 4E. Bars are colored in red if they correspond to significant TFs from Fig. 4D, otherwise they are colored in blue. There is a significant enrichment of significant TFs at the top of the list (XL-minimal hypergeometric test [5], *p*-value < 6e-4 and optimal cutoff value at 17, which corresponds to BRCA1). The parameters X and L were chosen to be 10% of the number of significant TFs and 150% of the number of significant TFs, respectively.

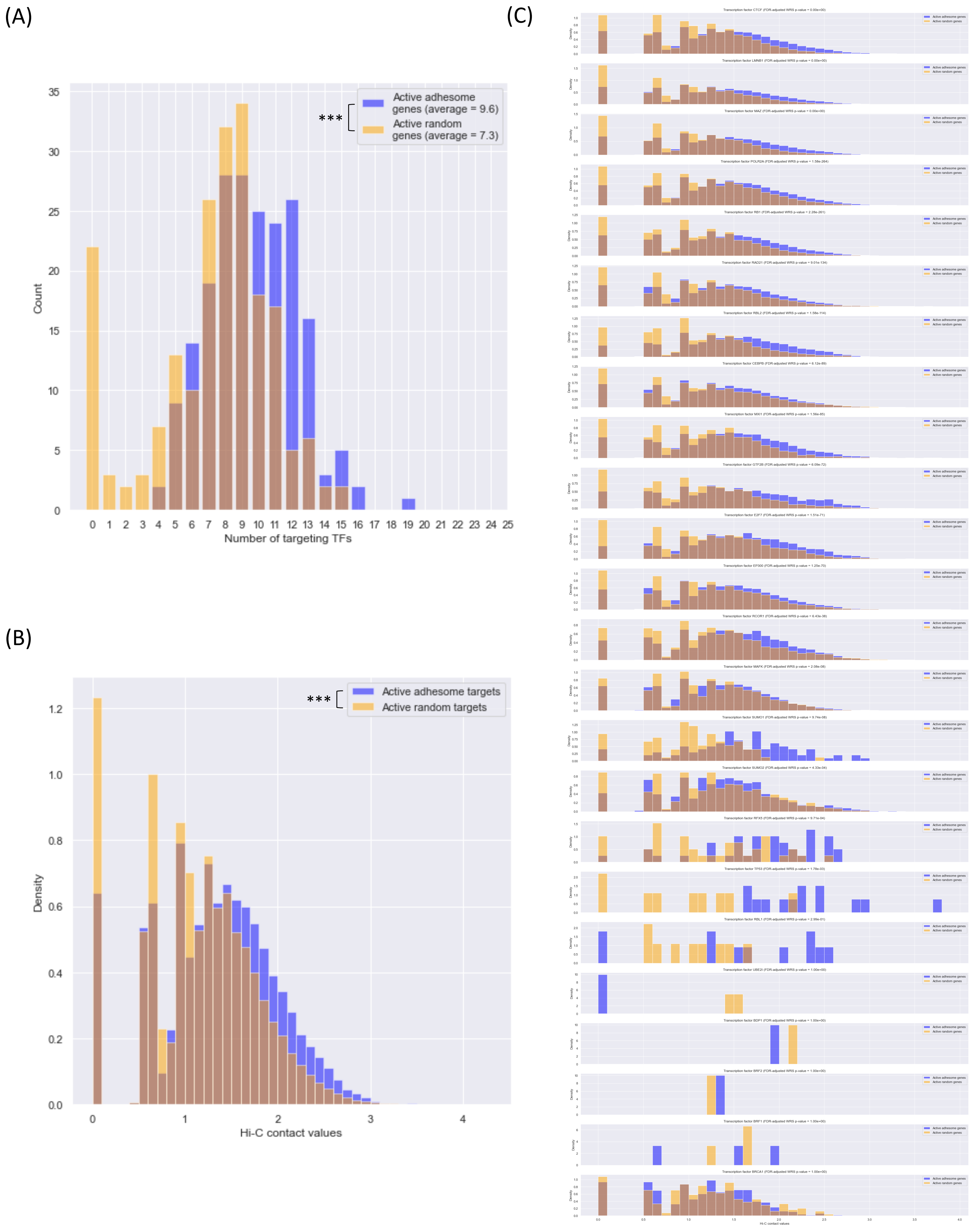


**Supplementary Figure 11**

**Supplementary Fig. 11: Spatial proximity analysis of transcription factor targets.** (**A**) Histograms of the number of TFs targeting active adhesome genes and the number of TFs targeting a set of random active non-adhesome genes (Wilcoxon Rank-Sum *p*-value < 2e-10). (**B**) Histograms of interchromosomal Hi-C contacts among active adhesome TF targets and random active non-adhesome non-TF target genes located on the same chromosomes (Wilcoxon Rank-Sum test, *p*-value < 3e-308). (**C**) Histograms of interchromosomal Hi-C contacts among active adhesome TF targets and among random active non-adhesome non-TF target genes located on the same chromosomes, separately for each TF. The plots are organized in ascending order of Wilcoxon Rank-Sum *p*-value.

(A)

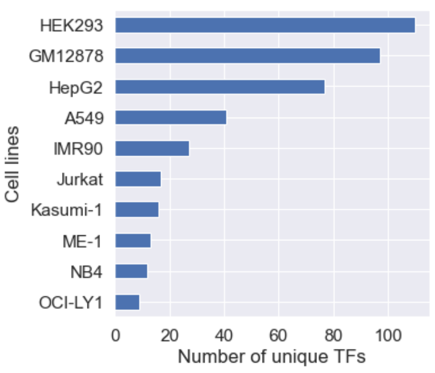


(B)

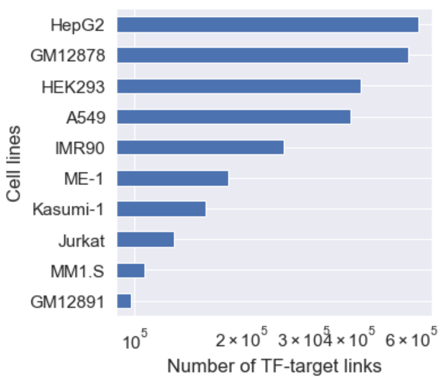


(C)

| Lung fibroblast adhesome TF | Proportion of adhesome genes targeted | | | | |
|---|---|---|---|---|---|
| | Lung fibroblasts | GM12878 | HEK293 | A549 | HepG2 |
| LMNB1 | 1.00 | 0.00 | 0.00 | 0.00 | 0.00 |
| MAZ | 0.94 | 0.43 | 0.00 | 0.00 | 0.80 |
| POLR2A | 0.89 | 0.84 | 0.00 | 0.91 | 0.88 |
| CTCF | 0.86 | 0.86 | 0.99 | 0.91 | 0.91 |
| CEBPB | 0.86 | 0.02 | 0.00 | 0.66 | 0.50 |
| RB1 | 0.77 | 0.00 | 0.00 | 0.00 | 0.00 |
| SPI1 | 0.73 | 0.81 | 0.00 | 0.00 | 0.00 |
| RAD21 | 0.73 | 0.64 | 0.00 | 0.64 | 0.97 |
| CHD1 | 0.55 | 0.00 | 0.00 | 0.00 | 0.00 |
| EP300 | 0.54 | 0.77 | 0.00 | 0.41 | 0.45 |
| MAFK | 0.46 | 0.01 | 0.00 | 0.00 | 0.46 |
| RCOR1 | 0.37 | 0.01 | 0.00 | 0.00 | 0.16 |
| E2F7 | 0.36 | 0.00 | 0.00 | 0.00 | 0.00 |
| RBL2 | 0.35 | 0.00 | 0.00 | 0.00 | 0.00 |
| MXI1 | 0.34 | 0.14 | 0.00 | 0.00 | 0.59 |
| GTF2B | 0.29 | 0.00 | 0.00 | 0.00 | 0.00 |
| SUMO2 | 0.21 | 0.00 | 0.00 | 0.00 | 0.00 |
| BRCA1 | 0.15 | 0.03 | 0.00 | 0.00 | 0.08 |
| SUMO1 | 0.05 | 0.00 | 0.00 | 0.00 | 0.00 |
| RFX5 | 0.04 | 0.03 | 0.00 | 0.00 | 0.21 |
| RBL1 | 0.02 | 0.00 | 0.00 | 0.00 | 0.00 |
| TP53 | 0.02 | 0.00 | 0.00 | 0.00 | 0.00 |
| BRF1 | 0.01 | 0.00 | 0.00 | 0.00 | 0.00 |
| UBE2I | 0.01 | 0.00 | 0.00 | 0.00 | 0.00 |
| BDP1 | 0.01 | 0.00 | 0.00 | 0.00 | 0.00 |
| BRF2 | 0.01 | 0.00 | 0.00 | 0.00 | 0.00 |
| SNAPC2 | <0.01 | 0.00 | 0.00 | 0.00 | 0.00 |
| SNAPC4 | <0.01 | 0.00 | 0.00 | 0.00 | 0.00 |
| PIAS4 | <0.01 | 0.00 | 0.00 | 0.00 | 0.00 |

(D)

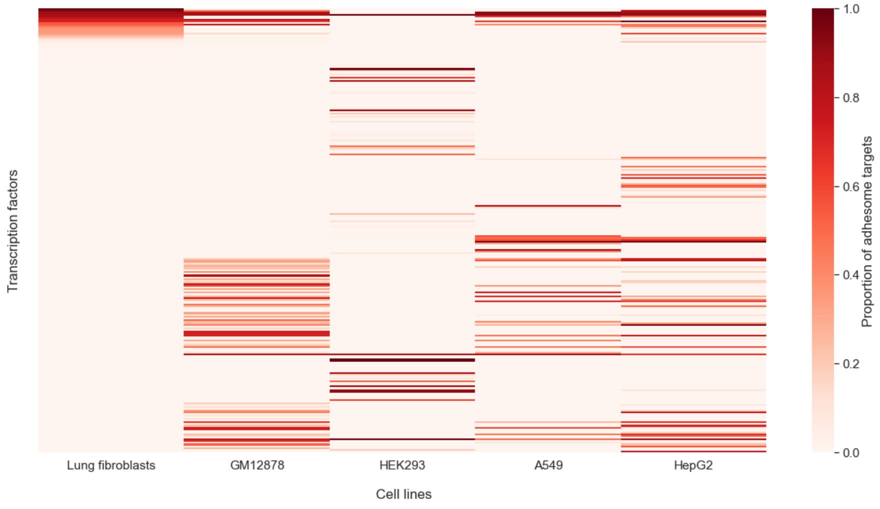


### Supplementary Figure 12

**Supplementary Fig. 12: The identified active human lung fibroblast TFs target fewer adhesome genes in other cell lines and are quite specific to human lung fibroblasts.** (**A**) Number of unique TFs present in the hTFtarget data set for different adherent and non-adherent cell lines. (**B**) Number of unique TF-target links present in the hTFtarget data set for different cell lines. The cell lines HepG2, GM12878, HEK293, A549 and normal lung fibroblasts (combination of IMR-90, WI38, AG04450, HPF and NHLF) have the highest amount of data. (**C**) Table showing the proportion of adhesome genes targeted by selected lung fibroblast TFs in different cell lines, including lung fibroblasts, HepG2, GM12878, HEK293, and A549. Rows shaded in red correspond to the 13 significant TFs identified in lung fibroblasts. (**D**) Heatmap of the proportion of adhesome genes targeted by TFs in different cell lines compared to the proportion of adhesome genes targeted in normal lung fibroblasts. These include cell lines with adherence properties (A549, HepG2, HEK293) and a non-adherent cell line (GM12878). TFs that target (active and/or inactive) adhesome genes in IMR-90 are in the top 29 rows. The remaining TFs do not target adhesome genes in IMR-90 but target at least one adhesome gene in other cell lines.

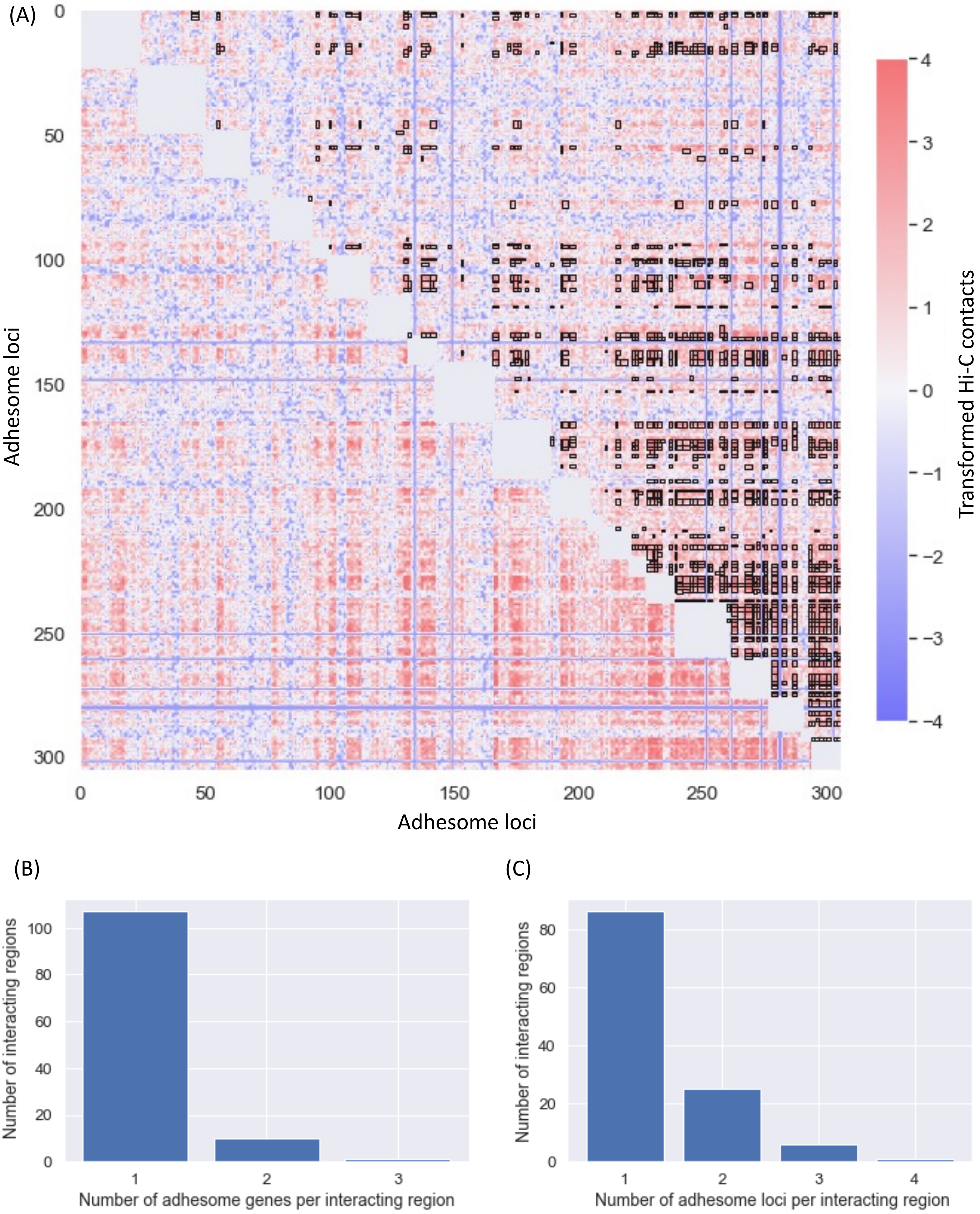


**Supplementary Figure 13**

**Supplementary Fig. 13: Projection of LAS submatrices onto adhesome loci.** (**A**) Heatmap of transformed Hi-C contact frequencies among active adhesome and adhesome TF loci (frequencies were log-transformed and centerdized using the mean and standard deviation of contact values across the entire genome). Significant submatrices identified by the LAS algorithm and whose average Hi-C value is higher than 2 are shown as black boxes. This heatmap is obtained by combining LAS interacting regions containing adhesome genes from all interchromosomal Hi-C contact matrices (such as, for example, the matrix in Fig. 5A). (**B**) Bar plot of the number of adhesome genes per adhesome interacting region. Adhesome LAS interacting regions contain at most 3 adhesome genes. (**C**) Bar plot of the number of adhesome loci per adhesome interacting region. Adhesome LAS interacting regions contain at most 4 adhesome loci. To relate this to (B), note that an adhesome gene may span multiple 250 kb loci.

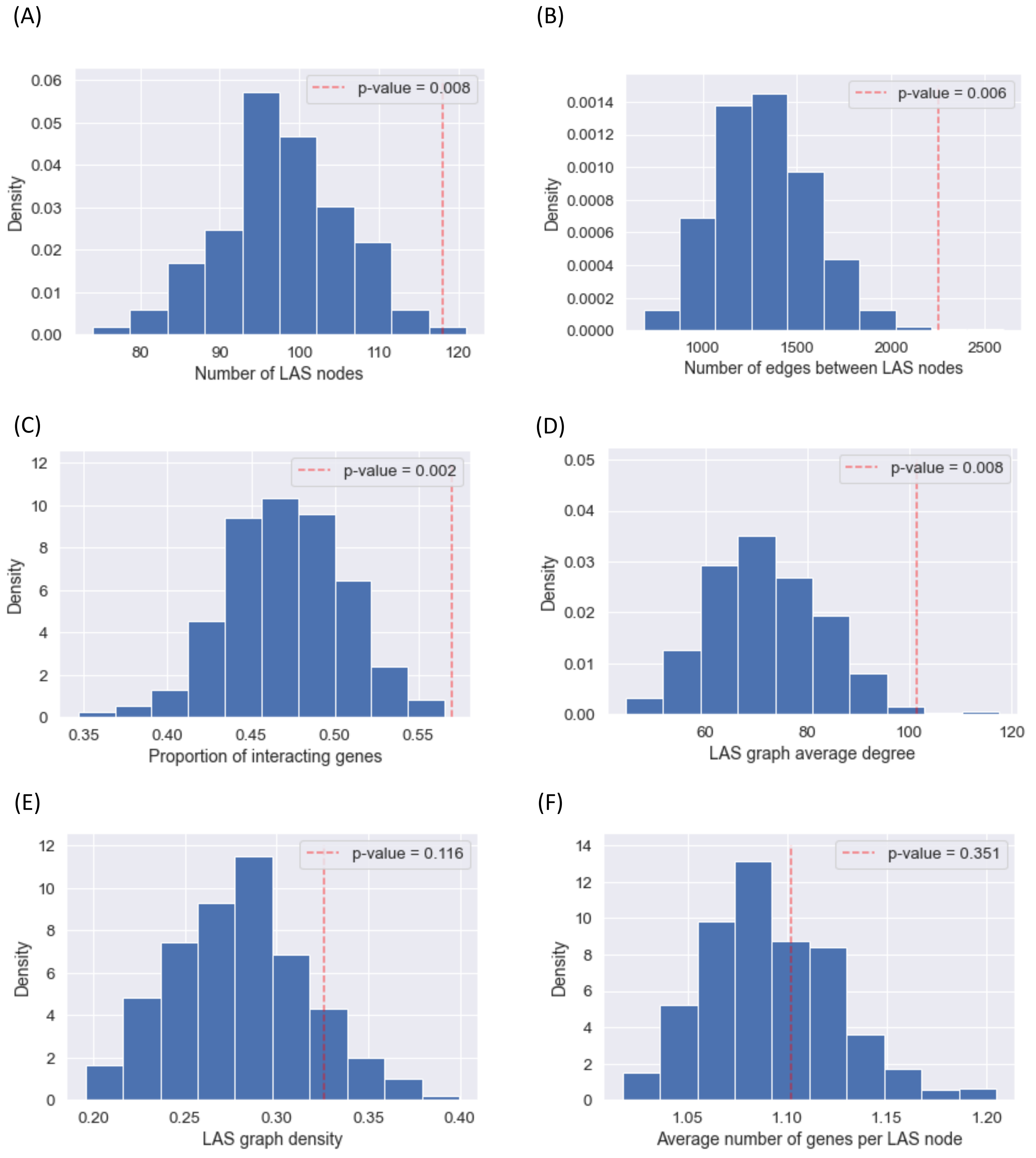


**Supplementary Figure 14**

**Supplementary Fig. 14: Statistical significance of the LAS adhesome network.** (**A**) Histogram of the number of LAS nodes containing at least one gene from a collection of 500 random gene sets. Each random gene set is sampled uniformly without replacement among all active non-adhesome genes. The number of nodes containing active adhesome and adhesome TF genes is shown with a red dashed vertical line, corresponding to a *p*-value of 0.008. This shows that a significantly high number of nodes contain adhesome genes under the 1% significance level. (**B**) Histogram of the number of interactions among LAS nodes containing at least one gene from the collection of 500 random gene sets in (A). The number of interactions among adhesome LAS nodes is shown with a red dashed vertical line, corresponding to a *p*-value of 0.006. This shows that there is a significantly high number of interactions between adhesome LAS nodes at the 1% significance level. (**C**) Histogram of the proportion of genes belonging to LAS nodes from the collection of 500 random gene sets in (A). The proportion of adhesome genes belonging to adhesome LAS nodes is shown with a red dashed vertical line, corresponding to a *p*-value of 0.002. This shows that there is a significantly high proportion of adhesome genes that belong to adhesome LAS nodes at the 1% significance level. (**D**) Histogram of average degree computed on networks of LAS nodes containing genes from the collection of 500 random gene sets in (A). The average degree of the adhesome LAS network is shown with a red dashed vertical line, corresponding to a *p*-value of 0.008. This shows that, on average, the degree of adhesome LAS nodes is significantly higher than the degree of LAS nodes containing random genes at the 1% significance level. (**E**) Histogram of graph density computed on networks of LAS nodes containing genes from the collection of 500 random gene sets in (A). The graph density of the adhesome LAS network is shown with a red dashed vertical line, corresponding to a *p*-value of 0.116. (**F**) Histogram of the average number of genes per LAS node from the collection of 500 random gene sets in (A); only LAS nodes containing at least one gene from the random gene set at hand are considered. The average number of adhesome genes per adhesome LAS node is shown with a red dashed vertical line, corresponding to a *p*-value of 0.351.

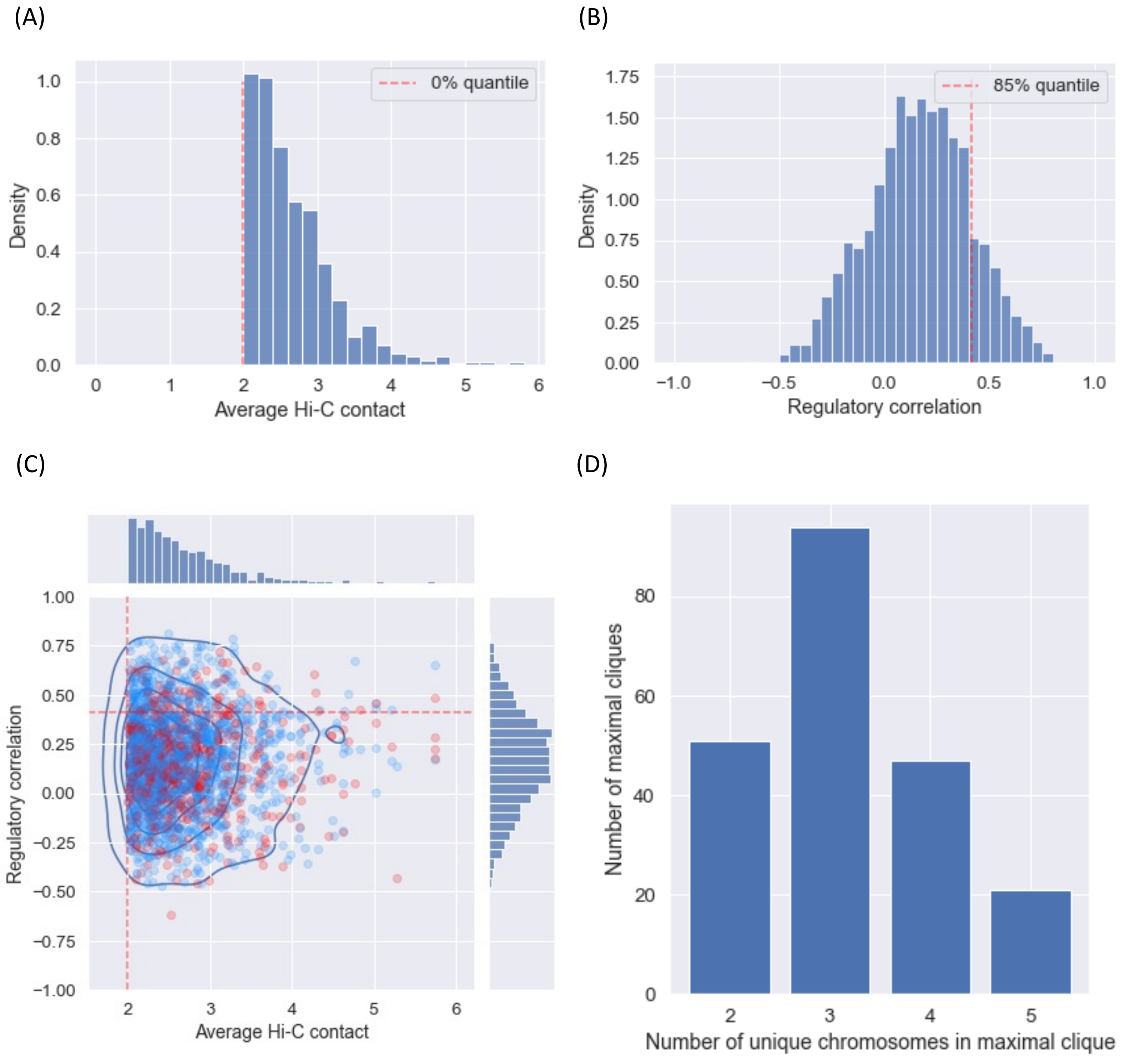


**Supplementary Figure 15**

**Supplementary Fig. 15: Regulatory correlation of adhesome LAS nodes.** (**A**) Histogram of average transformed Hi-C contact values among all adhesome LAS nodes. The red dashed vertical line corresponds to the minimum value, which is equal to 2 (by design from the LAS thresholding we performed in a previous step, see Supplementary Fig. 9C). (**B**) Histogram of average Spearman regulatory correlation among all adhesome LAS nodes. The average Spearman regulatory correlation between two adhesome LAS nodes is computed by taking the average of Spearman regulatory correlations between all tuples of adhesome genes in the two nodes, using the 48 regulatory marks used throughout this article [3]. The red dashed vertical line corresponds to the 85% quantile, used as a lower regulatory threshold to select interactions among adhesome LAS nodes for the adhesome LAS network of Fig. 5C. (**C**) Scatterplot and 2D contour plot of pairs of adhesome LAS nodes. The location of a given pair (u, v) on the scatterplot is determined by the average Hi-C contact value (x-axis) and the regulatory correlation (y-axis) associated with the edge u–v of the adhesome LAS network. Pairs that involve at least one TF targeting active adhesome genes are shown in red. The red dashed lines correspond to marginal thresholds on the distribution of average Hi-C contact values (vertical, 0% quantile) and regulatory correlations (horizontal, 85% quantile). (**D**) Bar plot of the number of unique chromosomes per maximal clique in the adhesome LAS network obtained after dropping interactions whose average Spearman regulatory correlation is below the threshold in (B). Maximal cliques contain at most 5 unique chromosomes, which is consistent with the hypothesis that no more than 5 chromosomes can physically interact at a given location.

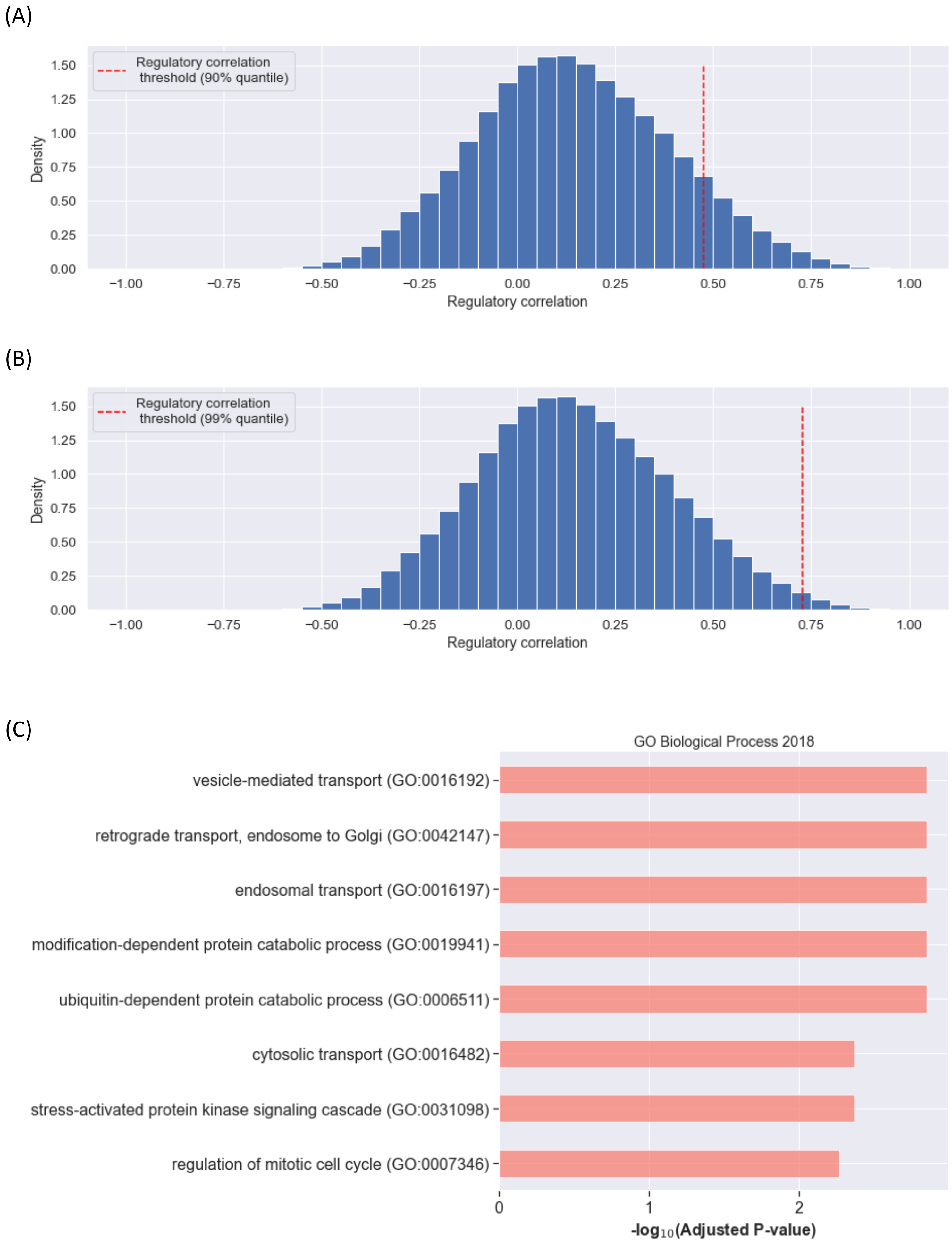


**Supplementary Figure 16**

**Supplementary Fig. 16: Gene set enrichment analysis of active non-adhesome genes interacting with active adhesome genes.** (**A**) Histogram of Spearman regulatory correlation between (i) active non-adhesome genes in adhesome interacting regions and (ii) the active adhesome genes with which these genes interact interchromosomally. Regulatory correlation is computed from the 48 regulatory marks used throughout this article [3]. The red dashed vertical threshold line marks the upper decile of the distribution. (**B**) Same as A, with the red dashed vertical threshold line marking the upper percentile of the distribution. (**C**) Significant Gene Ontology (GO) terms associated to active non-adhesome genes belonging to genomic regions that interact with adhesome interacting regions. Only genes having a strong positive regulatory correlation (upper percentile in B) with the active adhesome genes they interact with (in LAS) are considered. A GO term is reported as significant if its FDR-adjusted *p*-value is lower than 1e-2.

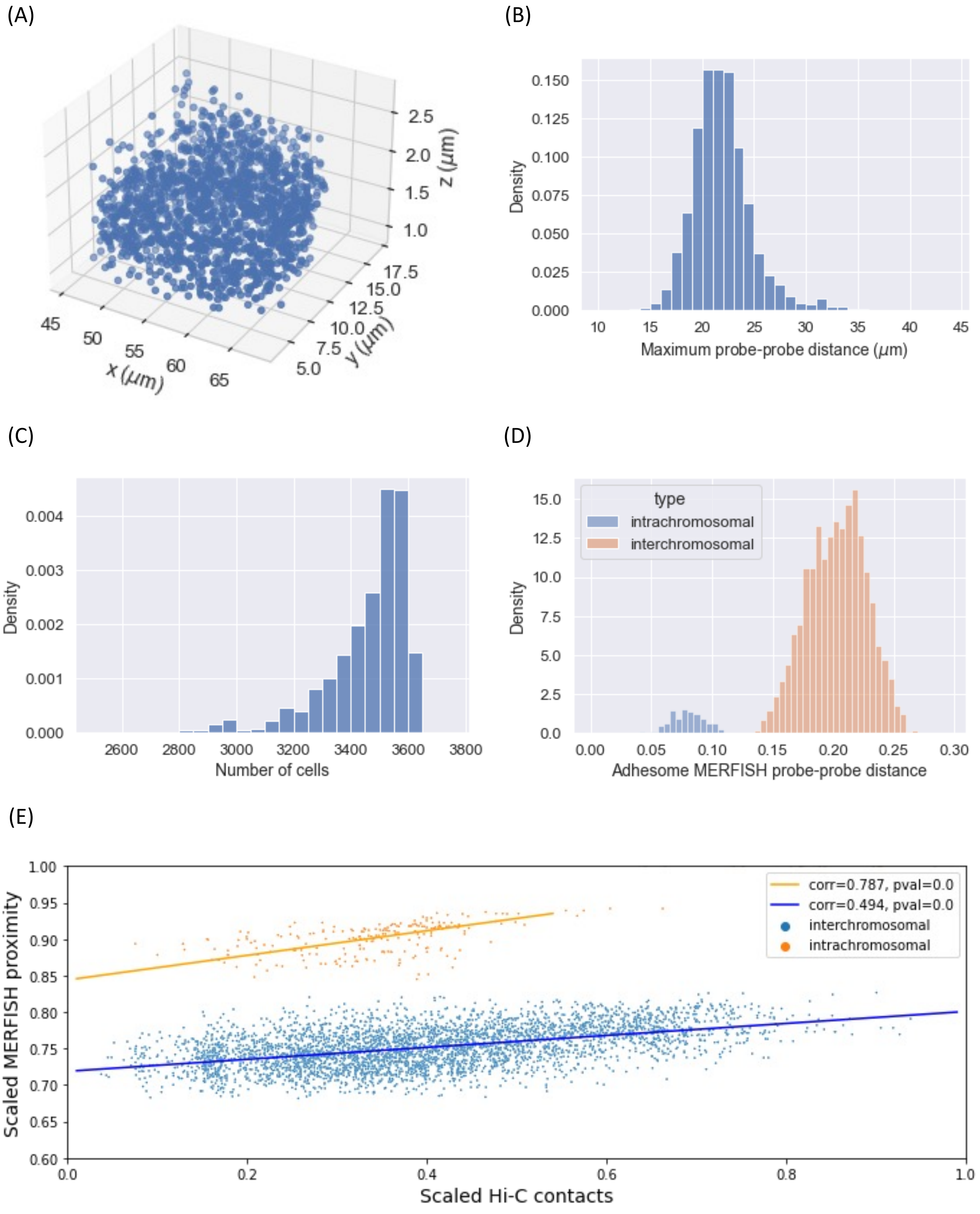


**Supplementary Figure 17**

**Supplementary Fig. 17: Normalization of Euclidean distances between DNA-MERFISH probes.** (**A**) 3D scatterplot of all 959 DNA-MERFISH probes in the nucleus of a given cell (cell ID 1788). The shape of the nucleus is approximately an ellipsoid with major axis length ~20μm, which is consistent with the expected nucleus size of IMR-90 cells. (**B**) Histogram of the maximum distance between pairs of DNA-MERFISH probes for all 3,668 cells in the DNA-MERFISH data set. The maximum distance for each cell is used to normalize distances between probes in that cell. (**C**) Histogram of the number of cells in which a pair of DNA-MERFISH probes is found. The distance metric we use in the main text, i.e., the average cell-normalized Euclidean distance between two probes, corresponds to normalizing the distance between two probes in a cell by the maximum probe-probe distance in the cell in which the two probes are found, then taking the average over all cells that contain these two probes. The histogram shows that this average is taken on at least 2,800 cells and should thus enjoy a high statistical accuracy. (**D**) Histogram of average cell-normalized Euclidean distances between adhesome DNA-MERFISH probes. Intrachromosomal probe-probe distances are shown in blue, while interchromosomal probe-probe distances are shown in orange. (**E**) Scatter plot of max-min scaled Hi-C contacts and cell-normalized DNA-MERFISH Euclidean proximity (defined as (1-Euclidean distance)) for interchromosomal adhesome gene pairs (blue, $p$-value < 3e-308, see [3]) and intrachromosomal adhesome gene pairs (orange, $p$-value < 3e-308).

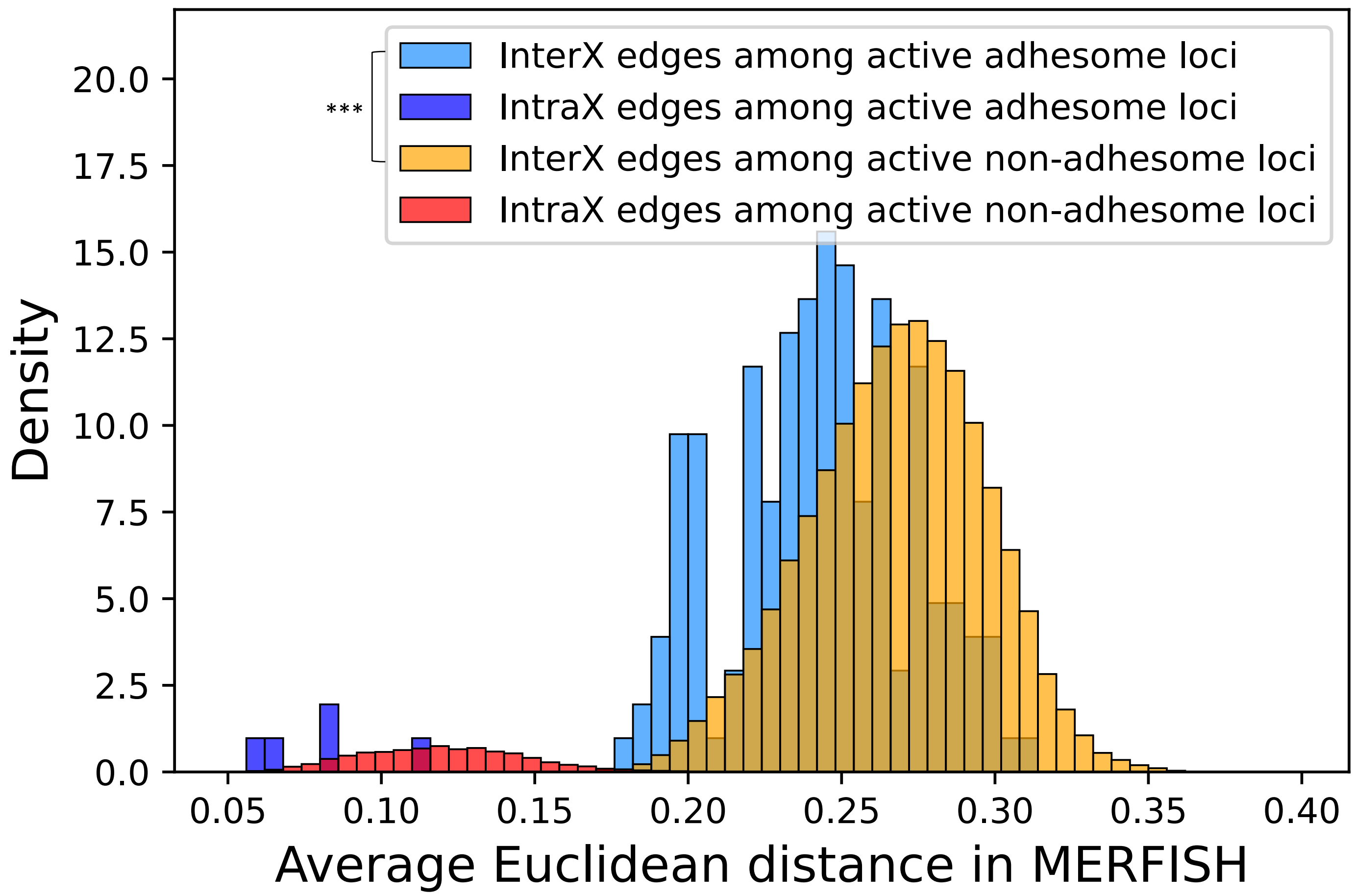


**Supplementary Figure 18**

**Supplementary Fig. 18: Extended MERFISH analysis.** Distribution of average normalized Euclidean distances between active adhesome gene probes and active non-adhesome gene probes (a MERFISH probe is associated with a gene if it overlaps with the gene). The average interchromosomal Euclidean distance between active adhesome probes is smaller than the average interchromosomal Euclidean distance among active non-adhesome probes (Wilcoxon Rank-Sum *p*-value < 1e-18).

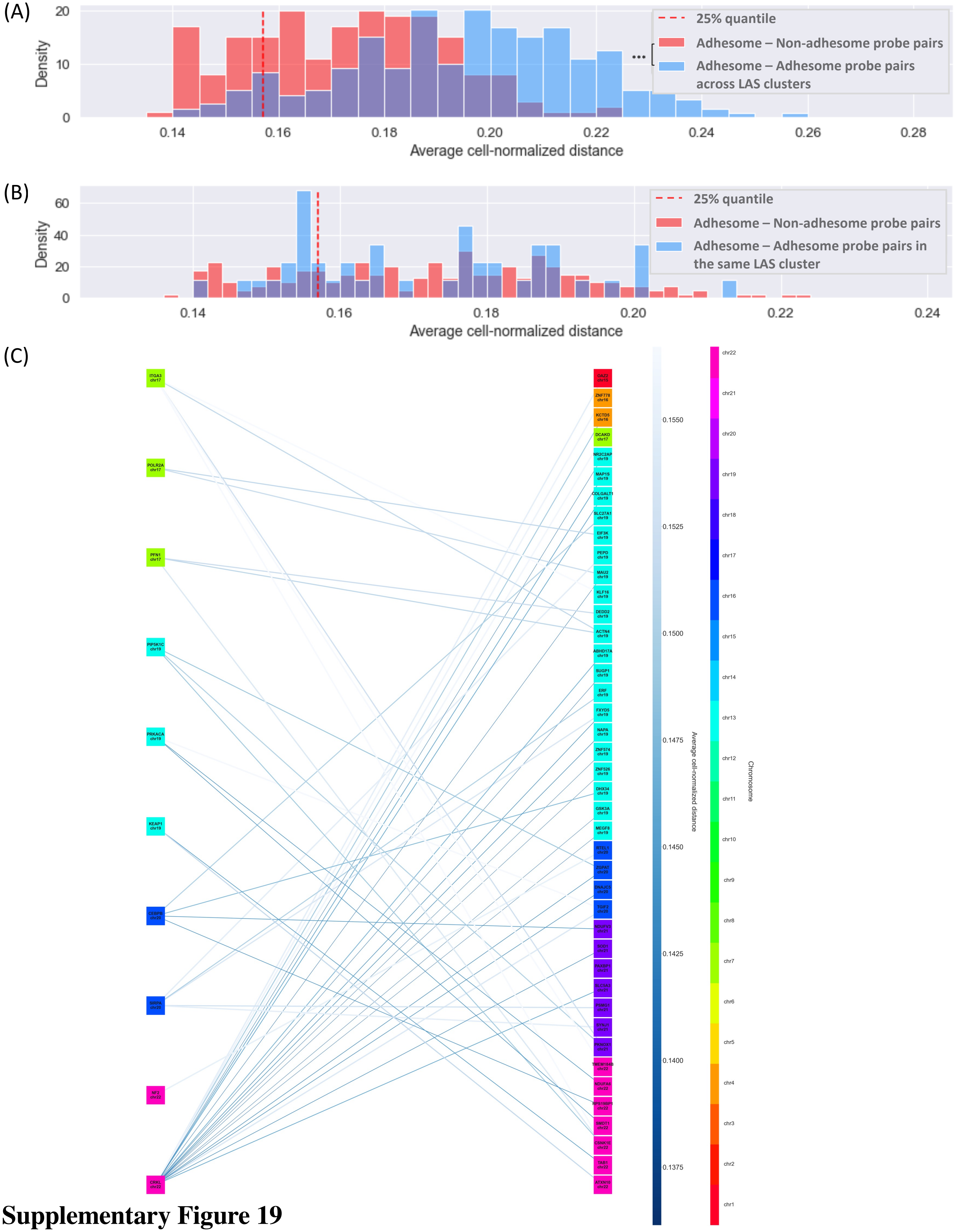


**Supplementary Figure 19**

**Supplementary Fig. 19: Bipartite network of active non-adhesome genes interacting with active adhesome genes.** (**A**) Red histogram of the average cell-normalized Euclidean distance between 135 active non-adhesome genes (61 MERFISH probes) and the 17 active adhesome genes (16 MERFISH probes) they interact with. MERFISH probes are associated with a gene if they are less than 100 kb upstream or downstream of the gene. Only gene pairs that interact in LAS and have a strong regulatory correlation (upper decile in Supplementary Fig. 10A) are considered. The red vertical dashed line corresponds to an upper threshold (first quartile) on the distance used to select edges for (C). Blue histogram of the average cell-normalized Euclidean distance between 23 active adhesome MERFISH probes across LAS regions, used for comparison with the red histogram (Wilcoxon Rank Sums test, *p*-value<2e-29). (**B**) Red histogram as in (A). Blue histogram of the average cell-normalized Euclidean distance between 13 active adhesome MERFISH probes (only LAS interactions are reported) (Wilcoxon Rank Sums test, *p*-value=0.75). (**C**) Bipartite network of active adhesome genes (10 genes, left) and active non-adhesome genes they interact with (42 genes, right). Only genes with an associated MERFISH probe (less than 100 kb upstream or downstream) are considered; more details on how this network is constructed can be found in the main text. Gene colors indicate the corresponding chromosomes. Edge colors and thicknesses reflect the physical distance between genes. Edges with an average cell-normalized distance above the first quartile threshold of (A) are not drawn.

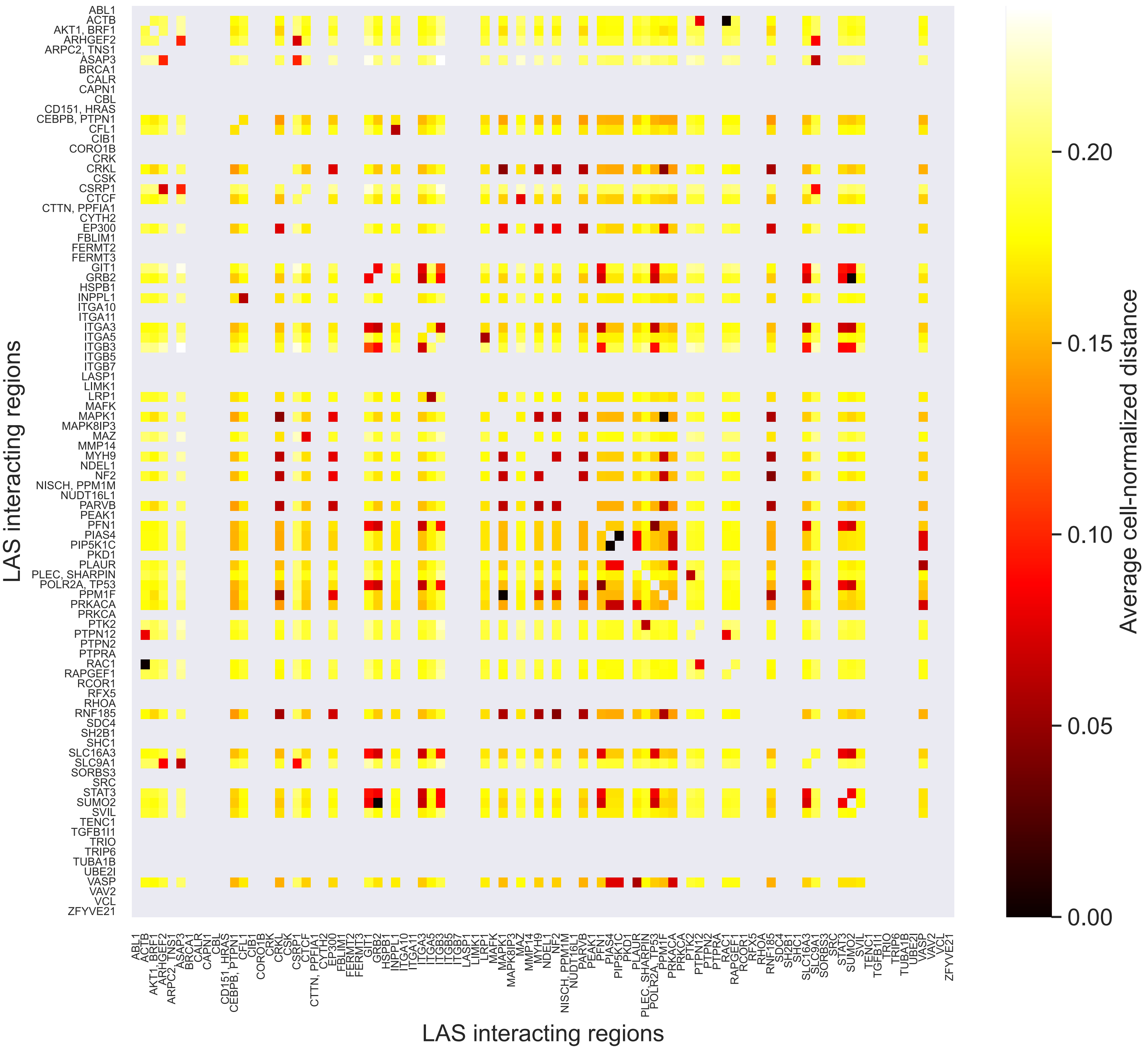


**Supplementary Figure 20**

**Supplementary Fig. 20: Relationship between Hi-C contact frequency and DNA-MERFISH Euclidean distance between adhesome loci.** Heatmap of average cell-normalized DNA-MERFISH Euclidean distance between adhesome LAS nodes. The distance between two nodes is defined by averaging the distances between all DNA-MERFISH probes associated to the first node and all DNA-MERFISH probes associated to the second node. A DNA-MERFISH probe is associated to a given adhesome LAS node if it is located at most 500kb upstream or downstream of an adhesome gene within that node. Adhesome LAS nodes that do not have an associated DNA-MERFISH probe are represented as grey rows/columns in the heatmap (missing values). Self-distances are also shown in grey. Dark entries (very low distance) typically correspond to pairs of adhesome LAS nodes on the same chromosome.

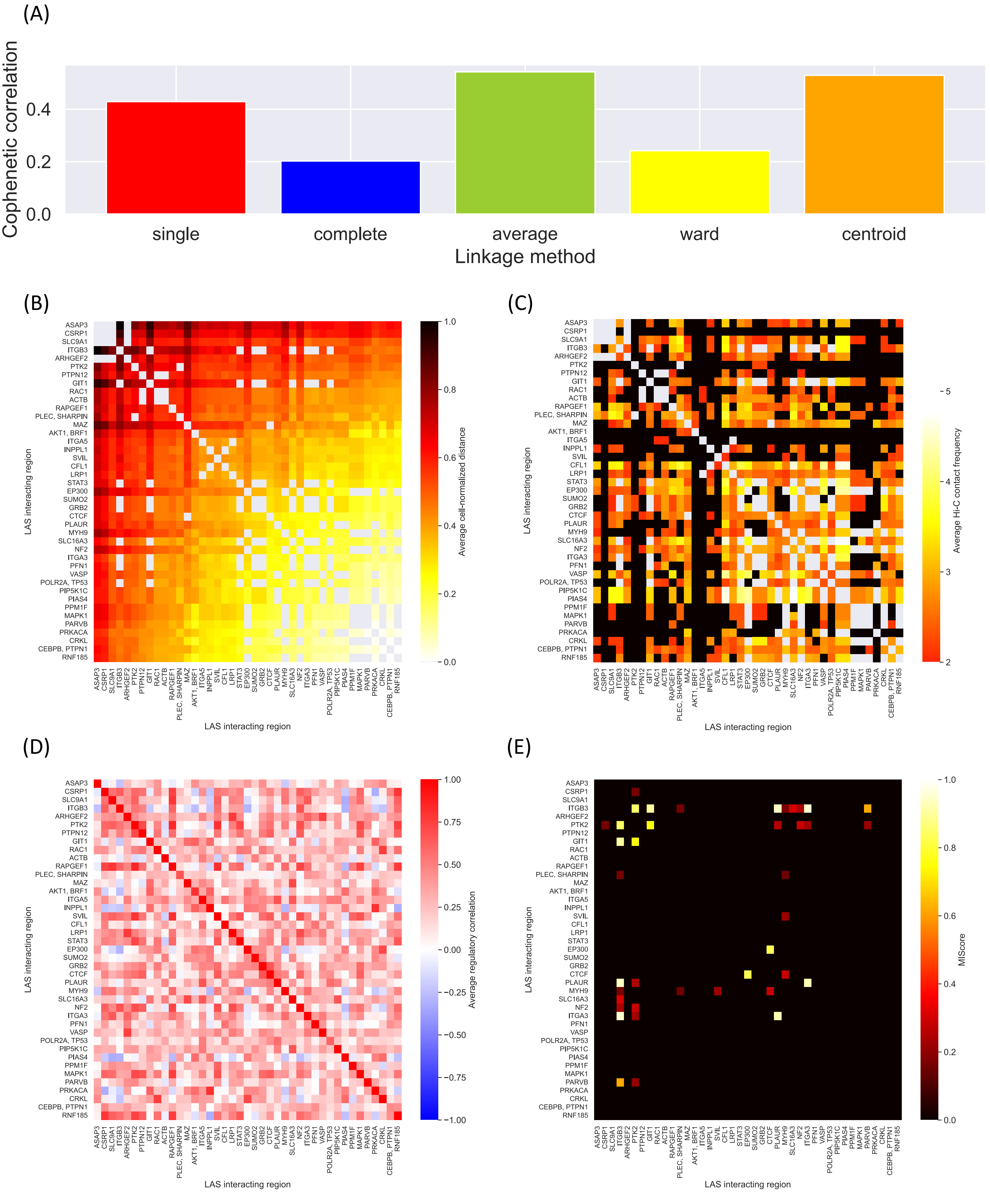


**Supplementary Figure 21**

**Supplementary Fig. 21: Joint analysis of DNA-MERFISH data, Hi-C data, regulatory data, and protein-protein interaction data identifies a cluster of highly co-localized and co-regulated adhesome genes.** (**A**) Bar plot of cophenetic correlation for different linkage functions applied to the average cell-normalized DNA-MERFISH Euclidean distance matrix between adhesome LAS nodes having at least one associated DNA-MERFISH probe. (**B**) Heatmap of average cell-normalized DNA-MERFISH Euclidean distance between adhesome LAS nodes having at least one associated DNA-MERFISH probe. Rows and columns are ordered using hierarchical clustering with single linkage. Single linkage produces a dendrogram that is reasonably faithful to the original Euclidean distances (cophenetic correlation ~0.4, see (A)) and yields an interpretable ordering of the heatmap. Entries of the heatmap that correspond to self-interactions and intrachromosomal interactions are shown in grey. (**C**) Heatmap of average transformed Hi-C contact frequency between adhesome LAS nodes having at least one associated DNA-MERFISH probe. Rows and columns are ordered as in (B). Entries of the heatmap that correspond to self-interactions and intrachromosomal interactions are shown in grey. Entries corresponding to interactions that were not identified by the LAS algorithm are shown in black. (**D**) Heatmap of average Spearman regulatory correlation between adhesome LAS nodes having at least one associated DNA-MERFISH probe. Rows and columns are ordered as in (B). (**E**) Heatmap of protein-protein physical interaction confidence score between genes in adhesome LAS nodes having at least one associated DNA-MERFISH probe. Rows and columns are ordered as in (B). Protein-protein physical interactions are retrieved from the physical STRING network. The confidence score for the interaction of two proteins is based on the number and type of sources reporting this interaction. A score of 0 indicates that two proteins are not known to physically interact.

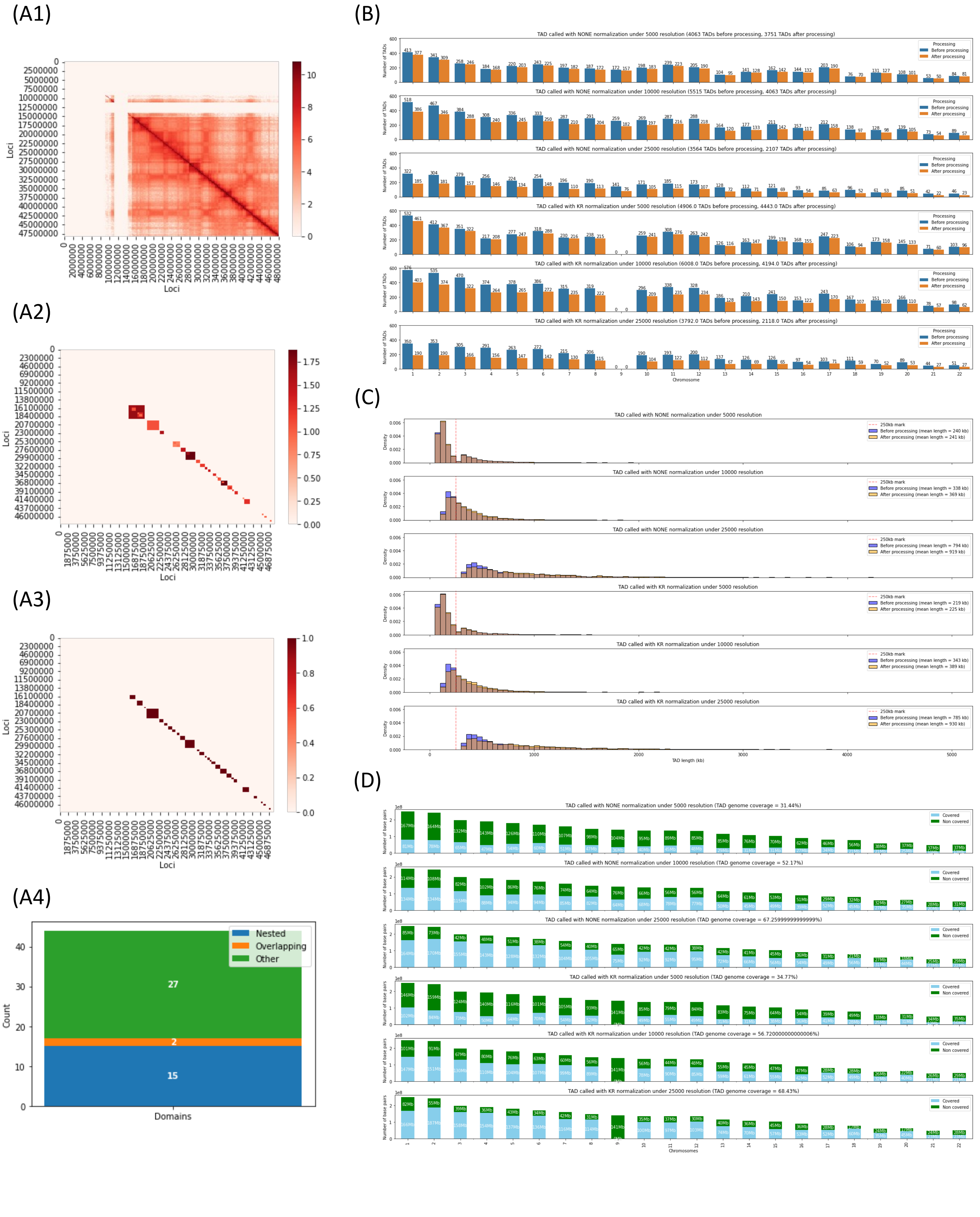


**Supplementary Figure 22**

**Supplementary Fig. 22: TAD calling.** (**A**) TAD calling pipeline shown for chromosome 21 at resolution 25 kb with Knight-Ruiz normalization. The Arrowhead software (see Methods section) is applied to the normalized intrachromosomal Hi-C contact matrix of chromosome 21 at a given resolution (A1), resulting in a list of potentially overlapping and nested TADs (A2). This TAD list is then processed to remove nested TADs and merge overlapping TADs (A3). Most TADs are disjoint, yet a minority of TADs are nested and only very few TADs are overlapping (A4). (**B**) Number of TADs called per chromosome before and after post-processing for TAD lists called by Arrowhead at different resolutions (5 kb, 10 kb, 25 kb), with and without Knight-Ruiz normalization. (**C**) Distribution of TAD length before and after post-processing for TAD lists called by Arrowhead at different resolutions (5 kb, 10 kb, 25 kb), with and without Knight-Ruiz normalization. (**D**) Number of base pairs covered per chromosome by processed TADs for TAD lists called by Arrowhead at different resolutions (5 kb, 10 kb, 25 kb), with and without Knight-Ruiz normalization.

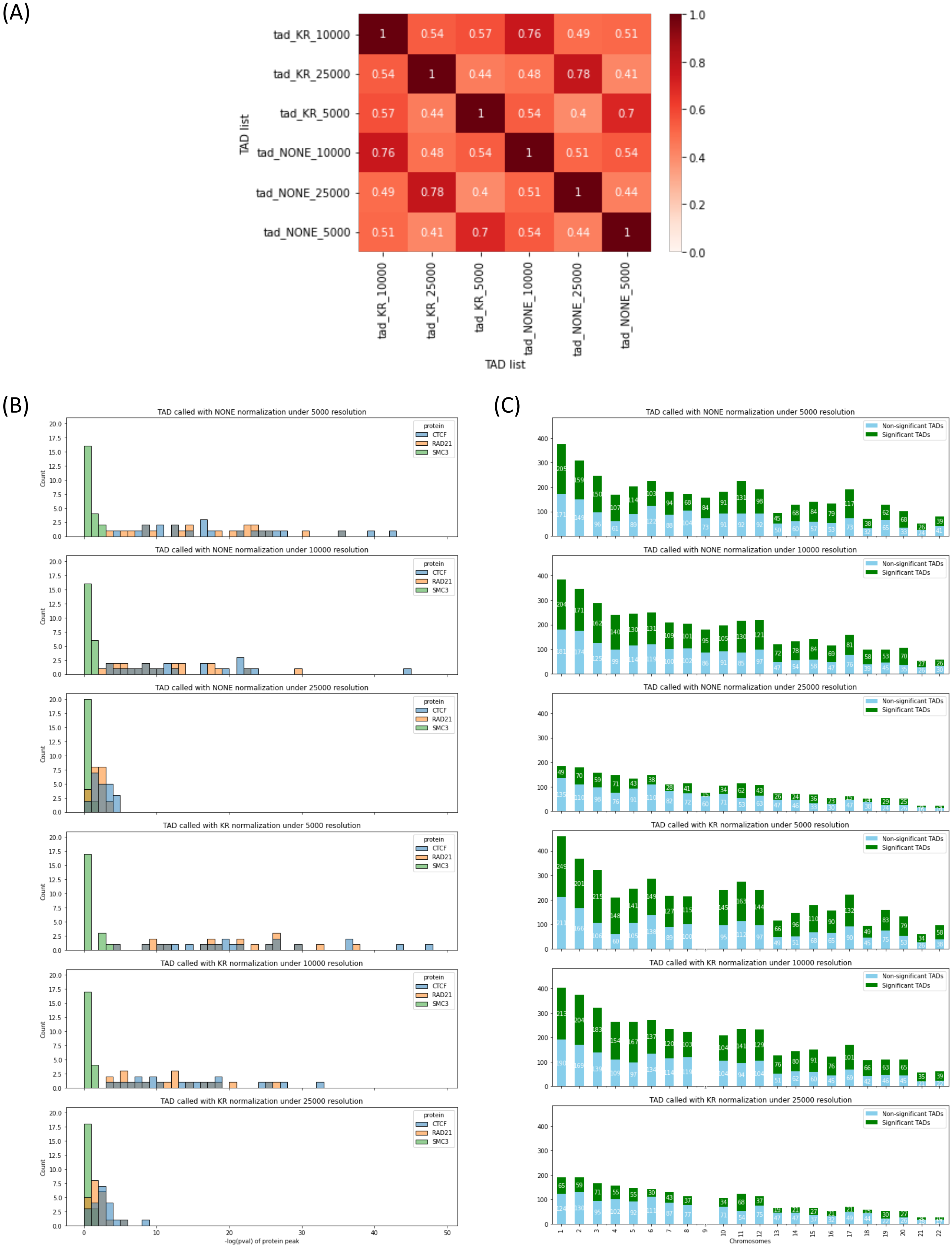


### Supplementary Figure 23

**Supplementary Fig. 23: TAD quality.** (**A**) Measure of Concordance (MoC) heatmap for TAD lists called by Arrowhead at different resolutions (5 kb, 10 kb, 25 kb), with and without Knight-Ruiz normalization. An MoC of 1 indicates that 2 TAD lists are identical, while an MoC of 0 indicates that 2 TAD lists are non-overlapping (see Methods). (**B**) Negative log-*p*-value of the structural protein profiles (see Methods) of CTCF, RAD21, and SMC3 for TAD lists called by Arrowhead at different resolutions (5 kb, 10 kb, 25 kb), with and without Knight-Ruiz normalization. (**C**) Number of TADs per chromosome significantly enriched for either transcriptional activating (H3K36me3) or repressing (H3K27me3) histone marks for TAD lists called by Arrowhead at different resolutions (5 kb, 10 kb, 25 kb), with and without Knight-Ruiz normalization. Enrichment is estimated as the log10-ratio between the ChIP-seq signals of H3K36me3 and H3K27me3 and compared with a null distribution derived from random permutations (see Methods).

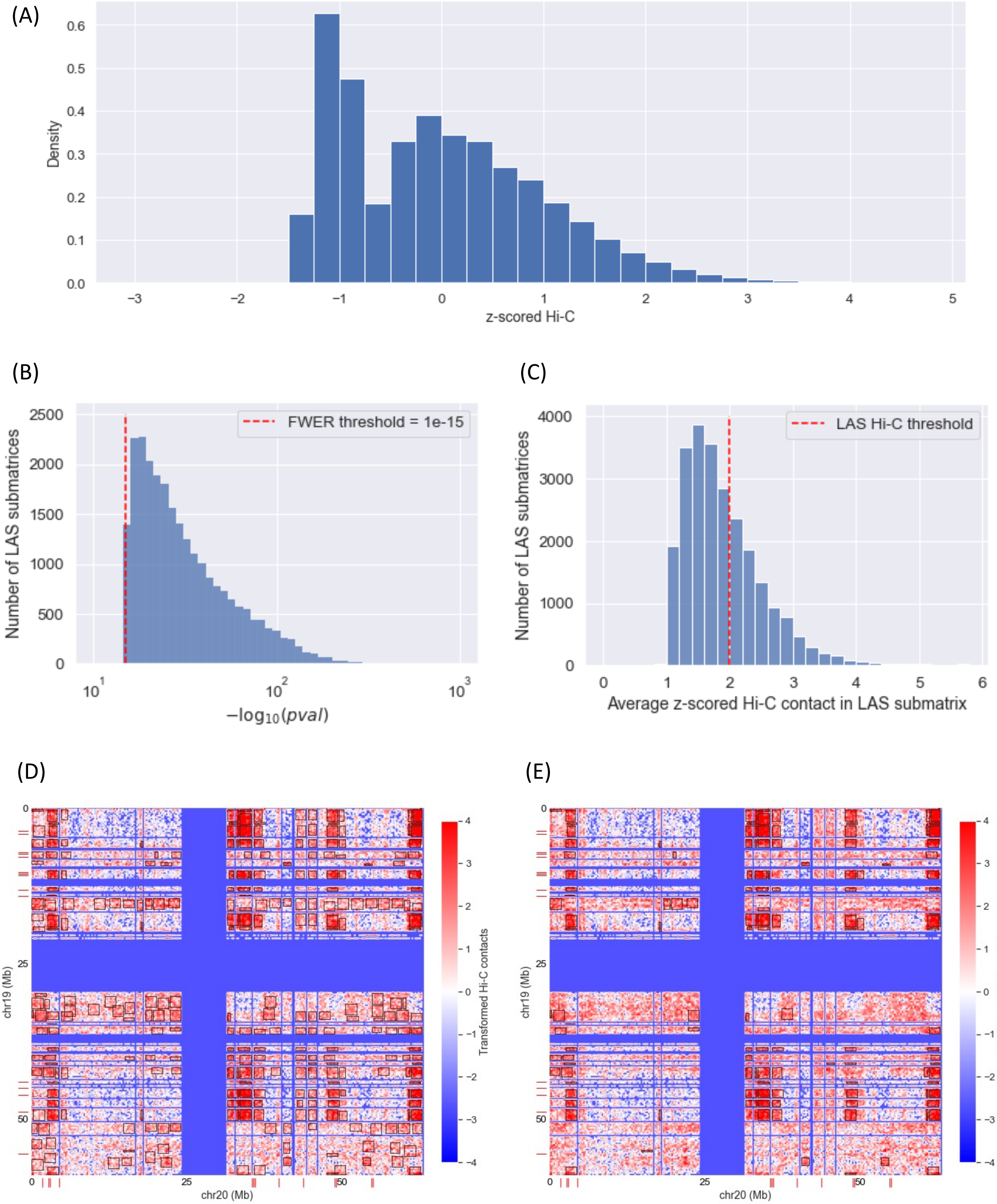


**Supplementary Figure 24**

**Supplementary Fig. 24: Clustering of transformed Hi-C contact frequencies using LAS algorithm.** (**A**) Histogram of non-zero log-transformed and centerdized Hi-C contact values in the whole genome. Centerdization was performed using the mean and standard deviation of the non-zero contact values across the whole genome. (**B**) Histogram of negative log10-transformed *p*-values of submatrices identified by the LAS algorithm [3] in all interchromosomal Hi-C contact matrices. The *p*-value threshold of 1e-15 used for signal detection is represented as a red dashed vertical line. This *p*-value threshold controls the FWER at <0.0001. (**C**) Histogram of average transformed Hi-C contact values in all LAS submatrices. The Hi-C threshold of 2 used to discard low-average submatrices is represented as a red dashed vertical line. (**D**) Heatmap of transformed Hi-C contact frequencies between genomic loci on chromosome 19 and chromosome 20. Adhesome loci are indicated by red ticks on the x-axis (for chromosome 20) and the y-axis (for chromosome 19). Significant submatrices identified by the LAS algorithm are shown in black boxes. (**E**) Same as (D), except that only significant submatrices identified by the LAS algorithm with an average Hi-C contact value higher than 2 are shown in black boxes.
